# The extensive diversity of nairoviruses reveals the complexity and high risk of tick-borne diseases in northeastern China

**DOI:** 10.1101/2025.02.08.637282

**Authors:** Ziyan Liu, Zhiwei Wei, Yu Liu, Zedong Wang

**Affiliations:** Department of Infectious Diseases, Center of Infectious Diseases and Pathogen Biology, Key Laboratory of Organ Regeneration and Transplantation of the Ministry of education, The First Hospital of Jilin University, State Key Laboratory of Zoonotic Diseases, Changchun, Jilin Province, China; International Center of Future Science, Jilin University, Changchun, Jilin Province, China

## Abstract

The increasing burden of tick-borne virus infections is becoming a global public health concern. In the current study, we identified nine tick-borne nairoviral species in northeastern China through metagenomic sequencing, which are phylogenetic classified into four groups: I) Wetland virus (WELV), II;) Songling virus (SGLV), Ji’an nairovirus (JANV), Yanbian nairo tick virus 1 (YBNTV-1), Shanxi tick virus 2 (SXTV-2), III;) Yezo virus (YEZV), and IV;) Beiji nairovirus (BJNV), Yichun nairovirus (YCNV), and Hunchun nairovirus (HCNV). These nairoviruses are influenced by both the species of ticks and their geographical distribution. Apart from the previously isolated pathogenic WELV, SGLV, YEZV, and BJNV, we successfully isolated JANV and YBNTV-1, which can replicate in Vero and HEK-293T cells. Two routes of nairovirus spread into northeastern China were evaluated. Group I and II viruses, vectored by *Haemaphysalis* sp. ticks, likely originated in Central or East China and spread via migratory birds along the Chinese coastline. Group III; and IV viruses, vectored by *Ixodes persulcatus* ticks, are believed to have originated in Europe and spread via rodents across the Eurasian continent. Our study reveals the extensive diversity and wide distribution of tick-borne nairoviruses in northeastern China, highlights the potential pathogenic risk of the newly discovered nairoviruses, and clarifies their origins and potential migration routes. The results will be of crucial significance for the tick-borne diseases control and prevention in China.

## Introduction

Emerging tick-borne viruses pose a significant global public health challenge^1^. Nairoviruses, belonging to the family *Nairoviridae*, order *Hareavirales*, are primarily maintained in arthropods and transmitted by ticks to mammals and birds^2^. Among these, Crimean-Congo hemorrhagic fever virus (CCHFV) and Nairobi sheep disease virus (NSDV) are two significant nairoviruses. CCHFV causes fatal hemorrhagic fever in humans, while NSDV leads to lethal hemorrhagic gastroenteritis in small ruminants ^3, 4^. Other nairoviruses, including Dugbe virus, Tacheng tick virus 1, Songling virus, Erve virus, Tamdy virus, Soldado virus, Ganjam virus, and Yezo virus, can cause symptoms in humans ranging from fever and headache to severe conditions like thrombocytopenia and liver injury ^5, 6, 7, 8, 9, 10, 11^.

In China, tick-borne diseases are prevalent across the nation, with higher incidence rates observed in the northeastern, northwestern, and central regions. The northeastern region known for its abundant forest resources, particularly in the Daxing’an Mountains (DXAM), Xiaoxing’an Mountains (XXAM), and Changbai Mountains (CBM), supports a large tick population. Since the first reported death from tick-borne encephalitis in the 1940s, a variety of viruses related to fever cases of tick-bitten patients, including Jingmen tick virus (JMTV), Alongshan virus (ALSV), BJNV, SGLV, YEZV, and WELV, have been identified in northeastern China ^12, 13, 14, 15, 16, 17^. Among these, SGLV, BJNV, YEZV, WELV belong to the family *Nairoviridae*. Additionally, other tick-borne nairoviruses, such as NSDV, JANV, YBNTV-1, SXTV-2, and YCNV, had also been discovered, highlighting the health threat posed by nairoviruses to humans and animals in this region ^18, 19^. However, no studies have systematically presented the spatial distribution of tick-borne nairoviruses, or explored the key factors contributing to the complexity and diversity of tick-borne nairoviruses in northeastern China.

To address this gap, we conducted a metagenomic sequencing study of major tick species from northeastern China. Our objectives were to characterize the tick nairo-virome, identify factors influencing viral diversity and distribution, and evaluate the potentially pathogenic risk of newly discovered viruses. Our findings will provide important data support for the differential diagnosis and control of tick-borne diseases in China.

## Results

### Tick collection in northeastern China

A total of 7,917 ticks from the *Ixodidae* family were collected from 20 sites located in Inner Mongolia, Liaoning, Jilin, and Heilongjiang provinces in northeastern China. The species included *Haemaphysalis concinna* (2429), *Haemaphysalis longicornis* (1329), *Ixodes persulcatus* (1855), and *Dermacentor nuttalli* (2304) (***Fig. 1a, Supplementary Table 1***). The distribution of these ticks varied by species. *I. persulcatus* (23.4%) was the most widely distributed, primarily found in mixed coniferous broad-leaved forest of 16 sites across all the four provinces. *H. longicornis* (16.8%) was mainly collected from shrub grasslands and wetlands in 10 coastal sites across Liaoning, Jilin, and Heilongjiang provinces. *H. concinna* (30.7%) and *D. nuttalli* (29.1%) were predominantly found in shrub grasslands and wetlands in 13 and 10 sites, respectively. The majority of ticks were collected from Jilin Province, (61.9%), while the fewest were from Liaoning Province (8.3%) (***Fig. 1a, Supplementary Table 1***).

**Fig. 1.**
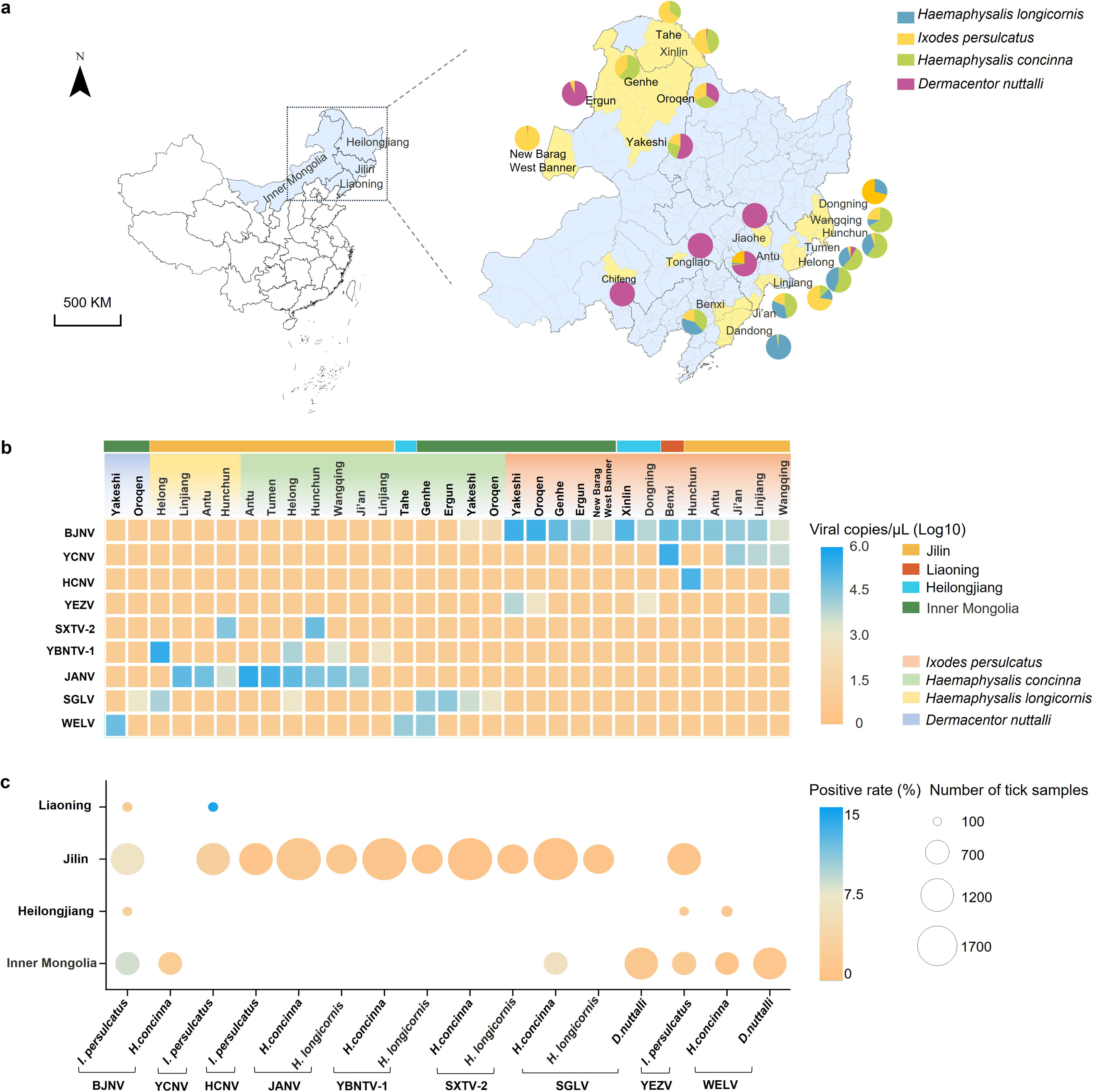
Analysis of collected ticks and tick-borne nairo-virome in northeastern China. **a)** Ticks collected from northeastern China. The collection sites of ticks were highlighted in yellow, and the number of each tick species within these regions was represented using pie charts. Four distinct tick species were distinguished by different colors: blue for *Haemaphysalis longicornis*, yellow for *Ixodes persulcatus*, green for *Haemaphysalis concinna*, and mauve for *Dermacentor nuttalli*. **b)** The abundances of nairoviruses in various tick species collected from different sites. **c)** The prevalence of identified nairoviruses in various tick species across multiple provinces. Abbreviations: WELV, Wetland virus; SGLV, Songling virus; JANV, Ji’an nairovirus; YBNTV-1, Yianbian nairo tick virus 1; SXTV-2, Shanxi tick virus 2; YEZV, Yezo virus; BJNV, Beiji nairovirus; YCNV, Yichun nairovirus; HCNV, Hunchun nairovirus.

### Analyzing of tick-borne nairo-virome

A total of 27 RNA libraries were constructed for Illumina HiSeq sequencing, generating 141.4 GB of clean data. From ∼1 billion non-rRNA reads, we *de novo* assembled ∼15.3 million contigs after removing rRNA reads. Of these, 9,480 contigs were assigned to RNA viruses, with 606 (6.4%) belonging to nairoviruses (***Supplementary Table 2***). A total of nine viral species from two genera within the family *Nairoviridae* were identified. Six species belong to the three-segmented genus *Orthonairovirus*: SGLV, JANV, YBNTV-1, SXTV-2, YEZV, and WELV. The remaining three species are from the two-segmented genus *Norwavirus*: BJNV, HCNV, and a novel species YCNV. Among these identified viral species, four of them including SGLV, YEZV, WELV, and BJNV have been confirmed associate with human febrile illness in previous studies^11, 14, 15, 16^.

### Quantification and prevalence of the identified nairoviruses

Upon confirmation by quantitative reverse transcription PCR (RT-qPCR), BJNV, YCNV, HCNV, and YEZV were predominantly identified in *I. persulcatus* ticks, whereas SGLV, JANV, YBNTV-1, SXTV-2, and WELV were mainly detected in *Haemaphysalis.* sp and *D. nuttalli* ticks (***Fig. 1b, Supplementary Table 3***). Among the genus *Orthonairovirus*, YEZV and SGLV were detected in Inner Mongolia, Heilongjiang, and Jilin provinces, with viral copies ranging from 2.3×10^2^ to 1.2×10^4^ copies/μL in ticks. SXTV-2, JANV, and YBNTV-1 were exclusive to Jilin, while WELV was only identified in Inner Mongolia and Heilongjiang sites, with viral copies from 2.2×10^2^ to 2.8×10^5^ copies/μL. Regarding the bi-segmented viruses in genus *Norwavirus*, BJNV was widely distributed in *I. persulcatus* (13 out of 16 collection sites) across the four provinces, with viral copies varying from 9.3×10^2^ to 1.7×10^5^ copies/μL. YCNV and HCNV were only detected in *I. persulcatus* in limited collection sites (4/16 and 1/16) located in Jilin and Liaoning provinces, with viral copies from 1.9×10^3^ to 1.7 ×10^5^ copies/μL. Among these norwaviruses, BJNV exhibited significantly higher prevalence in *I. persulcatus* ticks in Inner Mongolia and Jilin compared to Liaoning and Heilongjiang, while YCNV was more prevalent in Liaoning. SGLV showed significantly higher prevalence in *H. concinna* ticks in Inner Mongolia. The overall prevalence of nairoviruses ranged from 0.1% to 1.7% (***Fig. 1c, Supplementary Table 4***).

### Genomic characteristics of the identified nairoviruses

A total of 67 complete genome sequences were verified and amplified by semi-nested PCR, including 16 for BJNV, 11 each for YCNV and JANV, 9 each for SGLV and YBNTV-1, 4 each for YEZV and WELV, 2 for SXTV-2, and 1 for HCNV (***Supplementary Table 5***). Of these viruses, SGLV, JANV, YBNTV-1, SXTV-2, YEZV, and WELV, along with two other viruses tested in northeastern China (NSDV and HPTV-1)^18, 20^, belong to the three-segmented genus *Orthonairovirus*, which consists of three RNA segments: a small (S) segment encoding 483–503 aa nucleocapsid protein (NP), a medium (M) segment encoding 1,338–1,628 aa glycoprotein precursor (GPC), and a large (L) segment encoding 3,922–3,969 aa RNA-dependent RNA polymerase (RdRp). BJNV, HCNV, and YCNV belong to the bi-segmented genus *Norwavirus*, which is comprised of two RNA segments: a S segment encoding 545–556 aa NP and an L segment encoding 4,811–4,812 aa RdRp (***Fig. 2, Supplementary Fig. 1 and 2***).

**Fig. 2.**
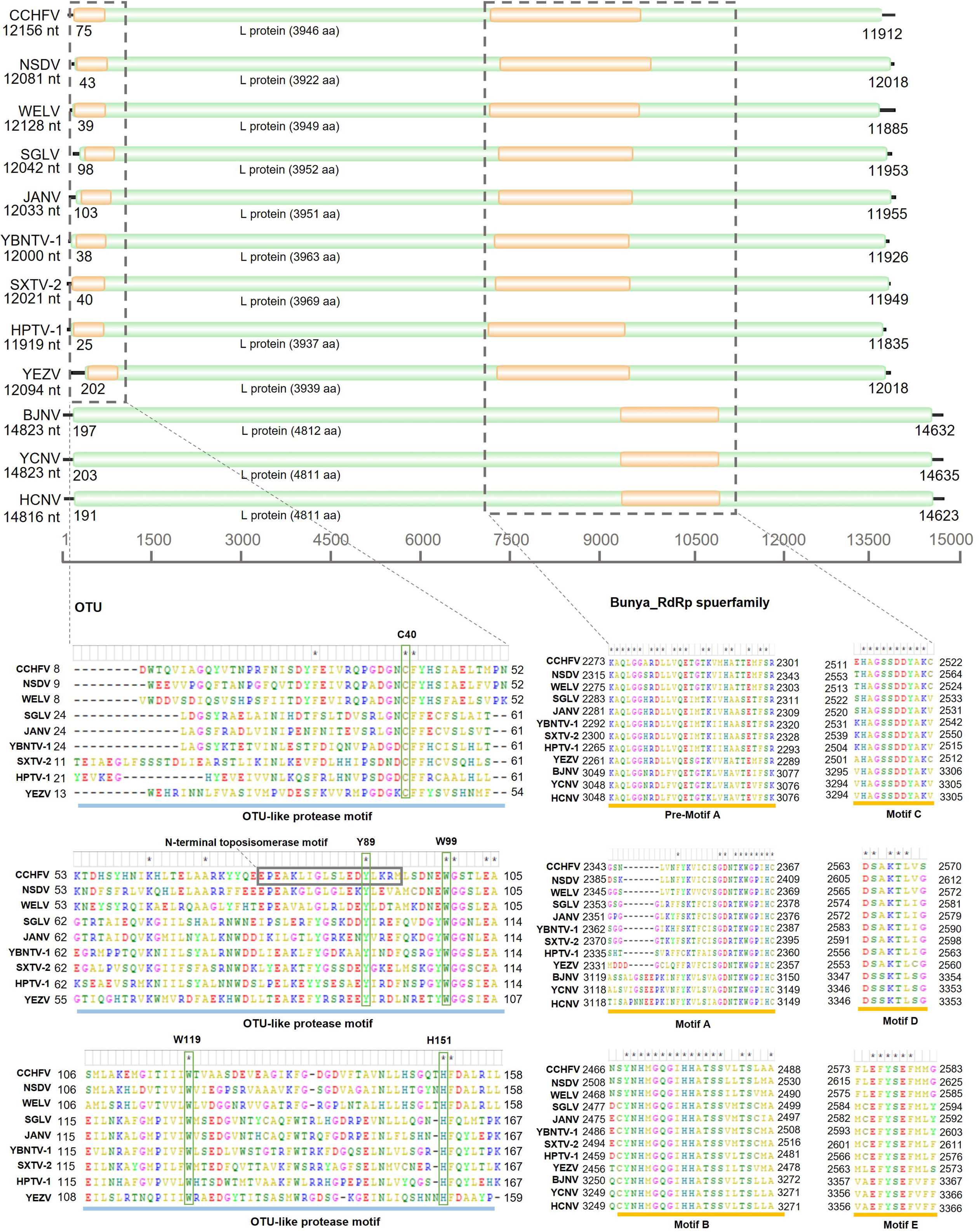
Genomic characterization of the large (L) segment of nairoviruses identified in northeastern China. The predicted open reading frame (ORF) of L segment that encodes RNA-dependent RNA polymerase (RdRp) is shaded in green. CCHFV were used as a reference viral species. Abbreviations: OUT, Ovarian tumor domain.

We found that all the three-segmented orthonairoviruses encode a deubiquitinase (DUB) of the ovarian tumor-associated protease (OTUs) family, which plays important roles in the ubiquitin- and ISG15-dependent innate immune responses ^21, 22^. This DUB was not identified in any of the bi-segmented norwaviruses (***Fig. 2)***. Moreover, several highly conserved OTU-characteristic residues (C40, Y89, W99, W119, and H151) identified in previous study were also observed among these orthonairoviruses^23^. Among these conserved residues, Y89 is the key residue in the topoisomerase I active site, which has been identified in most nairoviruses causing diseases in humans or other mammals, but not in those infecting only arthropods or establishing subclinical infections in vertebrates^23^. Conserved aa positions were found in the core region of the RdRp of all the nairoviruses, including pre-motif A and motifs A to E (***Fig. 2***).

### Phylogenetic and homology analyses

Phylogenetic analysis classified these nairoviruses into four groups: three three-segmented orthonairovirus groups (I-III) and one two-segmented norwavirus group (IV). Group I (WELV and NSDV) clustered with Tofla virus and CCHFV. Group II (SGLV, JANV, YBNTV-1, SXTV-2, and HPTV-1), grouped with Henan tick virus, Wenzhou tick virus, Tacheng tick virus 1, and Tamdy virus that were found in central and northwestern China. Group III (YEZV) grouped with Sulina virus. Group IV (BJNV, YCNV, and HCNV) clustered with other bi-segmented norwaviruses identified in Europe, including Pustyn virus, Norway nairovirus, and Grotenhout virus (***Fig. 3a, Supplementary Fig. 3***).

**Fig. 3.**
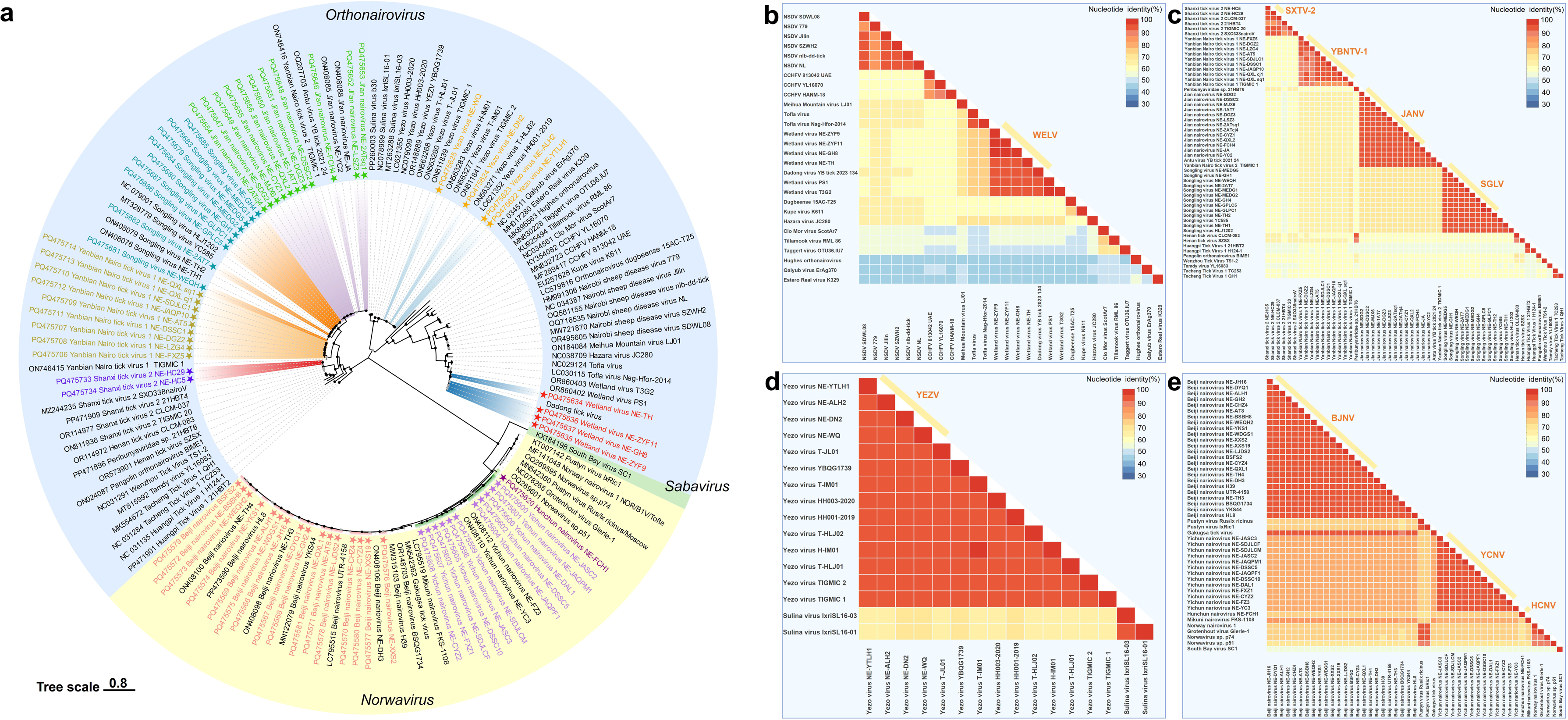
Phylogenetic and homology analysis of the identified tick-borne nairoviruses in northeastern China. **a)** Phylogenetic tree based on the RNA-dependent RNA polymerases (RdRp) amino acid sequences of nairoviruses. The tree was constructed by the maximum likelihood method using the best-fit model LG+I+G4. The newly identified nairoviruses in the present study were labeled by symbol stars. **b-d)** Nucleotide sequence identities of the complete Segment L of the four nairovirus groups (I-IV).

Sequence comparison revealed that WELV exhibits a close phylogenetic relationship with CCHFV and NSDV, with nucleotide (nt) sequence identities ranging from 40.2% to 69% and amino acid (aa) sequence identities ranging from 36.3% to 70% (***Fig. 3b, Extended Table 1***–***3***). Similarly, JANV, YBNTV-1, SXTV-2, and HPTV-1 are closely related to SGLV, with nt sequence identities ranging from 45.5% to 74.2% and aa sequence identities ranging from 44.8% to 85.2% (***Fig. 3c, Extended Table 4***–***6***). YEZV showed a close relationship with Sulina virus, with nt sequence identities ranging from 55.3% to 70.2% and aa sequence identities ranging from 52.3% to 82.1% (***Fig. 3d, Extended Table 7***–***9***). HCNV and YCNV showed close relationship with BJNV, with nt sequence identities ranging from 69.6% to 82.4% and aa sequence identities ranging from 84% to 88.6% (***Fig. 3e, Extended Table 10-11***).

### Isolation of potentially pathogenic nairoviruses

Among the five nairoviruses with potential pathogenic risk to humans—JANV, YBNTV-1, SXTV-2, YCNV, and HCNV—only JANV and YBNTV-1 were successfully isolated in Vero cells. Transmission electron microscopy revealed spherical virions with diameters ranging from 100 to 120 nm in the cytoplasm and supernatant of Vero cells, which is consistent with the morphology of the *Nairoviridae* family (***Fig. 4a-b, Supplementary Fig. 4***). Severe typical cytopathic effect (CPE) was noted in JANV-infected cells by the third passage, whereas mild CPE was observed in YBNTV-1-infected cells, which were subsequently confirmed by immunofluorescence assay (***Fig. 4c-d, Supplementary Fig. 4***). One-step growth curve assay in various human cell lines (HuH-7, 293T, WISH, THP-1) showed that 293T cells supported JANV replication to titers comparable to those in Vero cells, with low viral RNA replication observed in HuH-7 cells. No viral RNA replication was detected in WISH and THP-1 cells over a period of 15 days (***Fig. 4e***). For YBNTV-1, only 293T exhibited minimal viral RNA replication, while no viral RNA replication was observed in other human cell lines (***Fig. 4f***).

**Fig. 4.**
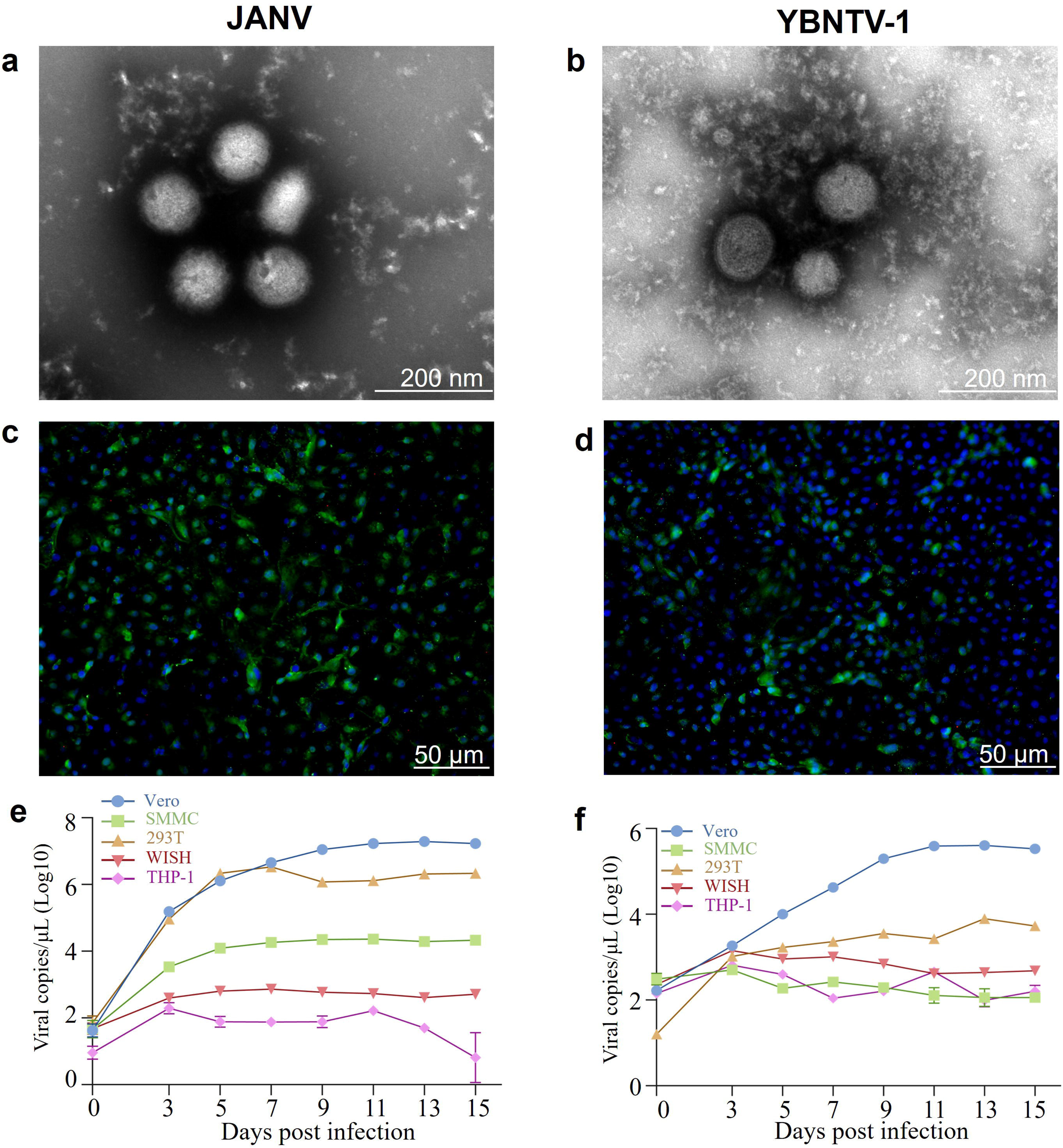
Isolation of potentially pathogenic tick-borne Ji’an nairovirus (JANV) and Yanbian nairo tick virus 1 (YBNTV-1). **a-b)** Spherical virions of JANV (a) and YBNTV-1 (b) observed in the supernatant of Vero cells using transmission electron microscopy. **c-d)** Detection of JANV (c) and YBNTV-1 (d) viral antigens in Vero cells using immunofluorescence assay. Viral antigens to nucleocapsid protein are labeled in green, and cell nuclei are stained blue. **e-f)** Growth of JANV (e) and YBNTV-1 (f) in different human cell lines. Viruses were inoculated at a multiplicity of infection (MOI) of 0.001 and cultured for 14 days. Viral RNA copy numbers at each time point were quantified using RT-qPCR.

### Mapping of the tick-borne nairoviruses in northeastern China

To map the distribution of tick-borne nairoviruses in northeastern China, in addition to the viruses found in this study, we also incorporated two additional viral species (NSDV and HPTV-1) identified in previous studies^18, 20^. Of the three-segmented viruses within the genera *Orthonairovirus*, SGLV and YEZV are widely distributed in Daxing’an Mountains (DXAM), Xiaoxing’an Mountains (XXAM), and Changbai Mountains (CBM). JANV emerges in XXAM and CBM, while YBNTV-1, SXTV-2, and HPTV-1 are merely discovered in CBM. With respect to viruses that are closely associated with CCHFV, WELV is widely distributed across DXAM and CBM, while NSDV is exclusively found in CBM. Regarding the bi-segmented nairoviruses, BJNV is extensively found across DXAM, XXAM and CBM. In contrast, YCNV occurs in XXAM and CBM, and HCNV is exclusive to CBM situated in Jilin Province (***Fig. 5, Supplementary Table 6***).

**Fig. 5.**
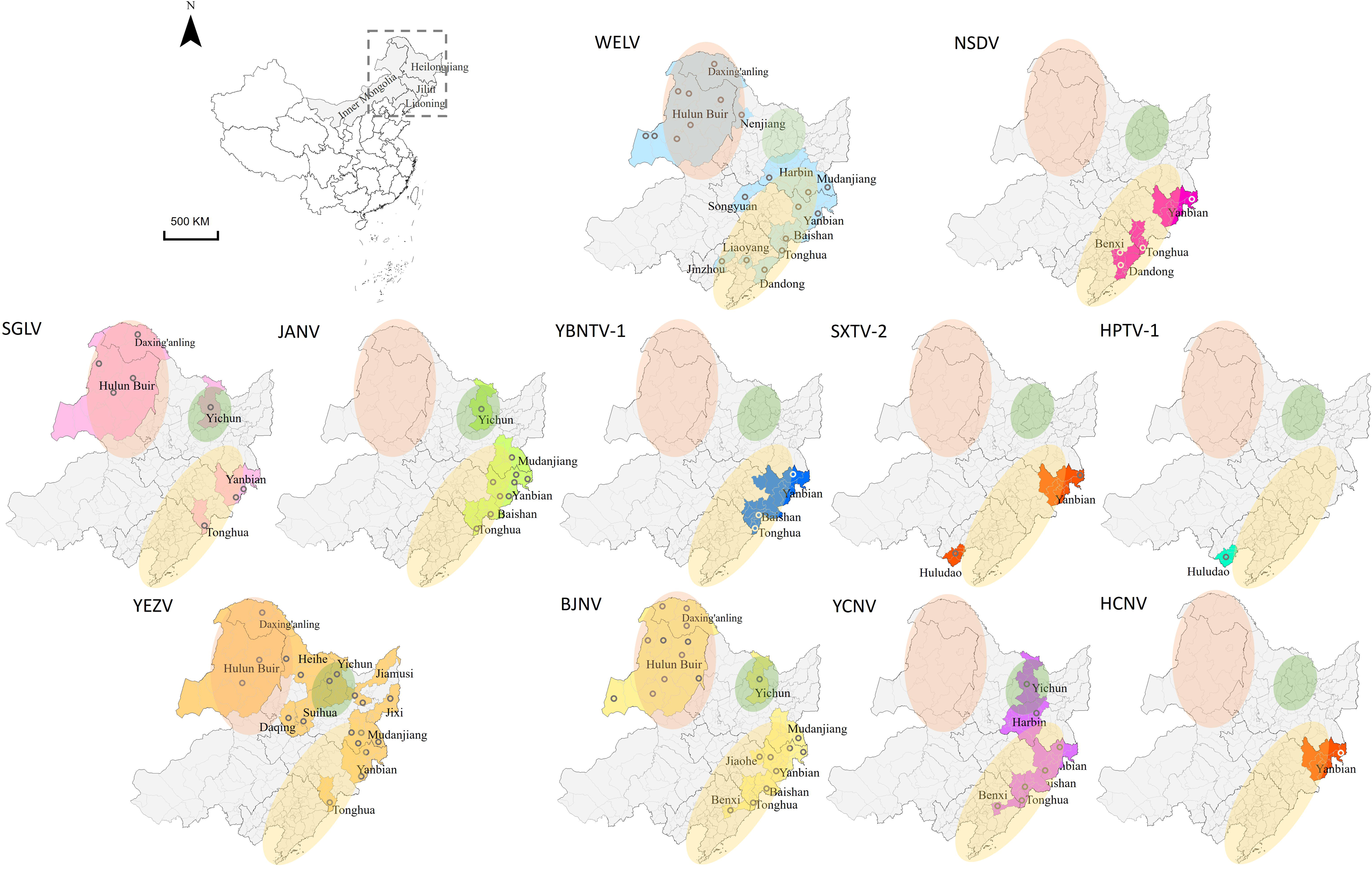
Geographic distribution of tick-borne nairoviruses in northeastern China. The shaded areas in different colors indicate the distribution of nairoviruses. Hollow circles denote the specific locations where the viruses were identified. The Daxing’an Mountain (DXAM), Xiaoxing’an Mountain (XXAM), and Changbai Mountain (CBM) regions are represented by orange, green, and yellow ellipses, respectively.

### Phylodynamic analysis of the dientified nairoviruses

To elucidate the diversity and origin of tick-borne nairoviruses in northeastern China, Bayesian analysis was conducted to estimate the time of the most recent common ancestor (tMRCA) of the four phylogenetically classified viral groups, based on nucleotide sequences of the S segments (***Supplementary Table 7***). The analyses disclosed that the tick-borne nairoviruses in northeastern China predominantly originated from Central or East China, or Europe, and were mainly disseminated through two distinct routes.

For the first route, viruses in Group I (NSDV and WELV) and Group II (SGLV, JANV, YBNTV-1, SXTV-2, and HPTV-1), which are mainly vectored by *Haemaphysalis sp*., are associated with tick-borne nairoviruses discovered in the Central or East China and spread to the northeast of China along the coastline. WELV was estimated to have a median tMRCA of 1,974 (95% HPD = 1,953 to 2,015) with Tofla virus identified in Japan, while for the NSDV strains in China shared a median tMRCA of 1,975 (95% HPD = 1,957 to 2,014) with other strains (***Fig. 6a***). SGLV, JANV, and YBNTV-1 were estimated to have a median tMRCA of 1,953 (95% HPD = 1,857 to 2,002) with Henan tick virus, SXTV-2, and HPTV-1 identified in Central and East China. The tMRCA for YBNTV-1 were estimated to be 1,989 (95% HPD = 1,930 to 2,016), and for SGLV and JANV to be 2,004 (95% HPD = 1,972 to 2,018), for HPTV-1 to be 1,975 (95% HPD = 1,910 to 2,007), and for SXTV-2 to be 1,986 (95% HPD = 1,930 to 2,011) (***Fig. 6b***). The migration pathways were illustrated in a regional map of China, South Korea, and Japan, showing routes from Shandong to CBM, Hubei to CBM, and Japan to DXAM. Moreover, migration pathways between mountains in northeastern China were also identified, as the viruses migrated from CBM to DXAM and XXAM, and DXAM to CBM (***Fig. 7a, 7b***).

**Fig. 6.**
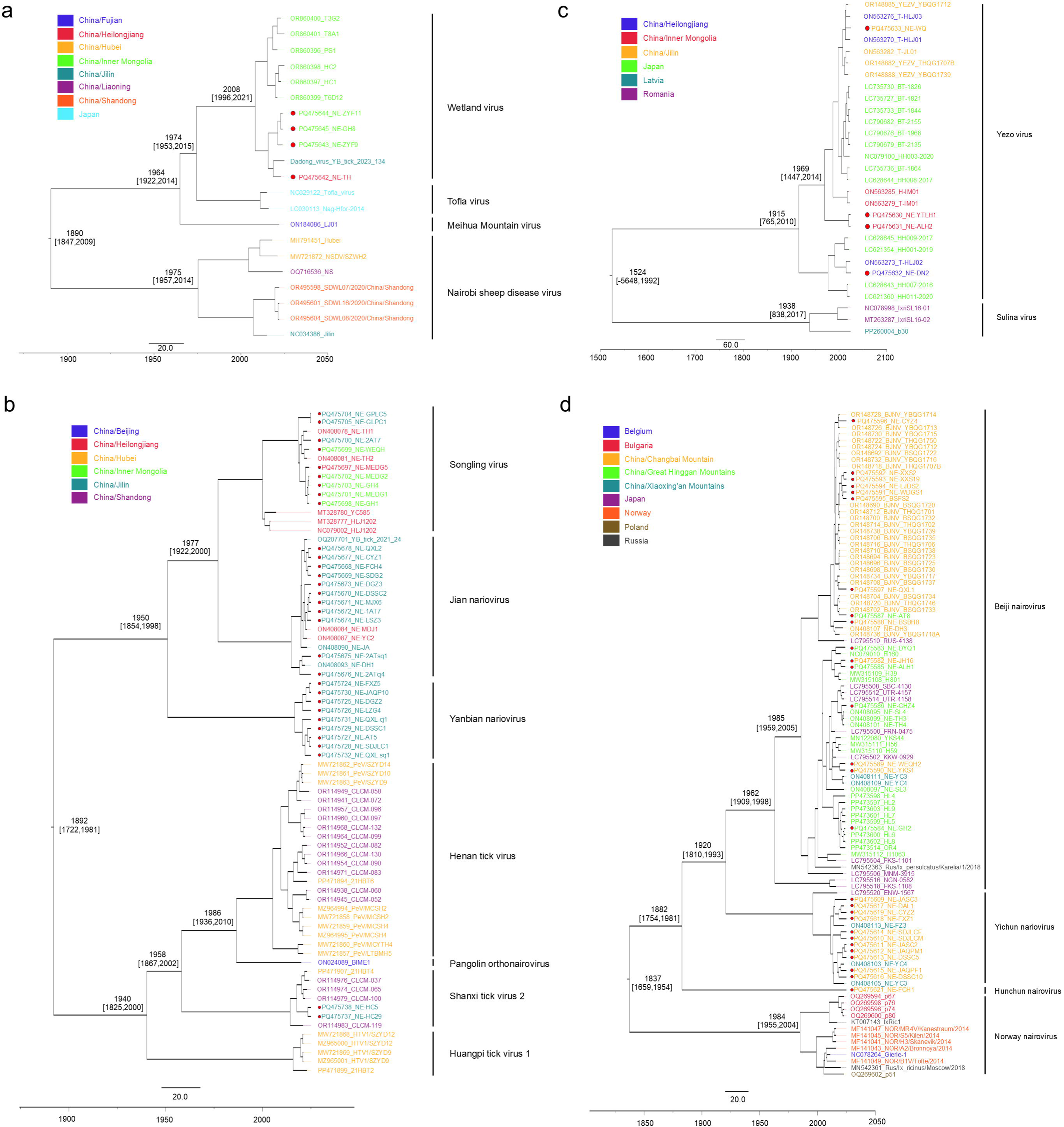
Time-scaled Bayesian MCC phylogenetic trees based on the nucleotide sequences of the S segments of nairoviruses in northeastern China. MCC phylogenetic tree nodes of virus groups I (**a**), II (**b**), III (**c**), IV (**d**) were annotated with estimated median dates of the time to most recent common ancestor (tMRCA) and 95% Highest Posterior Density (HPD). The horizontal axis indicates time in years.

**Fig. 7.**
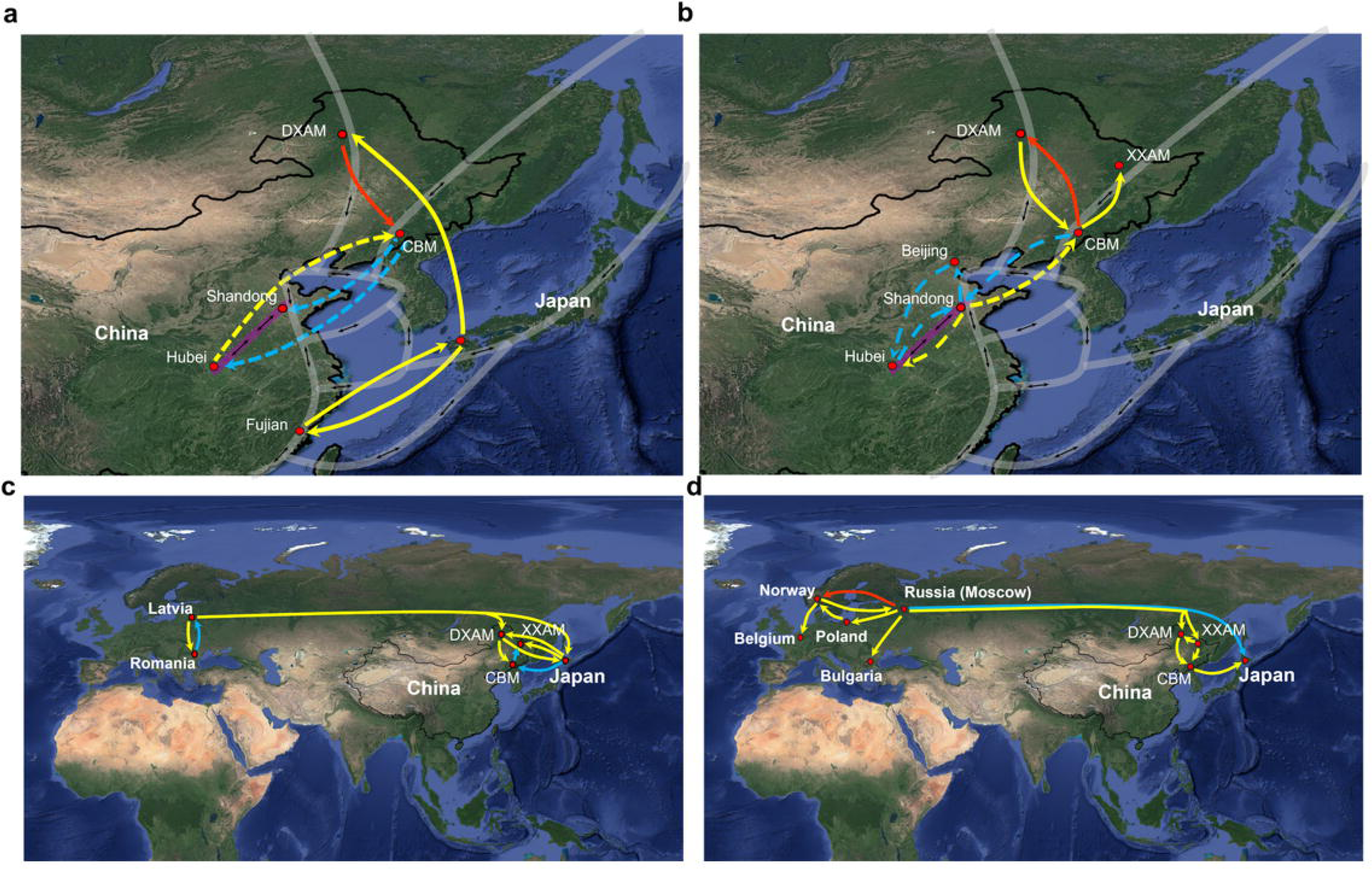
Predicted origin and migration routes of nairoviruses identified in northeastern China. Significant epidemiological pathways from one location to another for virus groups I (**a**), II (**b**), III (**c**), IV (**d**) were represented on the maps based on Bayes Factor (BF) > 3 and Posterior Probability (PP) > 0.5. Red arrows indicate very strongly supported rates with BF ≥ 100; blue arrows indicate strongly supported rates with 10 ≤ BF ≤ 100; and yellow arrows indicate supported rates with 3 ≤ BF ≤ 10. The transparent white lines with two-way arrows indicate the East Asia/Australasia flyway of migratory birds, while the transparent purple line with two-way arrows specifically denotes the migratory bird route in central China, which is integrated into the East Asia/Australasia flyway. The dashed arrows in (a) indicate the migration routes of NSDV, while the solid arrows denote the migration routes of WELV. The dashed arrows in (b) indicate the migration routes of SXTV-2 and HPTV-1, while the solid arrows denote the migration routes of SGLV, JANV, and YBNTV-1.

For another route, viruses in Group III (YEZV) and Group IV (BJNV, YCNV and HCNV), which are mainly vectored by *I. persulcatus* ticks and have close relationships with viruses found in Europe, are originated from Europe and might spread to northeastern China along the Eurasian continental plate.

YEZV was estimated to have a median tMRCA of 1524 (95% HPD = -5648 to 1992) with Sulina virus identified in Europe (Fig. 6c). BJNV, YCNV, and HCNV were estimated to have a median tMRCA of 1947 (95% HPD = 1648 to 1957) with other bi-segmented viruses such as Pustyn virus, Norway nairovirus, and Grotenhout virus identified in Europe.

YEZV was estimated to have a median tMRCA of 1,524 (95% HPD = -5,648 to 1,992) with Sulina virus identified in Europe (***Fig. 6c***). BJNV, YCNV, and HCNV were estimated to have a median tMRCA of 1,947 (95% HPD = 1,648 to 1,957) with other bi-segmented viruses such as Pustyn virus, Norway nairovirus, and Grotenhout virus identified in Europe. The tMRCAs were estimated to be 1,983 (95% HPD = 1,745 to 1,972), 1,918 (95% HPD = 1,810 to 1,985), and 1,965 (95% HPD = 1,914 to 1,999) for HCNV, YCNV, and BJNV, respectively (***Fig. 6d***). The migration pathways were illustrated in a regional map of Eurasia, showing routes from Russia (Moscow) to DXAM and XXAM, Latvia to DXAM, and Japan to DXAM, XXAM, and CBM (***Fig. 7c, 7d***).

## Discussion

Our study disclosed the extensive diversity and potential pathogenic risk of tick-borne nairoviruses in northeastern China. To date, such a considerable number of tick-borne nairoviral species (11 species) have not been identified in other areas of the country. Our previous investigation has revealed that the composition of tick-borne viruses is influenced by both the species of ticks and their geographical distribution^19^. This finding has been reaffirmed in tick-borne nairoviruses in the current study. Nairoviruses infecting *I. persulcatus* appear to face challenges in crossing species to infect *Haemaphysalis sp.*, but there is a certain possibility of infecting *D. nuttalli* (such as BJNV, ***Fig. 1b***). In contrast, viruses infecting *Haemaphysalis sp.* and *D. nuttalli* can achieve cross-species infection more readily, exemplified by SGLV and WELV. It is also notable that BJNV, YEZV, and SGLV were extensively distributed in DXAM, XXAM, and CBM of northeastern China, while WELV were detected in both DXAM and CBM. JANV and YCNV were observed in CBM as well as XXAM. NSDV, SXTV-2, YBNTV-1, and HCNV were only detected in CBM (***Fig. 5***).

Our study provides insights into the potential pathogenic risks posed by these newly discovered nairoviruses to humans. For instance, JANV, YBNTV-1, and SXTV-2 showed close relationship with pathogenic SGLV with aa sequence identities of 49.8 to 85.2%. Similarly, HCNV and YCNV exhibited close relationship with pathogenic BJNV with aa sequence identities of 84 to 88.6% (***Fig. 3, Extended Table 4–6, 10–11***). The analysis of key pathogenic sites of nairoviruses further highlights the potential pathogenic risk posed by these viruses (***Fig. 2***). Studies have posited that the CCHFV OTU domain is a critical virulence determinant. The observed structural and phylogenetic differences in the nairovirus OTUs between pathogenic and non-pathogenic nairoviruses are intriguing. The majority of nairoviruses pathogenic to humans (CCHFV, DUGV, ERVEV, ISKV, KASV, and NSDV) or other mammals (NSDV and GANV) contain an identifiable topoisomerase I active site motif, whereas most nairoviruses without that domain are only known to infect arthropods or to establish subclinical infections in vertebrates^23^. Additionally, the one-step growth curve assay revealed that JANV and YBNTV-1 can replicate in human 293T and HuH-7 cell lines, indicating the potential risk of these two viruses to human health.

Phylodynamic analyses identified two migration routes contributing to the extensive diversity and disparities in the geographical distribution of these tick-borne nairoviruses in northeastern China. Studies have affirmed that migratory birds play a crucial role in the dissemination of tick-borne viruses, as the ticks can adhere to the birds to achieve long-distance movement^24, 25^. The East Asia/Australasia flyway, stretching from Arctic Russia to the southern boundaries of Australia and New Zealand, encompasses extensive areas of eastern Asia, including the coastal regions of China, South Korea, and Japan^26^. Poyang Lake, the largest freshwater body located in central China, is also part of the East Asia/Australasia flyway^27^. These birds commute between breeding grounds and wintering areas along the flyway, taking rests at numerous stopover sites located along the coastal areas.

It has been speculated that Severe fever with thrombocytopenia syndrome virus (SFTSV), an emerging tick-borne phenuivirus with high mortality rates mainly vectored by *H. longicornis* in China, Japan, and Korea, can be disseminated by migratory birds via the East Asia/Australasia flyway to northeastern China^28, 29^. In this study, SGLV, JANV, YBNTV-1, SXTV-2, and HPTV-1 in Group I, and NSDV and WELV in Group II, which are mainly vectored by *Haemaphysalis* sp., were analyzed. These viruses might be originated from Central or East China and spread to northeastern China via migratory birds, similar to the pathway of SFTSV.

It is known that tick-borne encephalitis virus (TBEV) mainly relies on *I. persulcatus* (distributed in northeastern Asia) and *I. ricinus* (distributed in Europe) as main vectors. TBEV is grouped into three subtypes, namely Far Eastern (FE)-TBEV, Siberian (Sib)-TBEV, and European (Eu)-TBEV, of which, FE-TBEV strains were mainly distributed in northeastern Asia and have been traced back to central Europe. Rodents play a significant role in the dissemination of tick populations and the transmission of TBEV. As ticks not only utilize rodents as the principal blood-feeding host but also can adhere to their body for prolonged periods, this facilitates population diffusion. Global climate change may further expand the invasive range of rodents (particularly *Apodemus agrarius*) from Central Europe to East Asia^30^. The period ranging from the 1300s to the end of the 1900s, known as the Little Ice Age^31^, led to a decrease in predator numbers, including poikilotherms, resulting in a surge in rodent populations^32, 33^. The extensive invasion of rodents may have contributed to the dissemination of Eu-TBEV from central Russia and its evolution into FE-TBEV in Northeast Asia^34^.

In our study, YEZV in Group III, and BJNV, YCNV, and HCNV in Group IV, which are mainly vectored by *I. persulcatus* ticks, have close relationships with Sulina virus, and other bi-segmented viruses (Pustyn virus, Norway nairovirus, and Grotenhout virus) found in Europe and vectored by *I. ricinus* ticks. The tMRCAs of these viruses fall within the temporal range of the Little Ice Age, suggesting that the dissemination of rodent populations might have played a significant role in the spread and evolution of these viruses. They might have spread with the invasion of rodents to northeastern China via the same pathway as TBEV.

There are some limitations in this study. Firstly, among the nine nairoviruses identified in the study, except for the pathogenic BJNV, YEZV, SGLV, and WELV which have been successfully isolated in previous studies^11, 14, 15, 17^, only JANV and YBNTV-1 were successfully isolated. It is crucial to emphasize that the unsuccessful isolation of some viruses does not imply its inability to infect humans or animals and induce disease. Several factors might impede the successful isolation of the viruses, including variations in pathogenicity among different viral strains, the lack of an appropriate cell line, insufficient viral titers in samples, or the limited number of positive tick samples. Secondly, we did not perform any epidemiological investigations of the novel nairoviruses in either humans or animals, highlighting that our understanding of the mechanisms through which these viruses infect vertebrates and elicit disease remains limited. Lastly, owing to the extremely limited number of tick-borne nairoviruses sequences, such as in Europe and the Russian Far East, our phylodynamic analysis results are inevitably biased and lack evidence for some key nodes. Hence, more genomes of nairoviruses need to be discovered in related countries to better understand the spread and evolution of tick-borne nairoviruses in Eurasia.

## Conclusion

In summary, our study revealed the extensive diversity and wide distribution of tick-borne nairoviruses in northeastern China, clarified the reasons for the viral diversity, and inferred the potential migration routes by which viruses were introduced into the area. The results emphasize the potential pathogenic risk of the newly discovered nairoviruses and will help better understand the evolution mechanisms of the viruses, which is crucial for developing public-health measures and strategies for tick-borne disease control and prevention in China.

## Methods

### Tick collection

From April 2022 to July 2023, the questing ticks were collected by the flagging method in northeastern China. These ticks were identified to species following the methods detailed in previous studies ^35, 36^. Every 10 ticks were pooled into 1.5 mL Eppendorf tubes based on the collection sites and species, and subsequently stored at -80°C until use.

### RNA library construction and sequencing

After washing with 75% ethanol and RNA/DNA-free water, pooled ticks in tubes were added with 500 μL Dulbecco’s modified Eagle’s minimum essential medium (DMEM) and two stainless steel beads (3 mm diameter), and crushed using the Tissuelyser (Jingxin, Shanghai, China) at 70 Hz for 2 min. The lysates were centrifuged at 12000 rpm for 10 min at 4°C, and the supernatant was further pooled for library construction according to the collection sites and species (***Supplementary Table 8***). After digested with micrococcal nuclease (NEB, USA) in 37°C for 2 h, the pooled samples were used for viral RNA extraction with the TIANamp Virus RNA kit (TIANGEN, Beijing, China). The extracted RNA was subjected to metagenomic sequencing with the Illumina NovaSeq 6000 System at Tanpu Biological Technology (Shanghai, China), as detailed elsewhere ^19^.

### Transcriptome analysis and nairoviruses confirmation

Transcriptome analysis was conducted as described elsewhere ^19^. In summary, following the trimming and elimination of low-quality reads, the paired-raw reads were purified by eliminating ribosomal RNA, host contaminants, and bacterial sequences using the BBMap program. Subsequently, contigs were assembled utilizing SPAdes v3.14.1 and SOAPdenovo v2.04 ^37, 38^. Subsequent to a comparison with the non-redundant nucleotide (nt) and protein (nr) databases retrieved from GenBank through BLAST+ v2.10.0, the assembled contigs were filtered to eliminate host and bacterial sequences. The relative abundance of the identified viruses was evaluated by mapping the reads back onto the assembled contigs with Bowtie2 v2.3.3.1.

Following a comparison with the NCBI nucleotide and viral RefSeq databases via BLAST (V2.10.0+), the contigs assembled for nairoviruses were selected, and confirmation of the nairovirus-positive tick pools was achieved through RT-qPCR using primers designed from these contigs. The complete genomes of these nairoviruses were amplified using semi-nested PCR and the rapid amplification of cDNA ends (RACE) (***Supplementary Table 9***).

### Virus classification

All the viruses identified in this study were classified according to the latest International Committee on Taxonomy of Viruses (ICTV) report of virus taxonomy (https://talk.ictvonline.org/ictvreports/ictv_online_report/). A novel viral species should be satisfied with one of the following conditions as described before^39^, namely, (i) <80% nucleotide (nt) identity across the complete genome; or (ii) <90% amino acid (aa) identity of the RNA-dependent RNA polymerase (RdRp) domain with the known viruses. All novel viruses were named as the collection sites that the virus was first identified, followed by common viral names according to their taxonomy. All the viral strains would be marked with ‘Northeastern (NE)’ to distinguish them from the virus strains identified in other studies.

### Sequence analyses

Potential open reading frames (ORFs) in the viral sequences were predicted using ORFfinder (https://www.ncbi.nlm.nih.gov/orffinder/). For each of the predicted nairoviruses protein sequences, we used the TMHMM (http://www.cbs.dtu.dk/services/TMHMM2.0) program to predict the transmembrane structure domain, SignalP (https://services.healthtech.dtu.dk/services/SignalP-6.0/) to determine signal sequences, ProP (https://services.healthtech.dtu.dk/services/ProP-1.0/) to predict proprotein convertase cleavage sites, and NetNGlyc (https://services.healthtech.dtu.dk/services/NetNGlyc-1.0/) to identify N-linked glycosylation sites. CD-search (https://www.ncbi.nlm.nih.gov/Structure/cdd/wrpsb.cgi) to predict the conserved domain of protein sequences. These proteins were then aligned with their corresponding homologs by using ClustalW available within MEGA 7.0.

### Mapping of tick-borne nairoviruses in northeastern China

To map the distribution of tick-borne nairoviruses in northeastern China, a thorough search was conducted across five databases: Web of Science, PubMed, China National Knowledge Infrastructure, China WanFang Database, and Chinese Scientific Journal Database. This search focused on studies published before November 1, 2024, utilizing the keywords “Tick” and “China” Only studies that offered precisely delineated outcomes, such as the presence of tick-borne nairoviruses and the collection of ticks below the prefecture-level city, were integrated into our database. In situations where a tick-borne nairovirus was reported several times within the same county during the study period, it was accounted for only once in our analyses. The data derived from these studies, together with information from other sources, were merged to establish a comprehensive database at the prefecture-level for the final analysis.

### Sequence identities and phylogenetic analyses

After filtering out the partial or poor-quality sequences, the complete genomes or L segments of nairoviruses identified in northeastern China and the viral strains closely related to these viruses were downloaded from GenBank and employed for similarity and phylogenetic analyses in conjunction with the sequences amplified in this study (***Supplementary Table 10***). Viral nucleotide or amino acid sequences were aligned by means of Clustal W accessible within MEGA 7.0, and evaluated for sequence identities with the MegAlign program available within the DNAstar package V7.0. The amino acid sequences representing the RdRp, GPC, and NP proteins of nairoviruses were used for phylogenetic analysis employing the maximum-likelihood method in IQ-TREE version 2.3.6, with the LG+I+G4 and Q. yeast+I+R5 models. A bootstrapping procedure consisting of 1000 replicates was performed, with bootstrap values exceeding 70 regarded as statistically significant and depicted in the resulting trees.

### Bayesian phylodynamics analysis

The representative strains of nairoviruses with complete S segments and clear collection sites, discovered in northeastern China, along with other closely related viral species from outside the area, were used for the analysis of Bayesian phylodynamics (***Supplementary Table 7***). The analysis was performed utilizing the BEAST 1.10.4 software package, incorporating a relaxed lognormal molecular clock and constant population size models ^40^. The optimal nucleotide substitution model was evaluated using IQ-TREE 2.1.3 software according to the Bayesian Information Criterion, resulting in GTR+G4 and HKY+G4 models. This analysis comprised independent runs over 100 million generations, with sampling occurring every 10,000 generations. Tracer v.1.7 was employed to examine convergence, and an ESS greater than 200 is regarded as relevant ^41^. TreeAnnotator was utilized to summarize the Maximum Clade Credibility (MCC) phylogenetic tree, with the first 10% of the trees being burn-in. FigTree v1.4.3 was adopted to visualize the MCC tree ^42^. The Bayes factor (BF) and posterior probability (PP) were computed via SpreaD3 software, and the principal migration pathways were summarized based on BF > 3 and PP > 0.5 ^29^.

### Virus isolation

To evaluate the pathogenic risk of the newly discovered tick-borne nairoviruses, the supernatants of JANV, YBNTV-1, SXTV-2, YCNV, and HCNV positive tick samples were incubated in Vero, BHK-21, MDBK, and HuH-7 cell lines for virus isolation. Briefly, the lysate of supernatant tick samples was diluted 20-fold in DMEM, filtered through a syringe-driven 0.22-μm filter, and then applied onto the cell monolayer for 1 hour. Subsequently, the cells were maintained in DMEM supplemented with 2% fetal calf serum at 37°C in a 5% carbon dioxide atmosphere. After three passages, the cells were observed for cytopathic effects, and the cells and culture supernatant were collected for viral RNA detection by RT-qPCR.

### Immunofluorescence assay

Vero cells were infected with JANV or YBNTV-1. At 1- or 4-days post-infection, the cells were fixed with 4% paraformaldehyde for 30 minutes and subsequently rinsed three times with PBS. After permeabilization with 0.1% Triton X-100 for 20 minutes, the cells were incubated with anti-JANV rabbit sera or anti-YBNTV-1 mouse sera diluted 1:100 in PBS, overnight at 4°C, followed by three washes with PBS. Antigens were visualized using goat anti-rabbit or anti-mouse IgG (H+L) conjugated with FITC (Beyotime, China), diluted 1:100 in PBS. The samples were then counterstained with DAPI solution (Beyotime, China) and washed three times with PBS. Images were captured using an inverted microscope (Olympus).

### One-step growth curve

The isolated viral strains were employed for the one-step growth curve assay. After being incubated in 24-well plates overnight at 37°C, the Vero, 293T, WISH, HuH-7, and THP-1 cells were incubated with multiplicity of infection of 0.001 of the isolated virus. The culture supernatant was collected every two days and used for viral copy detection by RT-qPCR.

### Electron Microscopy

Four days after infection with tick-borne nairoviruses, the Vero cells were prepared for electron-microscopic analysis. After centrifugation, the cell pellets were fixed with a 2.5% glutaraldehyde solution in 0.1 M sodium cacodylate buffer, postfixed with 1% osmium tetroxide, dehydrated in escalating concentrations of ethanol solutions, and embedded in epoxy resin. The ultrathin sections were stained with 2% uranyl acetate and lead citrate and subsequently examined with a transmission electron microscope. The culture supernatants were centrifuged for the collection of viruses, and the deposits were negatively stained with 2% phosphotungstic acid and observed under the transmission electron microscope as described before^43^.

## Data availability

All sequences generated in this study were uploaded to the GenBank of the National Center for Biotechnology Information (NCBI), with accession numbers: PQ47556–PQ475738. Source data are provided with this paper.

## Acknowledgements

This study was supported by the National Key Research and Development Program of China (2022YFC2601900), Science and Technology Program of the Joint Fund of Scientific Research for the Public Hospitals of Inner Mongolia Academy of Medical Sciences, China (2023GLLH0342 and 2023GLLH0335), the National Natural Science Foundation of China (82272327), and the Outstanding Youth Foundation of National Natural Science Foundation of Jilin Province, China (20240101017JC).

## Author contributions

Z. Wang designed the study. Z.L, Z. Wei did the laboratory tests. Z.L, Z. Wei, and Y.L did the data analysis. Z.L, Z. Wei, Y.L, and Z. Wang wrote the paper. All authors contributed to the review and revision of the report. All authors had full access to all the data in the study and had final responsibility for the decision to submit for publication.

## Additional information

## Supplementary information

**Extended Data Table 1** Amino acid similarity (%) of RDRP (upper right) and nucleotide sequence similarity (%) of segment L (lower left) between WELV and the representative Orthonairovirus in the family Nairoviridae.

**Extended Data Table 2** Amino acid similarity (%) of GPC (upper right) and nucleotide sequence similarity (%) of segment M (lower left) between WELV and the representative Orthonairovirus in the family Nairoviridae.

**Extended Data Table 3** Amino acid similarity (%) of NP (upper right) and nucleotide sequence similarity (%) of segment S (lower left) between WELV and the representative Orthonairovirus in the family Nairoviridae.

**Extended Data Table 4** Amino acid similarity (%) of RDRP (upper right) and nucleotide sequence similarity (%) of segment L (lower left) between SGLV and the representative Orthonairovirus in the family Nairoviridae.

**Extended Data Table 5** Amino acid similarity (%) of GPC (upper right) and nucleotide sequence similarity (%) of segment M (lower left) between SGLV and the representative Orthonairovirus in the family Nairoviridae.

**Extended Data Table 6** Amino acid similarity (%) of NP (upper right) and nucleotide sequence similarity (%) of segment S (lower left) between SGLV and the representative Orthonairovirus in the family Nairoviridae.

**Extended Data Table 7** Amino acid similarity (%) of RDRP (upper right) and nucleotide sequence similarity (%) of segment L (lower left) between YEZV and the representative Orthonairovirus in the family Nairoviridae.

**Extended Data Table 8** Amino acid similarity (%) of GPC (upper right) and nucleotide sequence similarity (%) of segment M (lower left) between YEZV and the representative Orthonairovirus in the family Nairoviridae.

**Extended Data Table 9** Amino acid similarity (%) of NP (upper right) and nucleotide sequence similarity (%) of segment S (lower left) between YEZV and the representative Orthonairovirus in the family Nairoviridae.

**Extended Data Table 10** Amino acid similarity (%) of RDRP (upper right) and nucleotide sequence similarity (%) of segment L (lower left) between BJNV and the representative Norwavirus in the family Nairoviridae.

**Extended Data Table 11** Amino acid similarity (%) of NP (upper right) and nucleotide sequence similarity (%) of segment S (lower left) between BJNV and the representative Norwavirus in the family Nairoviridae.

