## Supplementary information for "The extensive diversity of nairoviruses reveals the complexity and high risk of tick-borne diseases in northeastern China"

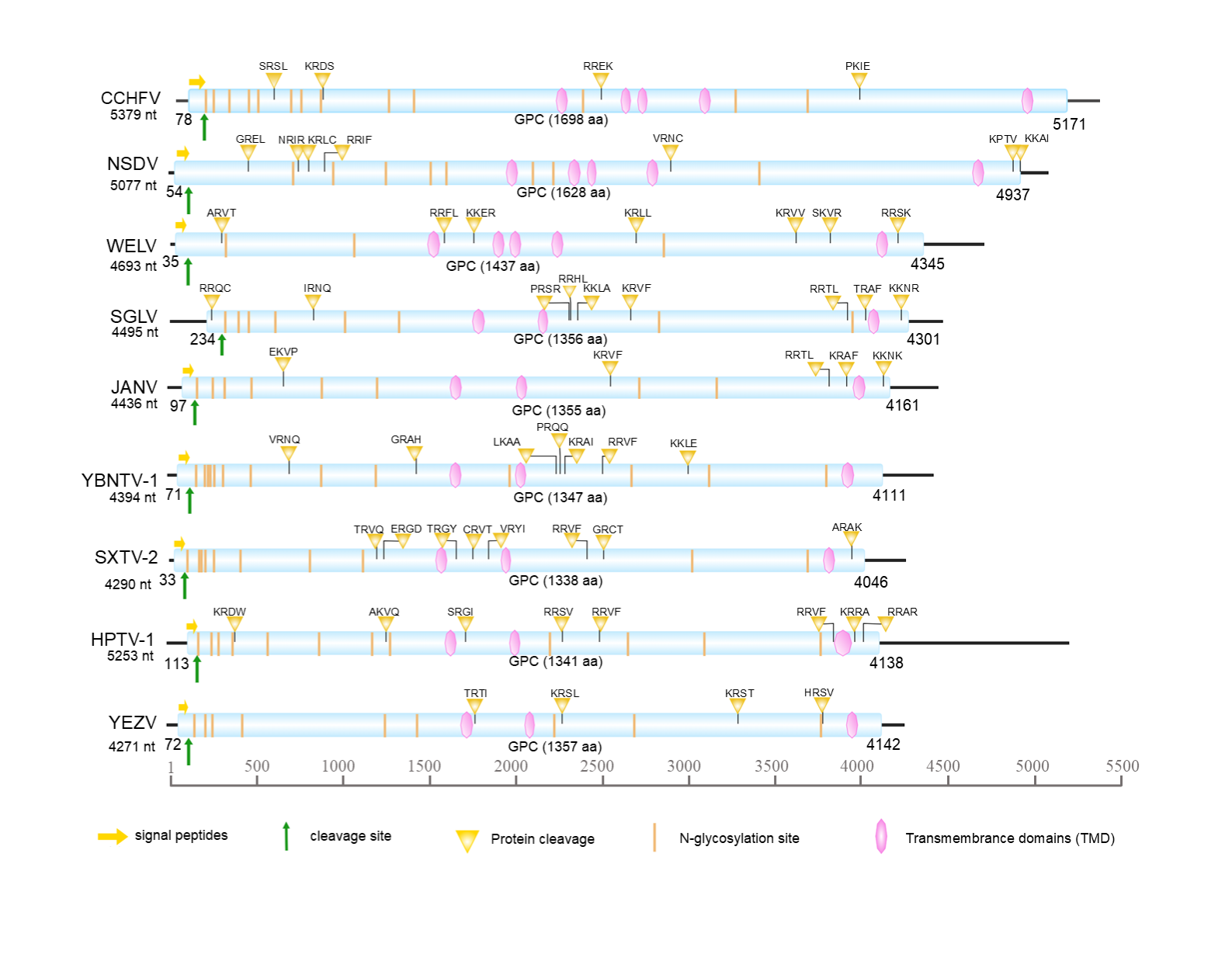

**Supplementary Fig. 1 Genomic characterization of the Medium (M) segment of nairoviruses identified in northeastern China.** The predicted open reading frame (ORF) of M segment that encodes glycoprotein precursor (GPC) is shaded in blue. CCHFV were used as a reference viral species. Abbreviations: CCHFV, Crimean Congo haemorrhagic fever virus; NSDV, Nairobi sheep disease virus;WELV, Wetland virus; SGLV, Songling virus; JANV, Ji’an nairovirus; YBNTV-1,Yianbian Nairo tick virus 1; SXTV-2, Shanxi tick virus 2; YEZV, Yezo virus; BJNV, Beiji nairovirus; YCNV, Yichun nairovirus; HCNV, Hunchun nairovirus.

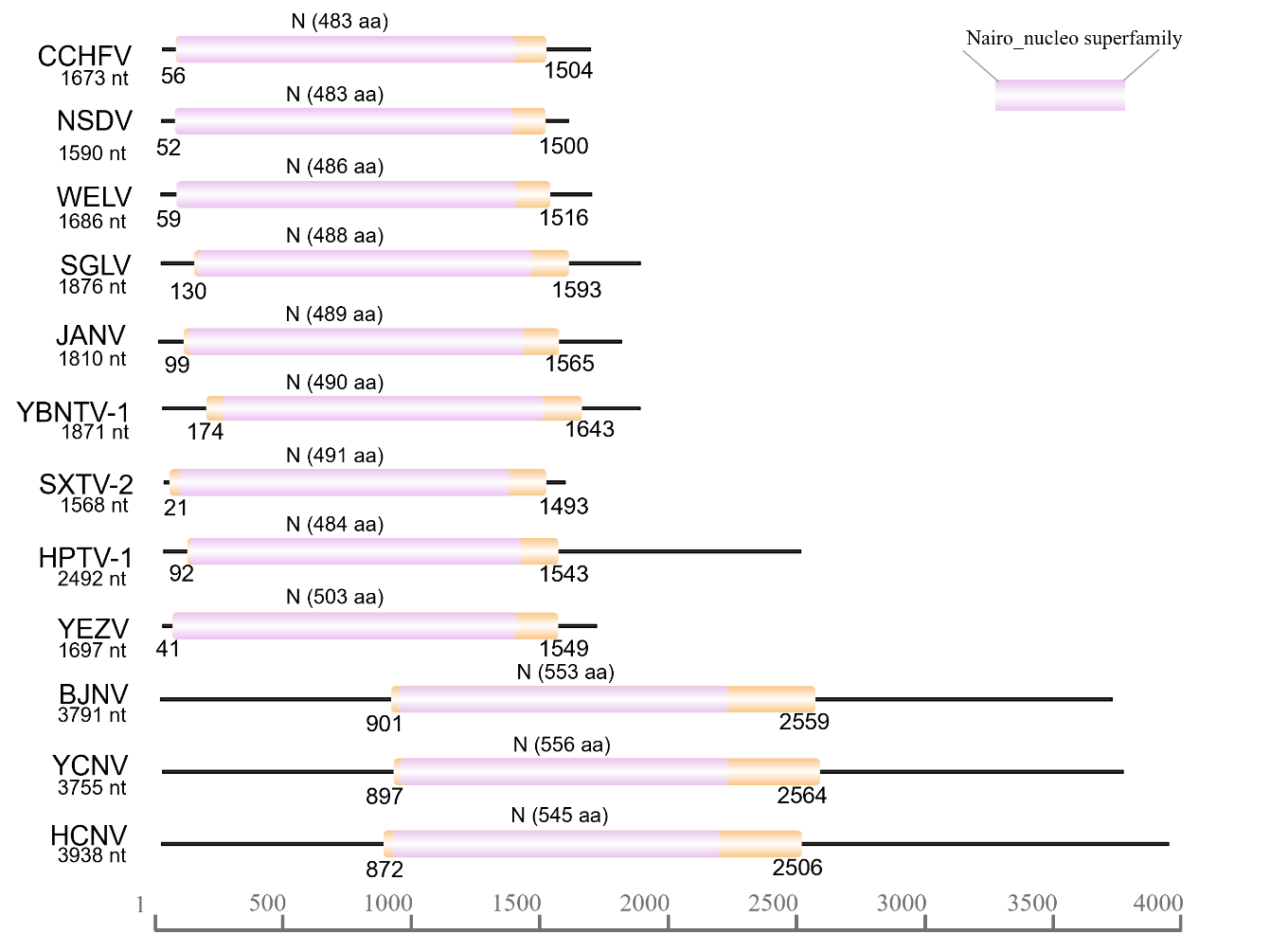

**Supplementary Fig. 2 Genomic characterization of the Small (S) segment of nairoviruses identified in northeastern China.** The predicted open reading frame (ORF) of S segment that encodes nucleoprotein (N) is shaded in orange. CCHFV were used as a reference viral species. Abbreviations: CCHFV, Crimean Congo haemorrhagic fever virus; NSDV, Nairobi sheep disease virus;WELV, Wetland virus; SGLV, Songling virus; JANV, Ji’an nairovirus; YBNTV-1,Yianbian Nairo tick virus 1; SXTV-2, Shanxi tick virus 2; YEZV, Yezo virus; BJNV, Beiji nairovirus; YCNV, Yichun nairovirus; HCNV, Hunchun nairovirus.

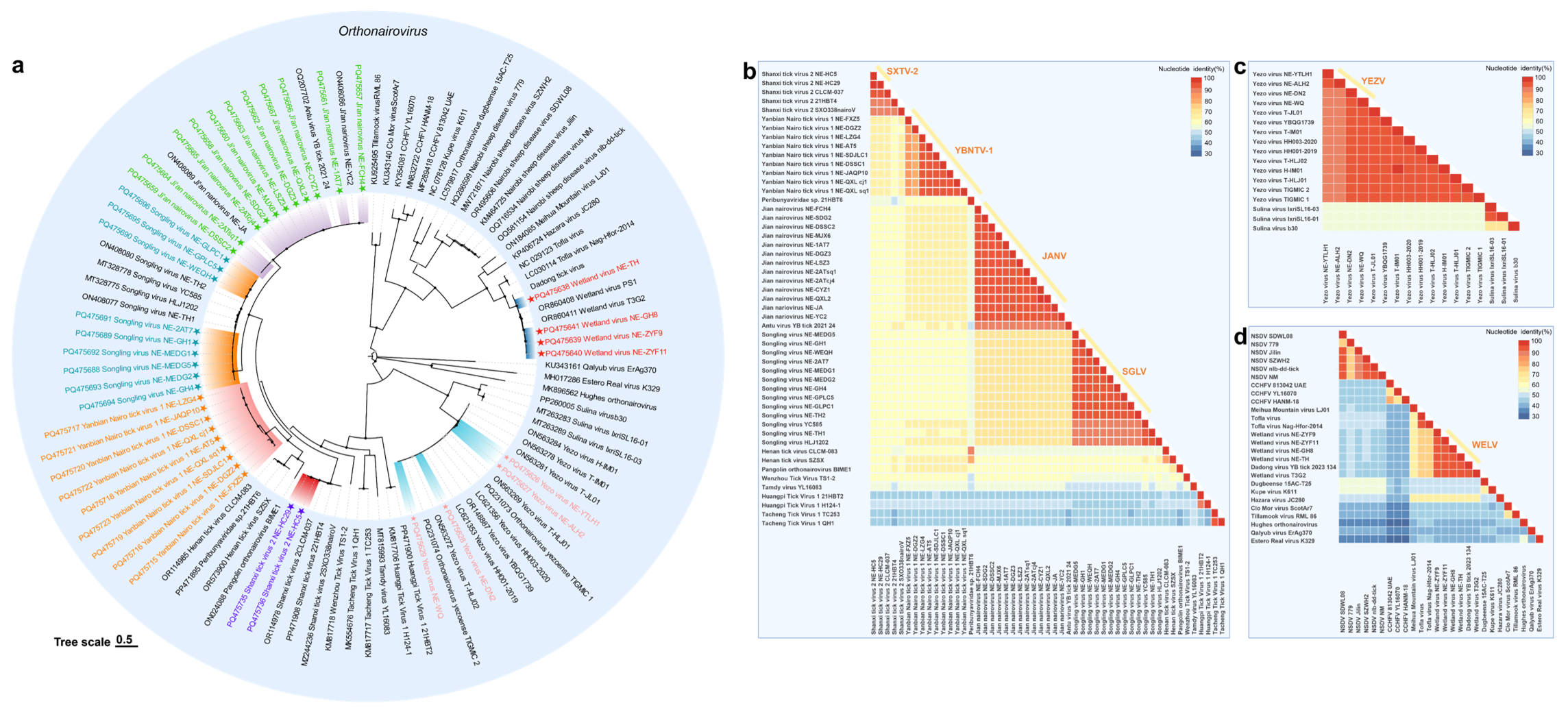

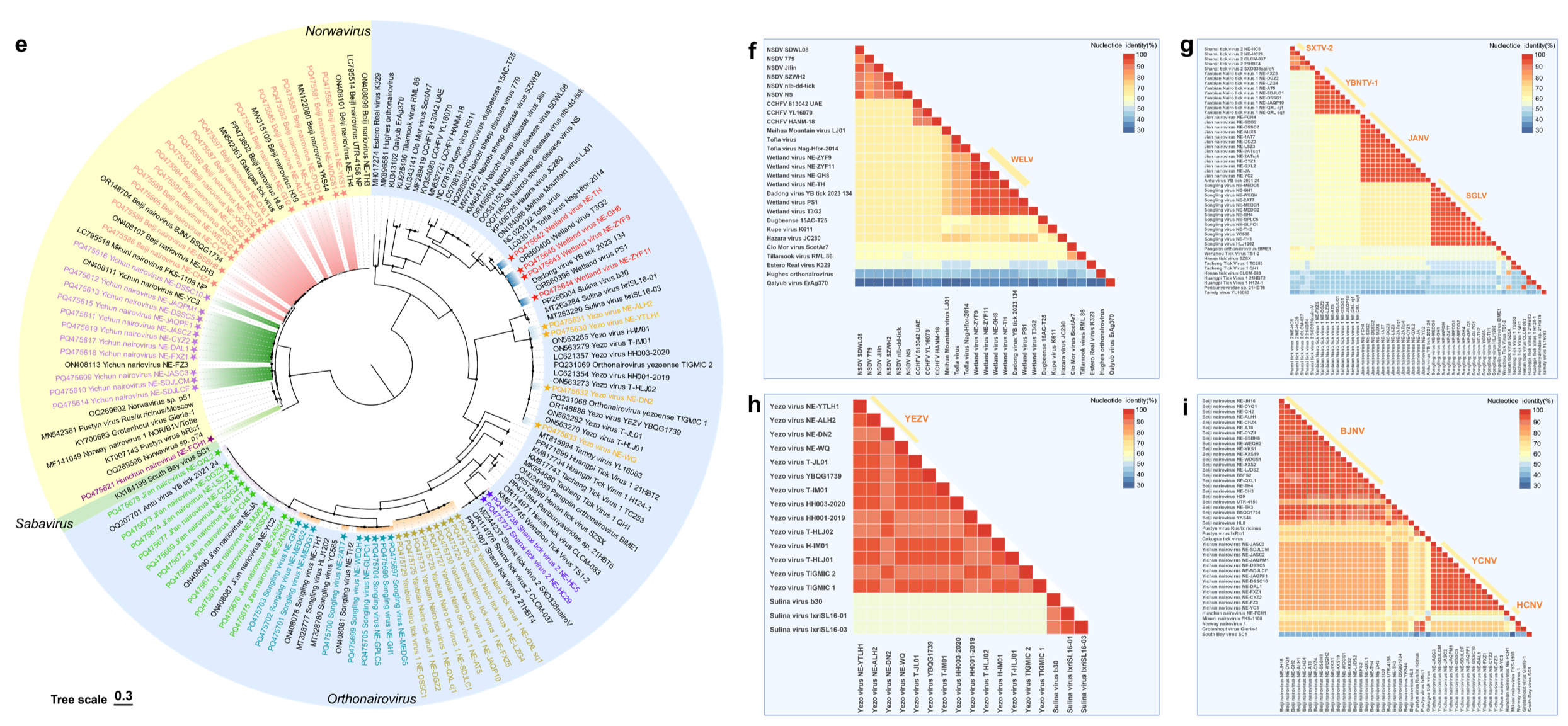

**Supplementary Fig. 3 Phylogenetic and homology analysis of the identified tick-borne nairoviruses in northeastern China.**

**a,e** Phylogenetic trees based on the glycoprotein (GPC), and nucleocapsid (N) protein amino acid sequences of nairoviruses. Tree trees were constructed by the maximum likelihood method using the best-fit models (Q. yeast+I+R5 for GPC protein, and LG+I+G4 for N protein). The newly identified nairoviruses in the present study were labeled by symbol stars. **b-d, f-i** Nucleotide sequence identities of the complete Segment M and S of the four nairovirus groups (ⅰ-ⅳ). Abbreviations: WELV, Wetland virus; SGLV, Songling virus; JANV, Ji’an nairovirus; YBNTV-1, Yianbian Nairo tick virus 1; SXTV-2, Shanxi tick virus 2; YEZV, Yezo virus; BJNV, Beiji nairovirus; YCNV, Yichun nairovirus; HCNV, Hunchun nairovirus.

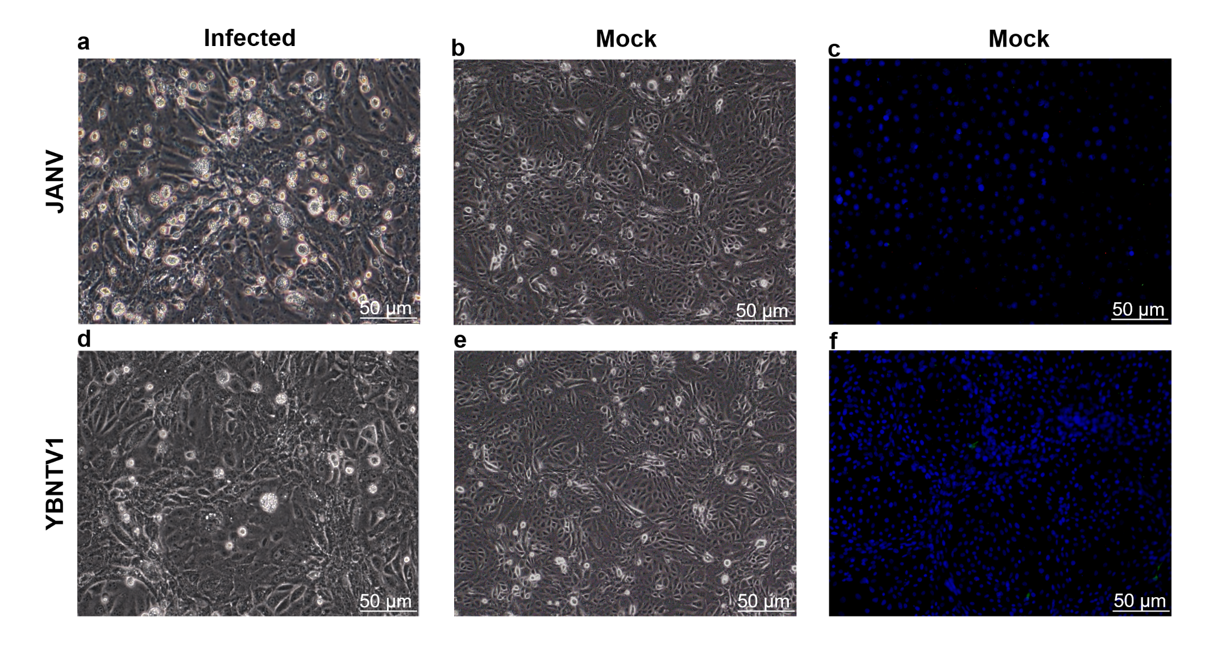

**Supplementary Fig. 4** The cytopathic effects (CPEs) induced by JANV and YBNTV1 or mock in Vero cells at 4 dpi.

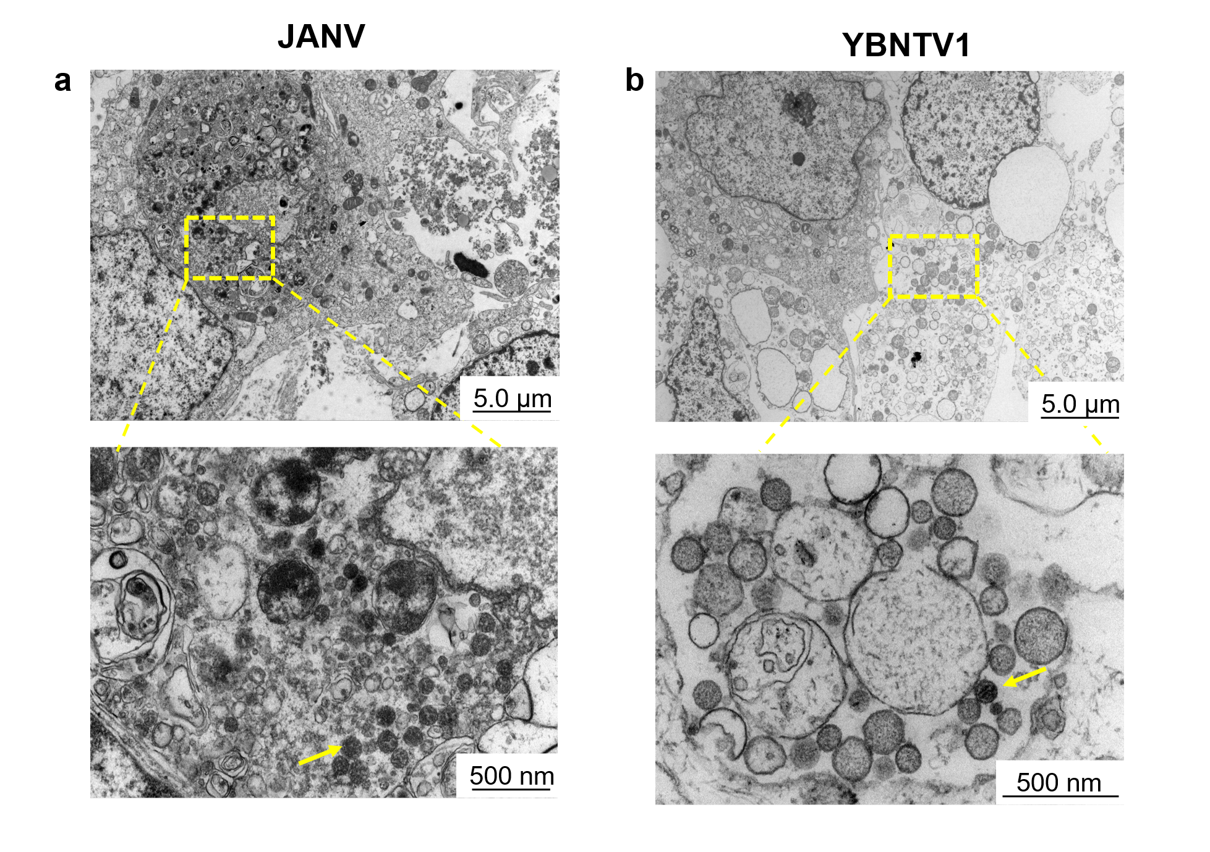

**Supplementary Fig. 5** Transmission electron microscopy showed JANV (a) and YBNTV1 (b) in the cytoplasm of infected Vero cells (yellow arrows). Electron images are representative of at least two independent experiments.

| **Supplementary Table 1.** Information of tick specimen collection in northeastern China. | | | | |  |  |  |
| --- | --- | --- | --- | --- | --- | --- | --- |
|  | **Location** | ***Dermacentor nuttalli*** | ***Haemaphysalis concinna*** | ***Haemaphysalis longicornis*** | | ***Ixodes persulcatus*** | **Total (no.)** |
| **Inner Mongolia** | Genhe | nd | 90 | nd | | 56 | 146 |
|  | Argun | 77 | nd | nd | | 5 | 82 |
|  | Oroqen | 43 | 42 | nd | | 39 | 124 |
|  | Yakeshi | 648 | 307 | nd | | 232 | 1187 |
|  | New Barag West Banner | 2 | nd | nd | | 180 | 182 |
|  | Tongliao | 60 | nd | nd | | nd | 60 |
|  | Chifeng | 118 | nd | nd | | nd | 118 |
| **Heilongjiang** | Tahe | nd | 113 | nd | | 220 | 333 |
|  | Xinlin | 1 | 27 | nd | | 34 | 62 |
|  | Mudanjiang | nd | nd | 20 | | 49 | 69 |
| **Jilin** | Wangqing | nd | 387 | 52 | | 140 | 579 |
|  | Hunchun | 8 | 982 | 571 | | 85 | 1646 |
|  | Tumen | 11 | 76 | 48 | | 6 | 141 |
|  | Helong | nd | 79 | 63 | | nd | 142 |
|  | Antu | 1197 | 27 | 35 | | 388 | 1647 |
|  | Jiaohe | 139 | nd | nd | | nd | 139 |
|  | Linjiang | nd | 51 | 61 | | 295 | 407 |
|  | Ji'an | nd | 92 | 68 | | 36 | 196 |
| **Liaoning** | Dandong | nd | nd | 237 | | 6 | 243 |
|  | Benxi | nd | 156 | 174 | | 84 | 414 |
| **Sub-total** | | 2304 | 2429 | 1329 | | 1855 | 7917 |
| *nt, no data | | | | | | | |

| **Supplementary Table 2.** Sampling information of RNA sequencing. | | | |  |  |
| --- | --- | --- | --- | --- | --- |
| **Sequencing libraries** | **clean data (GB)** | **non-rRNA (reads)** | **Contings** | **RNA viruses** | **nairoviruses** |
| XueXiong | 3.51 | 25311091 | 251778 | 43 | 7 |
| LNXUE | 4.07 | 29422200 | 982151 | 261 | 1 |
| JHGE | 4.39 | 31186680 | 1161384 | 190 | 1 |
| 22QUAN | 5.55 | 38227475 | 616074 | 241 | 35 |
| NEIQUAN | 3.36 | 22476859 | 891389 | 534 | 21 |
| 22XUE | 5.18 | 35237588 | 779719 | 182 | 32 |
| LVGE | 4.11 | 27143597 | 996350 | 205 | 34 |
| LVXUE | 4.12 | 26971157 | 900450 | 128 | 4 |
| LVQUAN | 3.38 | 22029869 | 888638 | 341 | 43 |
| XQ QUAN | 2.74 | 17742317 | 903074 | 446 | 14 |
| 927GEXUE | 5.11 | 34808070 | 753699 | 289 | 11 |
| 927GE | 8.41 | 61380022 | 427666 | 219 | 5 |
| 23GE | 7.59 | 59749399 | 356941 | 1485 | 0 |
| 23SQXUE | 4.63 | 31569547 | 745121 | 497 | 12 |
| 23CJXUE | 7.41 | 51095217 | 899548 | 370 | 13 |
| 23QUANF | 8.51 | 59921870 | 783710 | 477 | 60 |
| 23QUANM | 9.19 | 66577011 | 696625 | 562 | 16 |
| GE1 | 2.67 | 21440746 | 312156 | 179 | 13 |
| GE2 | 2.91 | 22048566 | 106331 | 120 | 1 |
| QUAN | 3.24 | 22376013 | 108277 | 343 | 9 |
| XUE | 2.91 | 22537738 | 278314 | 319 | 63 |
| WGQXUE | 3.61 | 22334344 | 898307 | 348 | 20 |
| CJSQ | 7.07 | 48234690 | 767977 | 431 | 73 |
| 725SQCJ | 5.07 | 33781228 | 877300 | 307 | 23 |
| 725QUAN | 7.22 | 50872559 | 405045 | 371 | 28 |
| 718SQ | 7.54 | 52466994 | 738682 | 317 | 30 |
| 718CJ | 7.93 | 53190449 | 863583 | 275 | 37 |
| **Total** | **141.43** | **990133296** | **15250404** | **9480** | **606** |

| **Supplementary Table 3.** The relative virus abundances of the identified viruses in the present study. | | | | | | | | | | | |
| --- | --- | --- | --- | --- | --- | --- | --- | --- | --- | --- | --- |
| **Species** | **Province** | **Location** | **Viral copies/μL (Log10)** | | | | | | | | |
|  |  |  | **BJNV** | **YCNV** | **HCNV** | **YEZV** | **SXTV2** | **YBNTV1** | **JANV** | **SGLV** | **THNV** |
| *Dermacentor nuttalli* | Inner Mongolia | Yakeshi | nd | nd | nd | nd | nd | nd | nd | nd | 4.57 |
|  |  | Oroqen | nd | nd | nd | nd | nd | nd | nd | 2.66 | nd |
| *Haemaphysalis longicornis* | Jilin | Helong | nd | nd | nd | nd | nd | 5.37 | nd | 3.70 | nd |
|  |  | Linjiang | nd | nd | nd | nd | nd | nd | 4.74 | nd | nd |
|  |  | Antu | nd | nd | nd | nd | nd | nd | 4.47 | nd | nd |
|  |  | Hunchun | nd | nd | nd | nd | 4.20 | nd | 3.07 | nd | nd |
| *Haemaphysalis concinna* |  | Antu | nd | nd | nd | nd | nd | nd | 5.45 | nd | nd |
|  |  | Tumen | nd | nd | nd | nd | nd | nd | 5.15 | nd | nd |
|  |  | Helong | nd | nd | nd | nd | nd | 3.63 | 4.75 | 2.53 | nd |
|  |  | Hunchun | nd | nd | nd | nd | 4.55 | nd | 4.27 | nd | nd |
|  |  | Wangqing | nd | nd | nd | nd | nd | 2.94 | 4.26 | nd | nd |
|  |  | Ji'an | nd | nd | nd | nd | nd | nd | 3.94 | nd | nd |
|  |  | Linjiang | nd | nd | nd | nd | nd | 2.34 | nd | nd | nd |
|  | Heilongjiang | Tahe | nd | nd | nd | nd | nd | nd | nd | nd | 3.94 |
|  | Inner Mongolia | Genhe | nd | nd | nd | nd | nd | nd | nd | 3.99 | 3.94 |
|  |  | Argun | nd | nd | nd | nd | nd | nd | nd | 4.08 | nd |
|  |  | Yakeshi | 2.01 | nd | nd | nd | nd | nd | nd | 3.16 | nd |
|  |  | Oroqen | 1.35 | nd | nd | nd | nd | nd | nd | 2.48 | nd |
| *Ixodes persulcatus* |  | Yakeshi | 5.22 | nd | nd | 3.47 | nd | nd | nd | nd | nd |
|  |  | Oroqen | 5.22 | nd | nd | 2.36 | nd | nd | nd | nd | nd |
|  |  | Genhe | 4.72 | nd | nd | nd | nd | nd | nd | nd | nd |
|  |  | Argun | 3.73 | nd | nd | nd | nd | nd | nd | nd | nd |
|  |  | New Barag Right Banner | 2.97 | nd | nd | nd | nd | nd | nd | nd | nd |
|  | Heilongjiang | Xinlin | 4.91 | nd | nd | nd | nd | nd | nd | nd | nd |
|  |  | Dongning | 3.48 | nd | nd | 2.57 | nd | nd | nd | nd | nd |
|  | Liaoning | Benxi | 4.56 | 5.23 | nd | nd | nd | nd | nd | nd | nd |
|  | Jilin | Hunchun | 4.33 | nd | 5.04 | nd | nd | nd | nd | nd | nd |
|  |  | Antu | 4.16 | nd | nd | nd | nd | nd | nd | nd | nd |
|  |  | Ji'an | 4.01 | 3.89 | nd | nd | nd | nd | nd | nd | nd |
|  |  | Linjiang | 3.96 | 3.46 | nd | nd | nd | nd | nd | nd | nd |
|  |  | Wangqing | 2.99 | 3.27 | nd | 3.74 | nd | nd | nd | nd | nd |
| *nd, no data | | | | | | | | | | | |

| **Supplementary Table 4.** Detection of nairovirus RNA in ticks in northeastern China. | | | |  |
| --- | --- | --- | --- | --- |
| **Virus** | **Species** | **Province** | **Number of ticks** | **Prevalance (95% CI)** |
| BJNV | *Ixodes persulcatus* | Heilongjiang | 83 | 2.59 (0.48-8.66) |
|  | *Ixodes persulcatus* | Jilin | 968 | 7.23 (5.49-9.39)^a^ |
|  | *Ixodes persulcatus* | Liaoning | 90 | 1.08 (0.06-5.09)^ab^ |
|  | *Ixodes persulcatus* | Inner Mongolia | 512 | 8.96 (6.32-12.49)^b^ |
|  | *Haemaphysalis concinna* | Inner Mongolia | 486 | 1.76 (0.83-3.31) |
| YCNV | *Ixodes persulcatus* | Jilin | 968 | 2.62 (1.72-3.85)^a^ |
|  | *Ixodes persulcatus* | Liaoning | 90 | 15.57 (7.06-32.55)^a^ |
| HCNV | *Ixodes persulcatus* | Jilin | 968 | 0.21 (0.04-0.67) |
| JANV | *Haemaphysalis concinna* | Jilin | 1665 | 0.94 (0.55-1.51) |
|  | *Haemaphysalis longicornis* | Jilin | 814 | 0.89 (0.40-1.76) |
| YBNTV1 | *Haemaphysalis concinna* | Jilin | 1665 | 0.49 (0.23-0.93) |
|  | *Haemaphysalis longicornis* | Jilin | 814 | 0.12 (0.01-0.59) |
| SXTV2 | *Haemaphysalis concinna* | Jilin | 1665 | 0.06 (0.00-0.29) |
|  | *Haemaphysalis longicornis* | Jilin | 814 | 0.12 (0.01-0.59) |
| SGLV | *Haemaphysalis concinna* | Jilin | 1665 | 0.12 (0.02-0.39)^a^ |
|  | *Haemaphysalis concinna* | Inner Mongolia | 486 | 6.53 (4.34-9.55)^a^ |
|  | *Haemaphysalis longicornis* | Jilin | 814 | 0.25 (0.04-0.81) |
|  | *Dermacentor silvarum* | Inner Mongolia | 948 | 0.76 (0.34-1.50) |
| YEZV | *Ixodes persulcatus* | Heilongjiang | 83 | 1.23 (0.07-6.12) |
|  | *Ixodes persulcatus* | Jilin | 968 | 0.31 (0.08-0.85) |
|  | *Ixodes persulcatus* | Inner Mongolia | 512 | 1.67 (0.79-3.16) |
| WELV | *Haemaphysalis concinna* | Inner Mongolia | 486 | 0.20 (0.01-0.99) |
|  | *Haemaphysalis concinna* | Heilongjiang | 116 | 0.86 (0.05-4.17) |
|  | *Dermacentor silvarum* | Inner Mongolia | 948 | 0.65 (0.27-1.34) |
| *Prevalence of nairovirus infection in ticks was calculated by using PooledInfRate Excel Add-In version 4.0 (Microsoft, https://www.microsoft.com) and a 1-sample analysis with a bias-corrected maximum likelihood estimation method with 95% CIs and a scale of 100. a,b Significant difference between the tick species (p < 0.05) by Fisher's exact test. | | | | |

| **Supplementary Table 5.** Nucleotide sequences of the identified viruses in the present study. | | | | |  |  |
| --- | --- | --- | --- | --- | --- | --- |
| **Viral species** | **Segments/Gene** | **Collection sites** | **Host species** | **Viral strain** | | **GenBank accession NO.** |
| Beiji nariovirus | L | China: Jiaohe, Jilin | *Ixodes persulcatus* | NE-JH16 | | PQ475566 |
| Beiji nariovirus | S | China: Jiaohe, Jilin | *Ixodes persulcatus* | NE-JH16 | | PQ475582 |
| Beiji nariovirus | L | China: Huma, Heilongjiang | *Ixodes persulcatus* | NE-DYQ1 | | PQ475567 |
| Beiji nariovirus | S | China: Huma, Heilongjiang | *Ixodes persulcatus* | NE-DYQ1 | | PQ475583 |
| Beiji nariovirus | L | China: Genhe, Inner Mongolia | *Ixodes persulcatus* | NE-GH2 | | PQ475568 |
| Beiji nariovirus | S | China: Genhe, Inner Mongolia | *Ixodes persulcatus* | NE-GH2 | | PQ475584 |
| Beiji nariovirus | L | China: Oroqen, Inner Mongolia | *Ixodes persulcatus* | NE-ALH1 | | PQ475569 |
| Beiji nariovirus | S | China: Oroqen, Inner Mongolia | *Ixodes persulcatus* | NE-ALH1 | | PQ475585 |
| Beiji nariovirus | L | China: Hunchun, Jilin | *Ixodes persulcatus* | NE-CHZ4 | | PQ475570 |
| Beiji nariovirus | S | China: Hunchun, Jilin | *Ixodes persulcatus* | NE-CHZ4 | | PQ475586 |
| Beiji nariovirus | L | China: Antu, Jilin | *Ixodes persulcatus* | NE-AT8 | | PQ475571 |
| Beiji nariovirus | S | China: Antu, Jilin | *Ixodes persulcatus* | NE-AT8 | | PQ475587 |
| Beiji nariovirus | L | China: Antu, Jilin | *Ixodes persulcatus* | NE-BSBH8 | | PQ475572 |
| Beiji nariovirus | S | China: Antu, Jilin | *Ixodes persulcatus* | NE-BSBH8 | | PQ475588 |
| Beiji nariovirus | L | China: Yakeshi, Inner Mongolia | *Ixodes persulcatus* | NE-WEQH2 | | PQ475573 |
| Beiji nariovirus | S | China: Yakeshi, Inner Mongolia | *Ixodes persulcatus* | NE-WEQH2 | | PQ475589 |
| Beiji nariovirus | L | China: Yakeshi, Inner Mongolia | *Ixodes persulcatus* | NE-YKS1 | | PQ475574 |
| Beiji nariovirus | S | China: Yakeshi, Inner Mongolia | *Ixodes persulcatus* | NE-YKS1 | | PQ475590 |
| Beiji nariovirus | L | China: Linjiang, Jilin | *Ixodes persulcatus* | NE-WDGS1 | | PQ475575 |
| Beiji nariovirus | S | China: Linjiang, Jilin | *Ixodes persulcatus* | NE-WDGS1 | | PQ475591 |
| Beiji nariovirus | L | China: Linjiang, Jilin | *Ixodes persulcatus* | NE-XXS2 | | PQ475576 |
| Beiji nariovirus | S | China: Linjiang, Jilin | *Ixodes persulcatus* | NE-XXS2 | | PQ475592 |
| Beiji nariovirus | L | China: Linjiang, Jilin | *Ixodes persulcatus* | NE-XXS19 | | PQ475577 |
| Beiji nariovirus | S | China: Linjiang, Jilin | *Ixodes persulcatus* | NE-XXS19 | | PQ475593 |
| Beiji nariovirus | L | China: Linjiang, Jilin | *Ixodes persulcatus* | NE-LJDS2 | | PQ475578 |
| Beiji nariovirus | S | China: Linjiang, Jilin | *Ixodes persulcatus* | NE-LJDS2 | | PQ475594 |
| Beiji nariovirus | L | China: Antu, Jilin | *Ixodes persulcatus* | BSFS2 | | PQ475579 |
| Beiji nariovirus | S | China: Antu, Jilin | *Ixodes persulcatus* | BSFS2 | | PQ475595 |
| Beiji nariovirus | L | China: Wangqing, Jilin | *Ixodes persulcatus* | NE-CYZ4 | | PQ475580 |
| Beiji nariovirus | S | China: Wangqing, Jilin | *Ixodes persulcatus* | NE-CYZ4 | | PQ475596 |
| Beiji nariovirus | L | China: Wangqing, Jilin | *Ixodes persulcatus* | NE-QXL1 | | PQ475581 |
| Beiji nariovirus | S | China: Wangqing, Jilin | *Ixodes persulcatus* | NE-QXL1 | | PQ475597 |
| Yichun nariovirus | L | China:Ji’an, Jilin | *Ixodes persulcatus* | NE-JASC3 | | PQ475598 |
| Yichun nariovirus | S | China:Ji’an, Jilin | *Ixodes persulcatus* | NE-JASC3 | | PQ475609 |
| Yichun nariovirus | L | China: Linjiang, Jilin | *Ixodes persulcatus* | NE-SDJLCM | | PQ475599 |
| Yichun nariovirus | S | China: Linjiang, Jilin | *Ixodes persulcatus* | NE-SDJLCM | | PQ475610 |
| Yichun nariovirus | L | China:Ji’an, Jilin | *Ixodes persulcatus* | NE-JASC2 | | PQ475600 |
| Yichun nariovirus | S | China:Ji’an, Jilin | *Ixodes persulcatus* | NE-JASC2 | | PQ475611 |
| Yichun nariovirus | L | China:Ji’an, Jilin | *Ixodes persulcatus* | NE-JAQPM1 | | PQ475601 |
| Yichun nariovirus | S | China:Ji’an, Jilin | *Ixodes persulcatus* | NE-JAQPM1 | | PQ475612 |
| Yichun nariovirus | L | China: Linjiang, Jilin | *Ixodes persulcatus* | NE-DSSC5 | | PQ475602 |
| Yichun nariovirus | S | China: Linjiang, Jilin | *Ixodes persulcatus* | NE-DSSC5 | | PQ475613 |
| Yichun nariovirus | L | China: Linjiang, Jilin | *Ixodes persulcatus* | NE-SDJLCF | | PQ475603 |
| Yichun nariovirus | S | China: Linjiang, Jilin | *Ixodes persulcatus* | NE-SDJLCF | | PQ475614 |
| Yichun nariovirus | L | China:Ji’an, Jilin | *Ixodes persulcatus* | NE-JAQPF1 | | PQ475604 |
| Yichun nariovirus | S | China:Ji’an, Jilin | *Ixodes persulcatus* | NE-JAQPF1 | | PQ475615 |
| Yichun nariovirus | L | China: Linjiang, Jilin | *Ixodes persulcatus* | NE-DSSC10 | | PQ475605 |
| Yichun nariovirus | S | China: Linjiang, Jilin | *Ixodes persulcatus* | NE-DSSC10 | | PQ475616 |
| Yichun nariovirus | L | China: Benxi, Liaoning | *Ixodes persulcatus* | NE-DAL1 | | PQ475606 |
| Yichun nariovirus | S | China: Benxi, Liaoning | *Ixodes persulcatus* | NE-DAL1 | | PQ475617 |
| Yichun nariovirus | L | China: Wangqing, Jilin | *Ixodes persulcatus* | NE-FXZ1 | | PQ475607 |
| Yichun nariovirus | S | China: Wangqing, Jilin | *Ixodes persulcatus* | NE-FXZ1 | | PQ475618 |
| Yichun nariovirus | L | China: Wangqing, Jilin | *Ixodes persulcatus* | NE-CYZ2 | | PQ475608 |
| Yichun nariovirus | S | China: Wangqing, Jilin | *Ixodes persulcatus* | NE-CYZ2 | | PQ475619 |
| Hunchun nairovirus | L | China: Hunchun, Jilin | *Ixodes persulcatus* | NE-FCH1 | | PQ475620 |
| Hunchun nairovirus | S | China: Hunchun, Jilin | *Ixodes persulcatus* | NE-FCH1 | | PQ475621 |
| Yezo virus | L | China: Yitulihe, Inner Mongolia | *Ixodes persulcatus* | NE-YTLH1 | | PQ475622 |
| Yezo virus | M | China: Yitulihe, Inner Mongolia | *Ixodes persulcatus* | NE-YTLH1 | | PQ475626 |
| Yezo virus | S | China: Yitulihe, Inner Mongolia | *Ixodes persulcatus* | NE-YTLH1 | | PQ475630 |
| Yezo virus | L | China: ALiHe, Inner Mongolia | *Ixodes persulcatus* | NE-ALH2 | | PQ475623 |
| Yezo virus | M | China: ALiHe, Inner Mongolia | *Ixodes persulcatus* | NE-ALH2 | | PQ475627 |
| Yezo virus | S | China: ALiHe, Inner Mongolia | *Ixodes persulcatus* | NE-ALH2 | | PQ475631 |
| Yezo virus | L | China: Dongning, Heilongjiang | *Ixodes persulcatus* | NE-DN2 | | PQ475624 |
| Yezo virus | M | China: Dongning, Heilongjiang | *Ixodes persulcatus* | NE-DN2 | | PQ475628 |
| Yezo virus | S | China: Dongning, Heilongjiang | *Ixodes persulcatus* | NE-DN2 | | PQ475632 |
| Yezo virus | L | China: Wangqing, Jilin | *Ixodes persulcatus* | NE-WQ | | PQ475625 |
| Yezo virus | M | China: Wangqing, Jilin | *Ixodes persulcatus* | NE-WQ | | PQ475629 |
| Yezo virus | S | China: Wangqing, Jilin | *Ixodes persulcatus* | NE-WQ | | PQ475633 |
| Wetland virus | L | China: Genhe, Inner Mongolia | *Haemaphysalis conicinna* | NE-TH | | PQ475634 |
| Wetland virus | M | China: Genhe, Inner Mongolia | *Haemaphysalis conicinna* | NE-TH | | PQ475638 |
| Wetland virus | S | China: Genhe, Inner Mongolia | *Haemaphysalis conicinna* | NE-TH | | PQ475642 |
| Wetland virus | L | China: Yakeshi, Inner Mongolia | *Dermacentor nuttalli* | NE-ZYF9 | | PQ475635 |
| Wetland virus | M | China: Yakeshi, Inner Mongolia | *Dermacentor nuttalli* | NE-ZYF9 | | PQ475639 |
| Wetland virus | S | China: Yakeshi, Inner Mongolia | *Dermacentor nuttalli* | NE-ZYF9 | | PQ475643 |
| Wetland virus | L | China: Yakeshi, Inner Mongolia | *Dermacentor nuttalli* | NE-ZYF11 | | PQ475636 |
| Wetland virus | M | China: Yakeshi, Inner Mongolia | *Dermacentor nuttalli* | NE-ZYF11 | | PQ475640 |
| Wetland virus | S | China: Yakeshi, Inner Mongolia | *Dermacentor nuttalli* | NE-ZYF11 | | PQ475644 |
| Wetland virus | L | China: Tahe, Heilongjiang | *Haemaphysalis conicinna* | NE-GH8 | | PQ475637 |
| Wetland virus | M | China: Tahe, Heilongjiang | *Haemaphysalis conicinna* | NE-GH8 | | PQ475641 |
| Wetland virus | S | China: Tahe, Heilongjiang | *Haemaphysalis conicinna* | NE-GH8 | | PQ475645 |
| Ji’an nariovirus | L | China: Hunchun, Jilin | *Haemaphysalis conicinna* | NE-FCH4 | | PQ475646 |
| Ji’an nariovirus | M | China: Hunchun, Jilin | *Haemaphysalis conicinna* | NE-FCH4 | | PQ475657 |
| Ji’an nariovirus | S | China: Hunchun, Jilin | *Haemaphysalis conicinna* | NE-FCH4 | | PQ475668 |
| Ji’an nariovirus | L | China: Hunchun, Jilin | *Haemaphysalis conicinna* | NE-SDG2 | | PQ475647 |
| Ji’an nariovirus | M | China: Hunchun, Jilin | *Haemaphysalis conicinna* | NE-SDG2 | | PQ475658 |
| Ji’an nariovirus | S | China: Hunchun, Jilin | *Haemaphysalis conicinna* | NE-SDG2 | | PQ475669 |
| Ji’an nariovirus | L | China: Linjiang, Jilin | *Haemaphysalis longicornis* | NE-DSSC2 | | PQ475648 |
| Ji’an nariovirus | M | China: Linjiang, Jilin | *Haemaphysalis longicornis* | NE-DSSC2 | | PQ475659 |
| Ji’an nariovirus | S | China: Linjiang, Jilin | *Haemaphysalis longicornis* | NE-DSSC2 | | PQ475670 |
| Ji’an nariovirus | L | China: Hunchun, Jilin | *Haemaphysalis longicornis* | NE-MJX6 | | PQ475649 |
| Ji’an nariovirus | M | China: Hunchun, Jilin | *Haemaphysalis longicornis* | NE-MJX6 | | PQ475660 |
| Ji’an nariovirus | S | China: Hunchun, Jilin | *Haemaphysalis longicornis* | NE-MJX6 | | PQ475671 |
| Ji’an nariovirus | L | China: Antu, Jilin | *Haemaphysalis conicinna* | NE-1AT7 | | PQ475650 |
| Ji’an nariovirus | M | China: Antu, Jilin | *Haemaphysalis conicinna* | NE-1AT7 | | PQ475661 |
| Ji’an nariovirus | S | China: Antu, Jilin | *Haemaphysalis conicinna* | NE-1AT7 | | PQ475672 |
| Ji’an nariovirus | L | China: Wangqing, Jilin | *Haemaphysalis conicinna* | NE-DGZ3 | | PQ475651 |
| Ji’an nariovirus | M | China: Wangqing, Jilin | *Haemaphysalis conicinna* | NE-DGZ3 | | PQ475662 |
| Ji’an nariovirus | S | China: Wangqing, Jilin | *Haemaphysalis conicinna* | NE-DGZ3 | | PQ475673 |
| Ji’an nariovirus | L | China: Tumen, Jilin | *Haemaphysalis conicinna* | NE-LSZ3 | | PQ475652 |
| Ji’an nariovirus | M | China: Tumen, Jilin | *Haemaphysalis conicinna* | NE-LSZ3 | | PQ475663 |
| Ji’an nariovirus | S | China: Tumen, Jilin | *Haemaphysalis conicinna* | NE-LSZ3 | | PQ475674 |
| Ji’an nariovirus | L | China: Antu, Jilin | *Haemaphysalis conicinna* | NE-2ATsq1 | | PQ475653 |
| Ji’an nariovirus | M | China: Antu, Jilin | *Haemaphysalis conicinna* | NE-2ATsq1 | | PQ475664 |
| Ji’an nariovirus | S | China: Antu, Jilin | *Haemaphysalis conicinna* | NE-2ATsq1 | | PQ475675 |
| Ji’an nariovirus | L | China: Antu, Jilin | *Haemaphysalis longicornis* | NE-2ATcj4 | | PQ475654 |
| Ji’an nariovirus | M | China: Antu, Jilin | *Haemaphysalis longicornis* | NE-2ATcj4 | | PQ475665 |
| Ji’an nariovirus | S | China: Antu, Jilin | *Haemaphysalis longicornis* | NE-2ATcj4 | | PQ475676 |
| Ji’an nariovirus | L | China: Wangqing, Jilin | *Haemaphysalis conicinna* | NE-CYZ1 | | PQ475655 |
| Ji’an nariovirus | M | China: Wangqing, Jilin | *Haemaphysalis conicinna* | NE-CYZ1 | | PQ475666 |
| Ji’an nariovirus | S | China: Wangqing, Jilin | *Haemaphysalis conicinna* | NE-CYZ1 | | PQ475677 |
| Ji’an nariovirus | L | China: Helong, Jilin | *Haemaphysalis conicinna* | NE-QXL2 | | PQ475656 |
| Ji’an nariovirus | M | China: Helong, Jilin | *Haemaphysalis conicinna* | NE-QXL2 | | PQ475667 |
| Ji’an nariovirus | S | China: Helong, Jilin | *Haemaphysalis conicinna* | NE-QXL2 | | PQ475678 |
| Songling virus | L | China: Tahe, Heilongjiang | *Haemaphysalis conicinna* | NE-MEDG5 | | PQ475679 |
| Songling virus | M | China: Tahe, Heilongjiang | *Haemaphysalis conicinna* | NE-MEDG5 | | PQ475688 |
| Songling virus | S | China: Tahe, Heilongjiang | *Haemaphysalis conicinna* | NE-MEDG5 | | PQ475697 |
| Songling virus | L | China: Genhe, Inner Mongolia | *Haemaphysalis conicinna* | NE-GH1 | | PQ475680 |
| Songling virus | M | China: Genhe, Inner Mongolia | *Haemaphysalis conicinna* | NE-GH1 | | PQ475689 |
| Songling virus | S | China: Genhe, Inner Mongolia | *Haemaphysalis conicinna* | NE-GH1 | | PQ475698 |
| Songling virus | L | China: Yakeshi, Inner Mongolia | *Haemaphysalis conicinna* | NE-WEQH | | PQ475681 |
| Songling virus | M | China: Yakeshi, Inner Mongolia | *Haemaphysalis conicinna* | NE-WEQH | | PQ475690 |
| Songling virus | S | China: Yakeshi, Inner Mongolia | *Haemaphysalis conicinna* | NE-WEQH | | PQ475699 |
| Songling virus | L | China: Hunchun, Jilin | *Haemaphysalis conicinna* | NE-2AT7 | | PQ475682 |
| Songling virus | M | China: Hunchun, Jilin | *Haemaphysalis conicinna* | NE-2AT7 | | PQ475691 |
| Songling virus | S | China: Hunchun, Jilin | *Haemaphysalis conicinna* | NE-2AT7 | | PQ475700 |
| Songling virus | L | China: Ergun, Inner Mongolia | *Haemaphysalis conicinna* | NE-MEDG1 | | PQ475683 |
| Songling virus | M | China: Ergun, Inner Mongolia | *Haemaphysalis conicinna* | NE-MEDG1 | | PQ475692 |
| Songling virus | S | China: Ergun, Inner Mongolia | *Haemaphysalis conicinna* | NE-MEDG1 | | PQ475701 |
| Songling virus | L | China: Ergun, Inner Mongolia | *Haemaphysalis conicinna* | NE-MEDG2 | | PQ475684 |
| Songling virus | M | China: Ergun, Inner Mongolia | *Haemaphysalis conicinna* | NE-MEDG2 | | PQ475693 |
| Songling virus | S | China: Ergun, Inner Mongolia | *Haemaphysalis conicinna* | NE-MEDG2 | | PQ475702 |
| Songling virus | L | China: Genhe, Inner Mongolia | *Haemaphysalis conicinna* | NE-GH4 | | PQ475685 |
| Songling virus | M | China: Genhe, Inner Mongolia | *Haemaphysalis conicinna* | NE-GH4 | | PQ475694 |
| Songling virus | S | China: Genhe, Inner Mongolia | *Haemaphysalis conicinna* | NE-GH4 | | PQ475703 |
| Songling virus | L | China: Helong, Jilin | *Haemaphysalis conicinna* | NE-GPLC5 | | PQ475686 |
| Songling virus | M | China: Helong, Jilin | *Haemaphysalis conicinna* | NE-GPLC5 | | PQ475695 |
| Songling virus | S | China: Helong, Jilin | *Haemaphysalis conicinna* | NE-GPLC5 | | PQ475704 |
| Songling virus | L | China: Helong, Jilin | *Haemaphysalis longicornis* | NE-GLPC1 | | PQ475687 |
| Songling virus | M | China: Helong, Jilin | *Haemaphysalis longicornis* | NE-GLPC1 | | PQ475696 |
| Songling virus | S | China: Helong, Jilin | *Haemaphysalis longicornis* | NE-GLPC1 | | PQ475705 |
| Yanbian Nairo tick virus 1 | L | China: Wangqing, Jilin | *Haemaphysalis conicinna* | NE-FXZ5 | | PQ475706 |
| Yanbian Nairo tick virus 1 | M | China: Wangqing, Jilin | *Haemaphysalis conicinna* | NE-FXZ5 | | PQ475715 |
| Yanbian Nairo tick virus 1 | S | China: Wangqing, Jilin | *Haemaphysalis conicinna* | NE-FXZ5 | | PQ475724 |
| Yanbian Nairo tick virus 1 | L | China: Wangqing, Jilin | *Haemaphysalis conicinna* | NE-DGZ2 | | PQ475707 |
| Yanbian Nairo tick virus 1 | M | China: Wangqing, Jilin | *Haemaphysalis conicinna* | NE-DGZ2 | | PQ475716 |
| Yanbian Nairo tick virus 1 | S | China: Wangqing, Jilin | *Haemaphysalis conicinna* | NE-DGZ2 | | PQ475725 |
| Yanbian Nairo tick virus 1 | L | China: Wangqing, Jilin | *Haemaphysalis conicinna* | NE-LZG4 | | PQ475708 |
| Yanbian Nairo tick virus 1 | M | China: Wangqing, Jilin | *Haemaphysalis conicinna* | NE-LZG4 | | PQ475717 |
| Yanbian Nairo tick virus 1 | S | China: Wangqing, Jilin | *Haemaphysalis conicinna* | NE-LZG4 | | PQ475726 |
| Yanbian Nairo tick virus 1 | L | China: Antu, Jilin | *Haemaphysalis conicinna* | NE-AT5 | | PQ475709 |
| Yanbian Nairo tick virus 1 | M | China: Antu, Jilin | *Haemaphysalis conicinna* | NE-AT5 | | PQ475718 |
| Yanbian Nairo tick virus 1 | S | China: Antu, Jilin | *Haemaphysalis conicinna* | NE-AT5 | | PQ475727 |
| Yanbian Nairo tick virus 1 | L | China: Linjiang, Jilin | *Haemaphysalis conicinna* | NE-SDJLC1 | | PQ475710 |
| Yanbian Nairo tick virus 1 | M | China: Linjiang, Jilin | *Haemaphysalis conicinna* | NE-SDJLC1 | | PQ475719 |
| Yanbian Nairo tick virus 1 | S | China: Linjiang, Jilin | *Haemaphysalis conicinna* | NE-SDJLC1 | | PQ475728 |
| Yanbian Nairo tick virus 1 | L | China: Linjiang, Jilin | *Haemaphysalis conicinna* | NE-DSSC1 | | PQ475711 |
| Yanbian Nairo tick virus 1 | M | China: Linjiang, Jilin | *Haemaphysalis conicinna* | NE-DSSC1 | | PQ475720 |
| Yanbian Nairo tick virus 1 | S | China: Linjiang, Jilin | *Haemaphysalis conicinna* | NE-DSSC1 | | PQ475729 |
| Yanbian Nairo tick virus 1 | L | China:Ji’an, Jilin | *Haemaphysalis conicinna* | NE-JAQP10 | | PQ475712 |
| Yanbian Nairo tick virus 1 | M | China:Ji’an, Jilin | *Haemaphysalis conicinna* | NE-JAQP10 | | PQ475721 |
| Yanbian Nairo tick virus 1 | S | China:Ji’an, Jilin | *Haemaphysalis conicinna* | NE-JAQP10 | | PQ475730 |
| Yanbian Nairo tick virus 1 | L | China: Helong, Jilin | *Haemaphysalis longicornis* | NE-QXL cj1 | | PQ475713 |
| Yanbian Nairo tick virus 1 | M | China: Helong, Jilin | *Haemaphysalis longicornis* | NE-QXL cj1 | | PQ475722 |
| Yanbian Nairo tick virus 1 | S | China: Helong, Jilin | *Haemaphysalis longicornis* | NE-QXL cj1 | | PQ475731 |
| Yanbian Nairo tick virus 1 | L | China: Helong, Jilin | *Haemaphysalis conicinna* | NE-QXL sq1 | | PQ475714 |
| Yanbian Nairo tick virus 1 | M | China: Helong, Jilin | *Haemaphysalis conicinna* | NE-QXL sq1 | | PQ475723 |
| Yanbian Nairo tick virus 1 | S | China: Helong, Jilin | *Haemaphysalis conicinna* | NE-QXL sq1 | | PQ475732 |
| Shanxi tick virus 2 | L | China: Hunchun, Jilin | *Haemaphysalis conicinna* | NE-HC29 | | PQ475733 |
| Shanxi tick virus 2 | M | China: Hunchun, Jilin | *Haemaphysalis conicinna* | NE-HC29 | | PQ475735 |
| Shanxi tick virus 2 | S | China: Hunchun, Jilin | *Haemaphysalis conicinna* | NE-HC29 | | PQ475737 |
| Shanxi tick virus 2 | L | China: Hunchun, Jilin | *Haemaphysalis longicornis* | NE-HC5 | | PQ475734 |
| Shanxi tick virus 2 | M | China: Hunchun, Jilin | *Haemaphysalis longicornis* | NE-HC5 | | PQ475736 |
| Shanxi tick virus 2 | S | China: Hunchun, Jilin | *Haemaphysalis longicornis* | NE-HC5 | | PQ475738 |

**Supplementary Table 6.** The distribution of tick-borne nairoviruses in northeastern China.

| **Province** | **Location** | **SGLV** | **JANV** | **YBNTV1** | **YEZV** | **NSDV** | **WELV** | **SXTV2** | **HPTV** | **BJNV** | **YCNV** | **HCNV** |
| --- | --- | --- | --- | --- | --- | --- | --- | --- | --- | --- | --- | --- |
| Liaoning | Benxi | nd | nd | nd | nd | **+** | nd | nd | nd | **+** | **+** | nd |
|  | Dandong | nd | nd | nd | nd | nd | **+** | nd | nd | nd | nd | nd |
|  | Dandong,Fengcheng | nd | nd | nd | nd | **+** | nd | nd | nd | nd | nd | nd |
|  | Jinzhou | nd | nd | nd | nd | nd | **+** | nd | nd | nd | nd | nd |
|  | Liaoyang | nd | nd | nd | nd | nd | **+** | nd | nd | nd | nd | nd |
|  | Huludao | nd | nd | nd | nd | nd | nd | **+** | **+** | nd | nd | nd |
|  |  | nd | nd | nd | nd | nd | nd | nd | nd | nd | nd | nd |
| Heilongjiang | Daxing'anling, Mohe | nd | nd | nd | nd | nd | nd | nd | nd | **+** | nd | nd |
|  | Daxing'anling, Songling | nd | nd | nd | nd | nd | nd | nd | nd | **+** | nd | nd |
|  | Daxing'anling, Xinlin | nd | nd | nd | nd | nd | nd | nd | nd | **+** | nd | nd |
|  | Daxing'anling, Tahe | **+** | nd | nd | **+** | nd | **+** | nd | nd | **+** | nd | nd |
|  | Yichun | **+** | **+** | nd | **+** | nd | nd | nd | nd | **+** | **+** | nd |
|  | Yichun, Fenglin | nd | nd | nd | **+** | nd | nd | nd | nd | nd | nd | nd |
|  | Heihe, NenjiangRiver | nd | nd | nd | **+** | nd | **+** | nd | nd | nd | nd | nd |
|  | Heihe, Wudalianchi | nd | nd | nd | **+** | nd | nd | nd | nd | nd | nd | nd |
|  | Harbin | nd | nd | nd | nd | nd | **+** | nd | nd | nd | nd | nd |
|  | Harbin, Fangzheng | nd | nd | nd | nd | nd | nd | nd | nd | nd | **+** | nd |
|  | Mudanjiang | nd | **+** | nd | **+** | nd | nd | nd | nd | nd | nd | nd |
|  | Mudanjiang, Ningan | nd | nd | nd | **+** | nd | **+** | nd | nd | nd | nd | nd |
|  | Mudanjiang, Dongning | nd | nd | nd | **+** | nd | **+** | nd | nd | **+** | nd | nd |
|  | Mudanjiang, Hailin | nd | nd | nd | **+** | nd | nd | nd | nd | nd | nd | nd |
|  | Daqing | nd | nd | nd | **+** | nd | nd | nd | nd | nd | nd | nd |
|  | Suihua, Zhaodong | nd | nd | nd | **+** | nd | nd | nd | nd | nd | nd | nd |
|  | Jiamusi, Jiaoqu | nd | nd | nd | **+** | nd | nd | nd | nd | nd | nd | nd |
|  | Jiamusi, Huanan | nd | nd | nd | **+** | nd | nd | nd | nd | nd | nd | nd |
|  | Jixi, Hulin | nd | nd | nd | **+** | nd | nd | nd | nd | nd | nd | nd |
|  |  | nd | nd | nd | nd | nd | nd | nd | nd | nd | nd | nd |
| Inner Mongolia | Hulun Buir, Genhe | nd | nd | nd | nd | nd | **+** | nd | nd | **+** | nd | nd |
|  | Hulun Buir, Oroqen | **+** | nd | nd | **+** | nd | **+** | nd | nd | **+** | nd | nd |
|  | Hulun Buir, Yakeshi | **+** | nd | nd | **+** | nd | **+** | nd | nd | **+** | nd | nd |
|  | Hulun Buir, Ewenki | nd | nd | nd | nd | nd | **+** | nd | nd | **+** | nd | nd |
|  | Hulun Buir, Ergun | **+** | nd | nd | nd | nd | **+** | nd | nd | **+** | nd | nd |
|  | Hulun Buir, New Barag West County | nd | nd | nd | nd | nd | **+** | nd | nd | **+** | nd | nd |
|  | Hulun Buir, New Barag East County | nd | nd | nd | nd | nd | **+** | nd | nd | nd | nd | nd |
|  | Hulun Buir, Morin Dawa | nd | nd | nd | nd | nd | nd | nd | nd | **+** | nd | nd |
|  |  | nd | nd | nd | nd | nd | nd | nd | nd | nd | nd | nd |
| Jilin | Jiaohe | nd | nd | nd | nd | nd | nd | nd | nd | **+** | nd | nd |
|  | Baishan, Linjiang | nd | **+** | **+** | nd | nd | **+** | nd | nd | **+** | **+** | nd |
|  | Tonghua, Ji'an | **+** | **+** | **+** | **+** | **+** | **+** | nd | nd | **+** | **+** | nd |
|  | Songyuan | nd | nd | nd | nd | nd | **+** | nd | nd | nd | nd | nd |
|  | Yanbian, Antu | nd | **+** | nd | nd | nd | nd | nd | nd | **+** | **+** | nd |
|  | Yanbian, Hunchun | nd | **+** | nd | nd | **+** | nd | **+** | nd | **+** | nd | **+** |
|  | Yanbian, Wangqing | nd | **+** | **+** | **+** | nd | nd | nd | nd | **+** | **+** | nd |
|  | Yanbian, Longjing | **+** | nd | nd | nd | nd | nd | nd | nd | nd | nd | nd |
|  | Yanbian, Helong | **+** | **+** | nd | **+** | nd | **+** | nd | nd | nd | nd | nd |
|  | Yanbian, Tumen | nd | **+** | nd | nd | nd | nd | nd | nd | nd | nd | nd |
|  | Yanbian, Dunhua | nd | **+** | nd | nd | nd | **+** | nd | nd | **+** | nd | nd |
| +，the distribution of nairoviruses; *nd, no data | | | | | | | | | | | | |

**Supplementary Table 7.** GenBank accession number (nucleotide) for Bayesian phylodynamics analysis.

| **Family** | **Genus** | **Virus** | **Strain** | **State** | **Date** | **Accession NO.** |
| --- | --- | --- | --- | --- | --- | --- |
| *Nairoviridae* | *Norwavirus* | Beiji nairovirus | UTR-4158 | Japan | 2021 | LC795514 |
| *Nairoviridae* | *Norwavirus* | Beiji nairovirus | SBC-4130 | Japan | 2021 | LC795508 |
| *Nairoviridae* | *Norwavirus* | Beiji nairovirus | UTR-4157 | Japan | 2021 | LC795512 |
| *Nairoviridae* | *Norwavirus* | Beiji nairovirus | KKW-0929 | Japan | 2014 | LC795502 |
| *Nairoviridae* | *Norwavirus* | Beiji nairovirus | HL8 | China/Great_Hinggan_Mountains | 2022 | PP473602 |
| *Nairoviridae* | *Norwavirus* | Beiji nairovirus | FKS-1101 | Japan | 2014 | LC795504 |
| *Nairoviridae* | *Norwavirus* | Beiji nairovirus | HL6 | China/Great_Hinggan_Mountains | 2022 | PP473600 |
| *Nairoviridae* | *Norwavirus* | Beiji nairovirus | MNM-3915 | Japan | 2020 | LC795506 |
| *Nairoviridae* | *Norwavirus* | Beiji nairovirus | HL7 | China/Great_Hinggan_Mountains | 2022 | PP473601 |
| *Nairoviridae* | *Norwavirus* | Beiji nairovirus | RUS-4138 | Japan | 2021 | LC795510 |
| *Nairoviridae* | *Norwavirus* | Beiji nairovirus | HL9 | China/Great_Hinggan_Mountains | 2022 | PP473603 |
| *Nairoviridae* | *Norwavirus* | Beiji nairovirus | FRN-0475 | Japan | 2013 | LC795500 |
| *Nairoviridae* | *Norwavirus* | Mikuni nairovirus | NGN-0582 | Japan | 2013 | LC795516 |
| *Nairoviridae* | *Norwavirus* | Gakugsa tick virus | Rus/Ix persulcatus/Karelia/1/2018 | Russia | 2018 | MN542363 |
| *Nairoviridae* | *Norwavirus* | Beiji nairovirus | NE-SL4 | China/Great_Hinggan_Mountains | 2021 | ON408095 |
| *Nairoviridae* | *Norwavirus* | Beiji nairovirus | NE-TH4 | China/Great_Hinggan_Mountains | 2021 | ON408101 |
| *Nairoviridae* | *Norwavirus* | Beiji nairovirus | NE-TH3 | China/Great_Hinggan_Mountains | 2021 | ON408099 |
| *Nairoviridae* | *Norwavirus* | Beiji nairovirus | NE-YC4 | China/Xiaoxing'an_Mountains | 2021 | ON408103 |
| *Nairoviridae* | *Norwavirus* | Beiji nairovirus | NE-SL3 | China/Great_Hinggan_Mountains | 2021 | ON408097 |
| *Nairoviridae* | *Norwavirus* | Beiji nairovirus | H801 | China/Great_Hinggan_Mountains | 2018 | MW315108 |
| *Nairoviridae* | *Norwavirus* | Beiji nairovirus | HL5 | China/Great_Hinggan_Mountains | 2022 | PP473599 |
| *Nairoviridae* | *Norwavirus* | Beiji nairovirus | H39 | China/Great_Hinggan_Mountains | 2018 | MW315109 |
| *Nairoviridae* | *Norwavirus* | Beiji nairovirus | NE-YC3 | China/Xiaoxing'an_Mountains | 2021 | ON408105 |
| *Nairoviridae* | *Norwavirus* | Beiji nairovirus | YKS44 | China/Great_Hinggan_Mountains | 2018 | MN122080 |
| *Nairoviridae* | *Norwavirus* | Beiji nairovirus | OR4 | China/Great_Hinggan_Mountains | 2022 | PP473514 |
| *Nairoviridae* | *Norwavirus* | Beiji nairovirus | H160 | China/Great_Hinggan_Mountains | 2018 | NC 079010 |
| *Nairoviridae* | *Norwavirus* | Beiji nairovirus | HL2 | China/Great_Hinggan_Mountains | 2022 | PP473597 |
| *Nairoviridae* | *Norwavirus* | Beiji nairovirus | BSQG1734 | China/Changbai_Mountain | 2017 | OR148704 |
| *Nairoviridae* | *Norwavirus* | Beiji nairovirus | H56 | China/Great_Hinggan_Mountains | 2018 | MW315111 |
| *Nairoviridae* | *Norwavirus* | Beiji nairovirus | YBQG1715 | China/Changbai_Mountain | 2017 | OR148730 |
| *Nairoviridae* | *Norwavirus* | Beiji nairovirus | H59 | China/Great_Hinggan_Mountains | 2018 | MW315110 |
| *Nairoviridae* | *Norwavirus* | Beiji nairovirus | BSQG1720 | China/Changbai_Mountain | 2017 | OR148690 |
| *Nairoviridae* | *Norwavirus* | Beiji nairovirus | H1063 | China/Great_Hinggan_Mountains | 2018 | MW315112 |
| *Nairoviridae* | *Norwavirus* | Beiji nairovirus | THQG1701 | China/Changbai_Mountain | 2017 | OR148712 |
| *Nairoviridae* | *Norwavirus* | Beiji nairovirus | THQG1702 | China/Changbai_Mountain | 2017 | OR148714 |
| *Nairoviridae* | *Norwavirus* | Beiji nairovirus | HL4 | China/Great_Hinggan_Mountains | 2022 | PP473598 |
| *Nairoviridae* | *Norwavirus* | Beiji nairovirus | YBQG1712 | China/Changbai_Mountain | 2017 | OR148724 |
| *Nairoviridae* | *Norwavirus* | Beiji nairovirus | YBQG1739 | China/Changbai_Mountain | 2017 | OR148738 |
| *Nairoviridae* | *Norwavirus* | Beiji nairovirus | BSQG1730 | China/Changbai_Mountain | 2017 | OR148698 |
| *Nairoviridae* | *Norwavirus* | Beiji nairovirus | YBQG1716 | China/Changbai_Mountain | 2017 | OR148732 |
| *Nairoviridae* | *Norwavirus* | Beiji nairovirus | BSQG1738 | China/Changbai_Mountain | 2017 | OR148710 |
| *Nairoviridae* | *Norwavirus* | Beiji nairovirus | BSQG1735 | China/Changbai_Mountain | 2017 | OR148706 |
| *Nairoviridae* | *Norwavirus* | Beiji nairovirus | THQG1706 | China/Changbai_Mountain | 2017 | OR148716 |
| *Nairoviridae* | *Norwavirus* | Beiji nairovirus | BSQG1725 | China/Changbai_Mountain | 2017 | OR148696 |
| *Nairoviridae* | *Norwavirus* | Beiji nairovirus | YBQG1717 | China/Changbai_Mountain | 2017 | OR148734 |
| *Nairoviridae* | *Norwavirus* | Beiji nairovirus | THQG1746 | China/Changbai_Mountain | 2017 | OR148720 |
| *Nairoviridae* | *Norwavirus* | Beiji nairovirus | YBQG1713 | China/Changbai_Mountain | 2017 | OR148726 |
| *Nairoviridae* | *Norwavirus* | Beiji nairovirus | BSQG1732 | China/Changbai_Mountain | 2017 | OR148700 |
| *Nairoviridae* | *Norwavirus* | Beiji nairovirus | BSQG1723 | China/Changbai_Mountain | 2017 | OR148694 |
| *Nairoviridae* | *Norwavirus* | Beiji nairovirus | BSQG1737 | China/Changbai_Mountain | 2017 | OR148708 |
| *Nairoviridae* | *Norwavirus* | Beiji nairovirus | THQG1750 | China/Changbai_Mountain | 2017 | OR148722 |
| *Nairoviridae* | *Norwavirus* | Beiji nairovirus | BSQG1722 | China/Changbai_Mountain | 2017 | OR148692 |
| *Nairoviridae* | *Norwavirus* | Beiji nairovirus | YBQG1714 | China/Changbai_Mountain | 2017 | OR148728 |
| *Nairoviridae* | *Norwavirus* | Beiji nairovirus | THQG1707B | China/Changbai_Mountain | 2017 | OR148718 |
| *Nairoviridae* | *Norwavirus* | Beiji nairovirus | BSQG1733 | China/Changbai_Mountain | 2017 | OR148702 |
| *Nairoviridae* | *Norwavirus* | Beiji nairovirus | YBQG1718A | China/Changbai_Mountain | 2017 | OR148736 |
| *Nairoviridae* | *Norwavirus* | Beiji nairovirus | NE-DH3 | China/Changbai_Mountain | 2020 | ON408107 |
| *Nairoviridae* | *Norwavirus* | Mikuni nairovirus | FKS-1108 | Japan | 2014 | LC795518 |
| *Nairoviridae* | *Norwavirus* | Yichun nairovirus | ENW-1567 | Japan | 2015 | LC795520 |
| *Nairoviridae* | *Norwavirus* | Yichun nairovirus | NE-YC3 | China/Xiaoxing'an_Mountains | 2021 | ON408111 |
| *Nairoviridae* | *Norwavirus* | Yichun nairovirus | NE-FZ3 | China/Xiaoxing'an_Mountains | 2021 | ON408113 |
| *Nairoviridae* | *Norwavirus* | Yichun nairovirus | NE-YC4 | China/Xiaoxing'an_Mountains | 2021 | ON408109 |
| *Nairoviridae* | *Norwavirus* | Pustyn virus | IxRic1 | Russia | 2015 | KT007143 |
| *Nairoviridae* | *Norwavirus* | Norway nairovirus 1 | NOR/H3/Skanevik/2014 | Norway | 2014 | MF141041 |
| *Nairoviridae* | *Norwavirus* | Norway nairovirus 1 | NOR/A2/Bronnoya/2014 | Norway | 2014 | MF141043 |
| *Nairoviridae* | *Norwavirus* | Norway nairovirus 1 | NOR/B1V/Tofte/2014 | Norway | 2014 | MF141049 |
| *Nairoviridae* | *Norwavirus* | Grotenhout virus | Gierle-1 | Belgium | 2009 | NC 078264 |
| *Nairoviridae* | *Norwavirus* | Norway nairovirus 1 | NOR/S5/Kilen/2014 | Norway | 2014 | MF141045 |
| *Nairoviridae* | *Norwavirus* | Norway nairovirus 1 | NOR/MR4V/Kanestraum/2014 | Norway | 2014 | MF141047 |
| *Nairoviridae* | *Norwavirus* | Norwavirus sp. | sp. p80 | Bulgaria | 2021 | OQ269600 |
| *Nairoviridae* | *Norwavirus* | Norwavirus sp. | sp. p74 | Bulgaria | 2021 | OQ269596 |
| *Nairoviridae* | *Norwavirus* | Norwavirus sp. | sp. p67 | Bulgaria | 2021 | OQ269594 |
| *Nairoviridae* | *Norwavirus* | Pustyn virus | Rus/Ix ricinus/Moscow/2018 | Russia | 2018 | MN542361 |
| *Nairoviridae* | *Norwavirus* | Norwavirus sp. | p76 | Bulgaria | 2021 | OQ269598 |
| *Nairoviridae* | *Norwavirus* | Norwavirus sp. | p51 | Poland | 2021 | OQ269602 |
| *Nairoviridae* | *Orthonairovirus* | Peribunyaviridae sp. | 21HBT6 | China/Hubei | 2021 | PP471894 |
| *Nairoviridae* | *Orthonairovirus* | Peribunyaviridae sp. | PeV/MCYTH4 | China/Hubei | 2019 | MW721860 |
| *Nairoviridae* | *Orthonairovirus* | Peribunyaviridae sp. | PeV/LTBMH5 | China/Hubei | 2019 | MW721857 |
| *Nairoviridae* | *Orthonairovirus* | Peribunyaviridae sp. | PeV/MCSH2 | China/Hubei | 2019 | MW721858 |
| *Nairoviridae* | *Orthonairovirus* | Peribunyaviridae sp. | PeV/MCSH4 | China/Hubei | 2019 | MW721859 |
| *Nairoviridae* | *Orthonairovirus* | Peribunyaviridae sp. | PeV/SZYD10 | China/Hubei | 2019 | MW721861 |
| *Nairoviridae* | *Orthonairovirus* | Peribunyaviridae sp. | PeV/SZYD9 | China/Hubei | 2019 | MW721863 |
| *Nairoviridae* | *Orthonairovirus* | Peribunyaviridae sp. | PeV/SZYD14 | China/Hubei | 2019 | MW721862 |
| *Nairoviridae* | *Orthonairovirus* | Peribunyaviridae sp. | PeV/MCSH4 | China/Hubei | 2019 | MZ964995 |
| *Nairoviridae* | *Orthonairovirus* | Peribunyaviridae sp. | PeV/MCSH2 | China/Hubei | 2019 | MZ964994 |
| *Nairoviridae* | *Orthonairovirus* | Henan tick virus | CLCM-083 | China/Shandong | 2021 | OR114971 |
| *Nairoviridae* | *Orthonairovirus* | Henan tick virus | CLCM-132 | China/Shandong | 2020 | OR114968 |
| *Nairoviridae* | *Orthonairovirus* | Henan tick virus | CLCM-130 | China/Shandong | 2020 | OR114966 |
| *Nairoviridae* | *Orthonairovirus* | Henan tick virus | CLCM-099 | China/Shandong | 2020 | OR114964 |
| *Nairoviridae* | *Orthonairovirus* | Henan tick virus | CLCM-097 | China/Shandong | 2020 | OR114960 |
| *Nairoviridae* | *Orthonairovirus* | Henan tick virus | CLCM-096 | China/Shandong | 2020 | OR114957 |
| *Nairoviridae* | *Orthonairovirus* | Henan tick virus | CLCM-090 | China/Shandong | 2021 | OR114954 |
| *Nairoviridae* | *Orthonairovirus* | Henan tick virus | CLCM-082 | China/Shandong | 2021 | OR114952 |
| *Nairoviridae* | *Orthonairovirus* | Henan tick virus | CLCM-058 | China/Shandong | 2022 | OR114949 |
| *Nairoviridae* | *Orthonairovirus* | Henan tick virus | CLCM-052 | China/Shandong | 2022 | OR114945 |
| *Nairoviridae* | *Orthonairovirus* | Henan tick virus | CLCM-072 | China/Shandong | 2018 | OR114941 |
| *Nairoviridae* | *Orthonairovirus* | Henan tick virus | CLCM-060 | China/Shandong | 2021 | OR114938 |
| *Nairoviridae* | *Orthonairovirus* | Huangpi Tick Virus 1 | 21HBT2 | China/Hubei | 2021 | PP471899 |
| *Nairoviridae* | *Orthonairovirus* | Huangpi Tick Virus 1 | HTV1/SZYD9 | China/Hubei | 2019 | MW721869 |
| *Nairoviridae* | *Orthonairovirus* | Huangpi Tick Virus 1 | HTV1/SZYD12 | China/Hubei | 2019 | MW721868 |
| *Nairoviridae* | *Orthonairovirus* | Huangpi Tick Virus 1 | HTV1/SZYD09 | China/Hubei | 2019 | MZ965001 |
| *Nairoviridae* | *Orthonairovirus* | Huangpi Tick Virus 1 | HTV1/SZYD12 | China/Hubei | 2019 | MZ965000 |
| *Nairoviridae* | *Orthonairovirus* | Jian nariovirus | NE-JA | China/Changbai_Mountain | 2021 | ON408090 |
| *Nairoviridae* | *Orthonairovirus* | Jian nariovirus | NE-DH1 | China/Changbai_Mountain | 2020 | ON408093 |
| *Nairoviridae* | *Orthonairovirus* | Jian nariovirus | NE-YC2 | China/Xiaoxing'an_Mountains | 2021 | ON408087 |
| *Nairoviridae* | *Orthonairovirus* | Jian nariovirus | NE-MDJ1 | China/Changbai_Mountain | 2021 | ON408084 |
| *Nairoviridae* | *Orthonairovirus* | Pangolin orthonairovirus | BIME1 | China/Beijing | 2018 | ON024089 |
| *Nairoviridae* | *Orthonairovirus* | Shanxi tick virus 2 | CLCM-119 | China/Shandong | 2019 | OR114983 |
| *Nairoviridae* | *Orthonairovirus* | Shanxi tick virus 1 | CLCM-100 | China/Shandong | 2020 | OR114979 |
| *Nairoviridae* | *Orthonairovirus* | Shanxi tick virus 0 | CLCM-037 | China/Shandong | 2022 | OR114976 |
| *Nairoviridae* | *Orthonairovirus* | Shanxi tick virus 1 | CLCM-065 | China/Shandong | 2021 | OR114974 |
| *Nairoviridae* | *Orthonairovirus* | Shanxi tick virus 2 | 21HBT4 | China/Hubei | 2021 | PP471907 |
| *Nairoviridae* | *Orthonairovirus* | Antu virus | YB tick 2021 24 | China/Changbai_Mountain | 2021 | OQ207701 |
| *Nairoviridae* | *Orthonairovirus* | Songling virus | HLJ1202 | China/Xiaoxing'an_Mountains | 2002 | NC 079002 |
| *Nairoviridae* | *Orthonairovirus* | Songling virus | NE-TH2 | China/Great_Hinggan_Mountains | 2021 | ON408081 |
| *Nairoviridae* | *Orthonairovirus* | Songling virus | NE-TH1 | China/Great_Hinggan_Mountains | 2021 | ON408078 |
| *Nairoviridae* | *Orthonairovirus* | Songling virus | YC585 | China/Xiaoxing'an_Mountains | 2005 | MT328780 |
| *Nairoviridae* | *Orthonairovirus* | Songling virus | HLJ1202 | China/Xiaoxing'an_Mountains | 2002 | MT328777 |
| *Nairoviridae* | *Orthonairovirus* | Yezo virus | BT-2135 | Japan | 2021 | LC790679 |
| *Nairoviridae* | *Orthonairovirus* | Yezo virus | BT-1968 | Japan | 2020 | LC790676 |
| *Nairoviridae* | *Orthonairovirus* | Yezo virus | BT-2155 | Japan | 2021 | LC790682 |
| *Nairoviridae* | *Orthonairovirus* | Yezo virus | BT-1844 | Japan | 2020 | LC735733 |
| *Nairoviridae* | *Orthonairovirus* | Yezo virus | BT-1826 | Japan | 2020 | LC735730 |
| *Nairoviridae* | *Orthonairovirus* | Yezo virus | BT-1821 | Japan | 2020 | LC735727 |
| *Nairoviridae* | *Orthonairovirus* | Yezo virus | HH003-2020 | Japan | 2020 | NC 079100 |
| *Nairoviridae* | *Orthonairovirus* | Yezo virus | T-JL01 | China/Jilin | 2020 | ON563282 |
| *Nairoviridae* | *Orthonairovirus* | Yezo virus | YBQG1739 | China/Jilin | 2017 | OR148888 |
| *Nairoviridae* | *Orthonairovirus* | Yezo virus | BT-1864 | Japan | 2020 | LC735736 |
| *Nairoviridae* | *Orthonairovirus* | Yezo virus | HH008-2017 | Japan | 2017 | LC628644 |
| *Nairoviridae* | *Orthonairovirus* | Yezo virus | YBQG1712 | China/Jilin | 2017 | OR148885 |
| *Nairoviridae* | *Orthonairovirus* | Yezo virus | H-IM01 | China/Inner_Mongolia | 2018 | ON563285 |
| *Nairoviridae* | *Orthonairovirus* | Yezo virus | T-IM01 | China/Inner_Mongolia | 2021 | ON563279 |
| *Nairoviridae* | *Orthonairovirus* | Yezo virus | T-HLJ01 | China/Heilongjiang | 2021 | ON563270 |
| *Nairoviridae* | *Orthonairovirus* | Yezo virus | T-HLJ03 | China/Heilongjiang | 2021 | ON563276 |
| *Nairoviridae* | *Orthonairovirus* | Yezo virus | THQG1707B | China/Jilin | 2017 | OR148882 |
| *Nairoviridae* | *Orthonairovirus* | Yezo virus | HH001-2019 | Japan | 2019 | LC621354 |
| *Nairoviridae* | *Orthonairovirus* | Yezo virus | HH007-2016 | Japan | 2016 | LC628643 |
| *Nairoviridae* | *Orthonairovirus* | Yezo virus | HH011-2020 | Japan | 2020 | LC621360 |
| *Nairoviridae* | *Orthonairovirus* | Yezo virus | T-HLJ02 | China/Heilongjiang | 2021 | ON563273 |
| *Nairoviridae* | *Orthonairovirus* | Yezo virus | HH009-2017 | Japan | 2017 | LC628645 |
| *Nairoviridae* | *Orthonairovirus* | Sulina virus | b30 | Latvia | 2022 | PP260004 |
| *Nairoviridae* | *Orthonairovirus* | Sulina virus | IxriSL16-01 | Romania | 2016 | NC 078998 |
| *Nairoviridae* | *Orthonairovirus* | Sulina virus | IxriSL16-02 | Romania | 2016 | MT263287 |
| *Nairoviridae* | *Orthonairovirus* | Nairobi sheep disease virus | NS | China/Changbai_Mountain | 2022 | OQ716536 |
| *Nairoviridae* | *Orthonairovirus* | Nairobi sheep disease virus | SDWL08/2020/China/Shandong | China/Shandong | 2020 | OR495604 |
| *Nairoviridae* | *Orthonairovirus* | Nairobi sheep disease virus | SDWL16/2020/China/Shandong | China/Shandong | 2020 | OR495601 |
| *Nairoviridae* | *Orthonairovirus* | Nairobi sheep disease virus | SDWL07/2020/China/Shandong | China/Shandong | 2020 | OR495598 |
| *Nairoviridae* | *Orthonairovirus* | Nairobi sheep disease virus | Jilin | China/Changbai_Mountain | 2013 | NC 034386 |
| *Nairoviridae* | *Orthonairovirus* | Nairobi sheep disease virus | SZWH2 | China/Hubei | 2019 | MW721872 |
| *Nairoviridae* | *Orthonairovirus* | Nairobi sheep disease virus | Hubei | China/Hubei | 2016 | MH791451 |
| *Nairoviridae* | *Orthonairovirus* | Meihua Mountain virus | LJ01 | China/Fujian | 2020 | ON184086 |
| *Nairoviridae* | *Orthonairovirus* | Tofla virus | Nag-Hfor-2014 | Japan | 2014 | LC030113 |
| *Nairoviridae* | *Orthonairovirus* | Tofla virus |  | Japan | 2013 | NC 029122 |
| *Nairoviridae* | *Orthonairovirus* | Wetland virus | PS1 | China/Great_Hinggan_Mountains | 2019 | OR860396 |
| *Nairoviridae* | *Orthonairovirus* | Wetland virus | HC1 | China/Great_Hinggan_Mountains | 2021 | OR860397 |
| *Nairoviridae* | *Orthonairovirus* | Wetland virus | HC2 | China/Great_Hinggan_Mountains | 2023 | OR860398 |
| *Nairoviridae* | *Orthonairovirus* | Wetland virus | T6D12 | China/Great_Hinggan_Mountains | 2019 | OR860399 |
| *Nairoviridae* | *Orthonairovirus* | Wetland virus | T3G2 | China/Great_Hinggan_Mountains | 2021 | OR860400 |
| *Nairoviridae* | *Orthonairovirus* | Wetland virus | T8A1 | China/Great_Hinggan_Mountains | 2023 | OR860401 |
| *Nairoviridae* | *Orthonairovirus* | Dadong virus YB tick 2023 134 | YB tick 2023 134 | China/Changbai_Mountain | 2023 |  |

**Supplementary Table 8.** Sampling information of sequencing libraries by collection sites and species.

| **Sequencing libraries** | **Region** | **Tick species (number analyzed)** | | | | | | | | |
| --- | --- | --- | --- | --- | --- | --- | --- | --- | --- | --- |
|  |  | ***Ixodes persulcatus*** | | ***Haemaphysalis concinna*** | | ***Haemaphysalis longicornis*** | | ***Dermacentor nuttalli*** | | **Total** |
|  |  | **Female** | **Male** | **Female** | **Male** | **Female** | **Male** | **Female** | **Male** |  |
| 22QUAN | Benxi | 37 | 47 | 0 | 0 | 0 | 0 | 0 | 0 | **118** |
|  | xinlin | 10 | 24 | 0 | 0 | 0 | 0 | 0 | 0 |  |
| NEIQUAN | Yakeshi | 30 | 7 | 0 | 0 | 0 | 0 | 0 | 0 | **93** |
|  | Genhe | 30 | 26 | 0 | 0 | 0 | 0 | 0 | 0 |  |
| LVQUAN | Yakeshi | 120 | 75 | 0 | 0 | 0 | 0 | 0 | 0 | **239** |
|  | Argun | 3 | 2 | 0 | 0 | 0 | 0 | 0 | 0 |  |
|  | Oroqen | 19 | 20 | 0 | 0 | 0 | 0 | 0 | 0 |  |
| XQ QUAN | New Barag West Banner | 50 | 130 | 0 | 0 | 0 | 0 | 0 | 0 | **180** |
| 23QUANF | Dandong | 6 | 0 | 0 | 0 | 0 | 0 | 0 | 0 | **188** |
|  | Ji'an | 18 | 0 | 0 | 0 | 0 | 0 | 0 | 0 |  |
|  | Linjiang | 101 | 0 | 0 | 0 | 0 | 0 | 0 | 0 |  |
|  | Hunchun | 33 | 0 | 0 | 0 | 0 | 0 | 0 | 0 |  |
|  | Dongning | 30 | 0 | 0 | 0 | 0 | 0 | 0 | 0 |  |
| 23QUANM | Ji'an | 0 | 18 | 0 | 0 | 0 | 0 | 0 | 0 | **283** |
|  | Linjiang | 0 | 194 | 0 | 0 | 0 | 0 | 0 | 0 |  |
|  | Hunchun | 0 | 52 | 0 | 0 | 0 | 0 | 0 | 0 |  |
|  | Dongning | 0 | 19 | 0 | 0 | 0 | 0 | 0 | 0 |  |
| QUAN | Antu | 168 | 220 | 0 | 0 | 0 | 0 | 0 | 0 | **394** |
|  | Tumen | 2 | 4 | 0 | 0 | 0 | 0 | 0 | 0 |  |
| 725QUAN | Wangqing | 78 | 62 | 0 | 0 | 0 | 0 | 0 | 0 | **140** |
| 927GEXUE | Yakeshi | 0 | 0 | 96 | | 0 | 0 | 161 | 121 | **378** |
| JHGE | Jiaohe | 0 | 0 | 0 | 0 | 0 | 0 | 79 | 60 | **139** |
| LVGE | Yakeshi | 0 | 0 | 0 | 0 | 0 | 0 | 40 | 28 | **188** |
|  | Argun | 0 | 0 | 0 | 0 | 0 | 0 | 37 | 40 |  |
|  | Oroqen | 0 | 0 | 0 | 0 | 0 | 0 | 33 | 10 |  |
| 927GE | Yakeshi | 0 | 0 | 0 | 0 | 0 | 0 | 180 | 118 | **298** |
| 23GE | Tongliao | 0 | 0 | 0 | 0 | 0 | 0 | 29 | 31 | **180** |
|  | chifeng | 0 | 0 | 0 | 0 | 0 | 0 | 71 | 47 |  |
|  | New Barag West Banner | 0 | 0 | 0 | 0 | 0 | 0 | 2 | 0 |  |
| GE1 | Antu | 0 | 0 | 0 | 0 | 0 | 0 | 278 | 320 | **617** |
|  | Tumen | 0 | 0 | 0 | 0 | 0 | 0 | 3 | 8 |  |
|  | Hunchun | 0 | 0 | 0 | 0 | 0 | 0 | 8 | 0 |  |
| GE2 | Antu | 0 | 0 | 0 | 0 | 0 | 0 | 0 | 599 | **599** |
| XueXiong | Tahe | 0 | 0 | 113 | | 0 | 0 | 0 | 0 | **113** |
| LNXUE | Benxi | 0 | 0 | 156 | | 174 | | 0 | 0 | **330** |
| 22XUE | Yakeshi | 0 | 0 | 131 | | 0 | 0 | 0 | 0 | **131** |
| LVXUE | Yakeshi | 0 | 0 | 80 | | 0 | 0 | 0 | 0 | **212** |
|  | Oroqen | 0 | 0 | 42 | | 0 | 0 | 0 | 0 |  |
|  | Genhe | 0 | 0 | 90 | | 0 | 0 | 0 | 0 |  |
| 23SQXUE | Ji'an | 0 | 0 | 92 | | 0 | 0 | 0 | 0 | **348** |
|  | Linjiang | 0 | 0 | 51 | | 0 | 0 | 0 | 0 |  |
|  | Hunchun | 0 | 0 | 178 | | 0 | 0 | 0 | 0 |  |
|  | xinlin | 0 | 0 | 27 | | 0 | 0 | 0 | 0 |  |
| 23CJXUE | Dandong | 0 | 0 | 0 | 0 | 237 | | 0 | 0 | **595** |
|  | Ji'an | 0 | 0 | 0 | 0 | 68 | | 0 | 0 |  |
|  | Linjiang | 0 | 0 | 0 | 0 | 61 | | 0 | 0 |  |
|  | Hunchun | 0 | 0 | 0 | 0 | 174 | | 0 | 0 |  |
|  | Dongning | 0 | 0 | 0 | 0 | 20 | | 0 | 0 |  |
|  | Antu | 0 | 0 | 0 | 0 | 35 | | 0 | 0 |  |
| XUE | Antu | 0 | 0 | 27 | | 0 | 0 | 0 | 0 | **280** |
|  | Hunchun | 0 | 0 | 253 | | 0 | 0 | 0 | 0 |  |
| WGQXUE | Wangqing | 0 | 0 | 387 | | 52 | | 0 | 0 | **439** |
| CJSQ | Helong | 0 | 0 | 79 | | 63 | | 0 | 0 | **266** |
|  | Tumen | 0 | 0 | 76 | | 48 | | 0 | 0 |  |
| 725SQCJ | Hunchun | 0 | 0 | 214 | | 77 | | 0 | 0 | **291** |
| 718SQ | Hunchun | 0 | 0 | 177 | | 180 | | 0 | 0 | **357** |
| 718CJ | Hunchun | 0 | 0 | 160 | | 140 | | 0 | 0 | **300** |

**Supplementary Table 9.** The information of primers used in this study.

| **Virus** | **Primer** | **Position (bp)^*^** | **Sequence (5'→3')** | **Polarity** | **Amplicon (bp)** |
| --- | --- | --- | --- | --- | --- |
| **Beiji virus** | | | | | |
|  | Detection |  |  |  |  |
|  | BJNV-F | 7258 | GGAAGGAGGATATTTTCTACC | + | 110 |
|  | BJNV-R | 7338 | CCAGGTATATGTCAGTCAATAG | - |  |
|  | BJNV-P | 7281 | AGCACAACAATCAACTCACTAAGCCT | + |  |
|  | Genome amplification | | | | |
| Segment L | 1 BJNV L F | 1 | CCCCGATTTCCAAAGACTTTTC | + | 268 |
|  | 1 BJNV L R2 | 247 | GTCAATGTACAACCACCACGTA | - |  |
|  | 1 BJNV L R1 | 452 | GGAGAAATACATCCCCTTCG | - |  |
|  | 2 BJNV L F | 193 | CAGGAATGGCATCTAGTAA | + | 843 |
|  | 2 BJNV L R2 | 1014 | TTGGTCCAGTGAGGTCTTCCGA | - |  |
|  | 2 BJNV L R1 | 1077 | CTGCAACCTCGGCTAAGGCTTT | - |  |
|  | 3 BJNV L F1 | 738 | TCTCTCATGGCTTCAACTGT | + | 792 |
|  | 3 BJNV L F2 | 851 | CRCTCATGATAGRCACACCA | + |  |
|  | 3 BJNV L R | 1622 | TCCAGGCACTTATCAACCTGT | - |  |
|  | 4 BJNV L F | 1561 | CCCCAAAGTCAGTACAAATCTCC | + | 997 |
|  | 4 BJNV L R2 | 2539 | CTAARGRGTTCTCCAACGC | + |  |
|  | 4 BJNV L R1 | 2617 | CTGTATCTTTRGTTGRYGTTT | - |  |
|  | 5 BJNV L F | 2406 | AAGAMAGACCAAAACACCG | + | 912 |
|  | 5 BJNV L R2 | 3292 | GCTATAAATAGAAATRTCAGCTGCAT | - |  |
|  | 5 BJNV L R1 | 3475 | TGGCAAATAACACCAAAAYTCAG | - |  |
|  | 6 BJNV L F1 | 3187 | TWAACATGCTAGTCCGGTTT | + | 996 |
|  | 6 BJNV L F2 | 3257 | TCACTTAGGAGCTGARCAGAC | + |  |
|  | 6 BJNV L R | 4232 | AYCATAGGCATYARTTCCACT | - |  |
|  | 7 BJNV L F | 4189 | TRCAYAARAGGTTAATGACA | + | 901 |
|  | 7 BJNV L R2 | 5069 | ACATCTGTGGCAATTATCAGG | - |  |
|  | 7 BJNV L R1 | 5206 | ATAARCCACTAACTATTCGAT | - |  |
|  | 8 BJNV L F | 5065 | CAACCCTGATAATTGCCACA | + | 944 |
|  | 8 BJNV L R2 | 5988 | AATAACCTTACGTGGAACACA | - |  |
|  | 8 BJNV L R1 | 6145 | TACTATTRGCCTGATCATCCA | - |  |
|  | 9 BJNV L F | 5982 | AAAATCYGTGYYCCACGTA | + | 864 |
|  | 9 BJNV L R2 | 6824 | TTTATCATAGTTTAACGTGCCT | - |  |
|  | 9 BJNV L R1 | 7095 | GCCCATTATTGTGGTRAACCG | - |  |
|  | 10 BJNV L F | 6786 | ATACAGAACTACAAGTGCCTA | + | 894 |
|  | 10 BJNV L R2 | 7660 | CCAGCTTCGAACACTTGAGA | - |  |
|  | 10 BJNV L R1 | 7837 | CCCCATAGTGTAACTTGGAT | - |  |
|  | 11 BJNV L F | 7549 | TAAAGCTGGATCAGTTACCTC | + | 922 |
|  | 11 BJNV L R2 | 8450 | TGGACGGTTTTAAGATACTCA | - |  |
|  | 11 BJNV L R1 | 8534 | GGTTTACTCATTTCTTCYGTC | - |  |
|  | 12 BJNV L F | 8217 | GCACGRTCTTTAAGACAAGTCAC | + | 849 |
|  | 12 BJNV L R2 | 9044 | TAAAACACCATCCAGATCRGTA | - |  |
|  | 12 BJNV L R1 | 9195 | TACTCCTTGGTAGCCGTTC | - |  |
|  | 13 BJNV L F1 | 9000 | AACCCACTGTCATATGTGC | + | 1143 |
|  | 13 BJNV L F2 | 9027 | AGGCTATCTAACATCAATACCG | + |  |
|  | 13 BJNV L R | 10149 | CCARTTTGTYTCATAWGYCTG | - |  |
|  | 14 BJNV L F | 10087 | GCTCTTCTGATGATTATGCG | + | 768 |
|  | 14 BJNV L R2 | 10897 | TGCCCATCATAACACTGTC | - |  |
|  | 14 BJNV L R1 | 11013 | CRAATCGGAGGTCTCCTCT | - |  |
|  | 15 BJNV L F | 10818 | CATGTTGCTGATAAAGTCCG | + | 961 |
|  | 15 BJNV L R2 | 11757 | TGGCATACTTTTCACATAGTCT | - |  |
|  | 15 BJNV L R1 | 11808 | TTCAGCCTGTTATGGTCACT | - |  |
|  | 16 BJNV L F | 11654 | ATGCCTGGTACTTGATGACA | + | 1006 |
|  | 16 BJNV L R2 | 12637 | TGTTGAAAAGGAAATACGACCTC | - |  |
|  | 16 BJNV L R1 | 12674 | CCAATGTAAARTTGCCTGAGCTA | - |  |
|  | 17 BJNV L F1 | 12482 | TTCTCTGGTTGCACTGTTCG | + | 937 |
|  | 17 BJNV L F2 | 12586 | CTATGTTTGCYCTAAATGCAC | + |  |
|  | 17 BJNV L R | 13502 | TTGAGTATGTCCCTTAGCCTA | - |  |
|  | 18 BJNV L F1 | 13308 | TCTGGYTTTAAGTCYTACTCA | + | 878 |
|  | 18 BJNV L F2 | 13192 | AACTAAGRAAAGAAGGTGCTC | + |  |
|  | 18 BJNV L R | 14047 | ATCACTTTTACGCTTATATCCCA | - |  |
|  | 19 BJNV L F | 14008 | GGCTCTATTTTCTTAGGGTT | + | 735 |
|  | 19 BJNV L R2 | 14723 | GGATTAATGTCCCCTACTGA | - |  |
|  | 19 BJNV L R1 | 14774 | AACATTTCTCTATARTCCRTTC | - |  |
| Segment S | 1 BJNV S F | 1 | TCTCGAAAATATCTAACCCCCG | + | 566 |
|  | 1 BJNV S R2 | 547 | RCCYTCTTGRDGTTTCCTCT | - |  |
|  | 1 BJNV S R1 | 648 | AAACCCAACAGAAGCAGGAG | - |  |
|  | 2 BJNV S F | 251 | CACWGTTTATCAYYTGTCAGYY | + | 740 |
|  | 2 BJNV S R2 | 976 | AGTARGTTCKGTCATTCACA | - |  |
|  | 2 BJNV S R1 | 1011 | AGTCTAGATCAGGAAGCCCAT | - |  |
|  | 3 BJNV S F1 | 841 | GRCCWCTRCGRAACCCAGT | + | 939 |
|  | 3 BJNV S F2 | 892 | CWAYCACAATGCCTGTTGCCT | + |  |
|  | 3 BJNV S R | 1815 | CTGTCACCAGYTCTGYYTCCY | - |  |
|  | 4 BJNV S F1 | 1705 | CYAACTTCAGGGAACACCAC | + | 784 |
|  | 4 BJNV S F2 | 1738 | GTGATGASRCCTCTGCTGAC | + |  |
|  | 4 BJNV S R | 2507 | CCTCTTCCAAARGGACGGAT | - |  |
|  | 5 BJNV S F | 2423 | ATTTGCTGGMCTAGARCCGTA | + | 914 |
|  | 5 BJNV S R2 | 3321 | CCTGACTWAGAACAGTGGCTA | - |  |
|  | 5 BJNV S R1 | 3368 | CACACTGACACTAAAGGGGAT | - |  |
|  | 6 BJNV S F1 | 3235 | ACATAACAGTTGACCATGSCTT | + | 400 |
|  | 6 BJNV S F2 | 3317 | AGCCACTGTTCTWAGTCARGA | + |  |
|  | 6 BJNV S R | 3701 | CTAAGATGTCATACCCCCGAA | - |  |
| **Songling virus** |  |  |  |  |  |
|  | Detection |  |  |  |  |
|  | SGLV-F | 4352 | CAGCAYTCTAGATGYGTAGA | + | 163 |
|  | SGLV-R | 4495 | CACAGTACTCCACATAACTC | - |  |
|  | SGLV-P | 4375 | RGTYTACGATGCCAACCACAGC | + |  |
|  | Genome amplification | | | | |
| Segment L | 1 SGLV L F | 1 | TCTCAAAGATATATATCCTGCAC | + | 556 |
|  | 1 SGLV L R2 | 536 | GTTCACYTCYGGCCTGTCACC | - |  |
|  | 1 SGLV L R1 | 762 | CCTGACATCYTYTCTACTGGCT | - |  |
|  | 2 SGLV L F1 | 356 | TCTTTRGAAMGGTTYTATGGC | + | 911 |
|  | 2 SGLV L F2 | 445 | TCTYAACAAGGCTTTTGGMAT | + |  |
|  | 2 SGLV L R | 1335 | GWGARAGCTCCTTRTYCTCCT | - |  |
|  | 3 SGLV L F1 | 1211 | AATCTAYAYCTCAATGGGGTC | + | 851 |
|  | 3 SGLV L F2 | 2107 | ATCCATGGCAGTACAAACAT | + |  |
|  | 3 SGLV L R | 2107 | ATGTTTGTACTGCCATGGAT | - |  |
|  | 4 SGLV L F1 | 1958 | GCCAGGATGCTCTTYACTCG | + | 958 |
|  | 4 SGLV L F2 | 1992 | ACACTCAYACAACYGCWTGGT | + |  |
|  | 4 SGLV L R | 2930 | AGGCTYTCCATTCTYTCAGG | - |  |
|  | 5 SGLV L F1 | 2798 | AGCACTCAGAAAATAAGRGCAGT | + | 959 |
|  | 5 SGLV L F2 | 2873 | ACAACCTGGGGAGAGAACAAA | + |  |
|  | 5 SGLV L R | 3812 | CGCTCRGACCTAGTTGCYTC | - |  |
|  | 6 SGLV L F1 | 3652 | CAAGACCTAYTCTGYYGTCGTY | + | 986 |
|  | 6 SGLV L F2 | 3697 | RCTCACYACCCATAGGCAGA | + |  |
|  | 6 SGLV L R | 4662 | TCTAGACCTCTGGCATTCCTY | - |  |
|  | 7 SGLV L F | 4506 | AGTACTGTGCCAAGTCAGGY | + | 848 |
|  | 7 SGLV L R2 | 5335 | CCCCAGTCCAAGGTCTCTG | - |  |
|  | 7 SGLV L R1 | 5457 | TTTGTCGTGCAACTTCGGYCA | - |  |
|  | 8 SGLV L F | 5201 | AARACAAGGTCCTTYAGCTCA | + | 912 |
|  | 8 SGLV L R2 | 6092 | CATCCAGATYGYATATAGCTC | - |  |
|  | 8 SGLV L R1 | 6268 | GRGCCTTKACTTCWATCTCC | - |  |
|  | 9 SGLV L F1 | 5927 | ATGCCTGCACTYAGATACTCY | + | 962 |
|  | 9 SGLV L F2 | 5963 | AGTGACCAYCCYAAACTCTC | + |  |
|  | 9 SGLV L R | 6904 | AAYCTATGCCCTGAATTGAGT | - |  |
|  | 10 SGLV L F | 6840 | ARATGATAAAGTTRGTCAGGAGC | + | 847 |
|  | 10 SGLV L R2 | 7665 | TAGTCATCAGAACTGCCCGCAT | - |  |
|  | 10 SGLV L R1 | 7791 | GGGTTGCAAGRGGCTGTCA | - |  |
|  | 11 SGLV L F | 7505 | TCCAAAGGAAAGCTGGCCCTA | + | 824 |
|  | 11 SGLV L R2 | 8308 | GAGCTGTCATCAATTGTCCCC | - |  |
|  | 11 SGLV L R1 | 8416 | ATGTGCTRGTCCTYAAATACTCA | - |  |
|  | 12 SGLV L F | 8115 | CATTTGGCCGYCTYTAYYTACCT | + | 860 |
|  | 12 SGLV L R2 | 8953 | TTTAATTCCACCRGCAAGACCA | - |  |
|  | 12 SGLV L R1 | 9092 | CCGTCYAGAAGCTTTCCTGT | - |  |
|  | 13 SGLV L F | 8747 | CTAATGCCAGAAAAGCTCAGA | + | 822 |
|  | 13 SGLV L R2 | 9546 | 69 | - |  |
|  | 13 SGLV L R1 | 9652 | GCATGGCTRACCACATCYTCC | - |  |
|  | 14 SGLV L F1 | 9306 | TCAGAGCACTTTCYATCCAGT | + | 905 |
|  | 14 SGLV L F2 | 9464 | GCRTTRGTTATAGGGTTCCAT | + |  |
|  | 14 SGLV L R | 10348 | GTGACTATGTCTGTTCGCATC | - |  |
|  | 15 SGLV L F1 | 10137 | THCTAATTGCCAAGAAGCCAT | + | 767 |
|  | 15 SGLV LF2 | 10203 | CTAGAACAGGHAACTTCACTCT | + |  |
|  | 15 SGLV LR | 10950 | TGTRCTGTCYCTRATCTGTC | - |  |
|  | 16 SGLV L F | 10799 | GCCTTGCTCCATTATTACCTT | + | 836 |
|  | 16 SGLV L R2 | 11616 | AGGCACTTYCGYAACTCCA | - |  |
|  | 16 SGLV L R1 | 11720 | TGAGATTCCRCAGAACAGCTT | - |  |
|  | 17 SGLV L F1 | 11466 | CAGTACGACAGCCACAAYATA | + | 413 |
|  | 17 SGLV L F2 | 11581 | AAAARGCACTRTGCGACA | + |  |
|  | 17 SGLV L R | 11975 | ACGTGAATCTTAACTAAAA | - |  |
| Segment M | 1 SGLV M F | 1 | CGATCACTCCCAAACATTACAT | + | 505 |
|  | 1 SGLV M R2 | 490 | GAAACAGCWCCRGTTCCCCAGGA | - |  |
|  | 1 SGLV M R1 | 556 | TTGAAGAGGAGCATYCCCAT | - |  |
|  | 2 SGLV M F1 | 292 | CCAGTGTGCCKCTCCRACA | + | 719 |
|  | 2 SGLV M F2 | 371 | AGCCMAGCCYACCRYCCAG | + |  |
|  | 2 SGLV M R | 1069 | CTTRGAYAGCTCTARCCGRTT | - |  |
|  | 3 SGLV M F | 834 | AAGRGTCATCAGGAACCAGC | + | 706 |
|  | 3 SGLV M R2 | 1519 | GCCAGGTGTCTTYACATCCAT | - |  |
|  | 3 SGLV M R1 | 1714 | CTACAGACCCARAGAGGCAA | - |  |
|  | 4 SGLV M F | 1414 | CTGTCCAAAGTYCAGTGYCCDA | + | 754 |
|  | 4 SGLV M R2 | 2148 | CAAAGATBCCCAGRAACCAT | - |  |
|  | 4 SGLV M R1 | 2209 | GTCCTCYTCTATGGCAGTGTC | - |  |
|  | 5 SGLV M F1 | 1966 | TGCCCCTTYTGYCAGACCA | + | 772 |
|  | 5 SGLV M F2 | 1986 | TGCYAGYAAYGAYAAGCTC | + |  |
|  | 5 SGLV M R | 2738 | CTGYCTRTCYCCTGTGGCAT | - |  |
|  | 6 SGLV M F | 2583 | CCACATAACAGTMAACGGCAAG | + | 759 |
|  | 6 SGLV M R2 | 3320 | GACATCCAYGAGTTWGTAGCAT | - |  |
|  | 6 SGLV M R1 | 3519 | TGGCCTYCCTTTCAGYAGR | - |  |
|  | 7 SGLV M F | 3239 | ACCTWCAGATTTACGACGTG | + | 817 |
|  | 7 SGLV M R2 | 4037 | CTGCCRAATGCYCTSGTYA | - |  |
|  | 7 SGLV M R1 | 4060 | GTATAGTGACRAYGCCRAGCCA | - |  |
|  | 8 SGLV M F1 | 3814 | TTAGGTTYTATGCTGCAGTG | + | 647 |
|  | 8 SGLV M F2 | 3837 | AAAGAAAAAGTGTGTTTTGA | + |  |
|  | 8 SGLV M R | 4465 | GACGGTGYGCTGCWGYTGG | - |  |
| Segment S | 1 SGLV S F | 1 | TGTCAAAGAAAAACGTGCTGCA | + | 382 |
|  | 1 SGLV S R2 | 360 | TCCAGGCGCACTCATACACA | - |  |
|  | 1 SGLV S R1 | 453 | AGGCTTTCGTACTCCTTGGAC | - |  |
|  | 2 SGLV S F | 275 | YAGCCTYSACATCAAGTCTGC | + | 761 |
|  | 2 SGLV S R2 | 1017 | CAAACCCGCTGTTCTTCGAG | - |  |
|  | 2 SGLV S R1 | 1087 | GGTYTGACTCCTGCCTTCCA | - |  |
|  | 3 SGLV S F | 958 | CCTTCTTCAARAGCCTGTGGT | + | 750 |
|  | 3 SGLV S R2 | 1690 | GTCCRTCCCYTGTTGCKCT | - |  |
|  | 3 SGLV S R1 | 1709 | CCTGCCATCTGCCCKTTCG | - |  |
|  | 4 SGLV S F1 | 1557 | GGTCTGGCGCTGGCTGTGAAGA | + | 422 |
|  | 4 SGLV S F2 | 1657 | YACCGAAGYGGACGMCCGA | + |  |
|  | 4 SGLV S R | 2062 | CTGCTGCTGAGTGGYTGA | - |  |
| **Ji’an nariovirus** |  |  |  |  |  |
|  | Detection |  |  |  |  |
|  | JANV-F | 1964 | CAAGGATGCTGTTCACAA | + | 132 |
|  | JANV-R | 2076 | GGTCTTCAGGAACATACTTTATG | - |  |
|  | JANV-P | 1984 | ATACGGAACTCAGACACGCACAC | + |  |
|  | Genome amplification | | | | |
| Segment L | 1 JANV L F | 1 | GTGGATGCCACAGAGTTGTC | + | 440 |
|  | 1 JANV L R2 | 423 | TTCCAAGTTCCCTCCCCAG | - |  |
|  | 1 JANV L R1 | 484 | CACCATCTTCGCTCACCCA | - |  |
|  | 2 JANV L F | 288 | CACAAGAACAGCYATAGACCA | + | 763 |
|  | 2 JANV L R2 | 1032 | GCTRTCAACAAAAGCCAGT | - |  |
|  | 2 JANV L R1 | 1063 | TTGAACCTGCCAAKACAGT | - |  |
|  | 3 JANV L F1 | 908 | TAGTGCCAGAACAYTCAACTCA | + | 843 |
|  | 3 JANV L F2 | 936 | TGCCTTTCCTTTGAATGACC | + |  |
|  | 3 JANV L R | 1758 | AGATTTGAAATGTAGGCCATC | - |  |
|  | 4 JANV L F1 | 1610 | TAGAGTTCCAGAAGATGCCAC | + | 799 |
|  | 4 JANV L F2 | 1634 | GATCCAGCAATGCATATCAGA | + |  |
|  | 4 JANV L R | 2412 | TGATAGCTTAGTTCTCGCTGT | - |  |
|  | 5 JANV L F | 2333 | CATTTCCATTGCCAGAGGTG | + | 804 |
|  | 5 JANV L R2 | 3115 | CTAAATCATCCAAAAGCCATCC | - |  |
|  | 5 JANV L R1 | 3179 | GAGAYGTGGTTCCTTTAATGGTC | - |  |
|  | 6 JANV L F | 3088 | CTTGAATCCGCTCAGAAGCTA | + | 847 |
|  | 6 JANV L R2 | 3914 | TTTCCCGAAGATTYAACAGGC | - |  |
|  | 6 JANV L R1 | 3963 | AGGTAGATCTCCTTRTGCAAA | - |  |
|  | 7 JANV L F1 | 3801 | AGTTAGAATAGAAGCYACCAG | + | 828 |
|  | 7 JANV L F2 | 3848 | ACAATGACAAAGTAATYCCCTT | + |  |
|  | 7 JANV L R | 4655 | CGTATAACATGCCAAACCTCA | - |  |
|  | 8 JANV L F | 4611 | CCTTCTCCTCAACTCGCAAC | + | 759 |
|  | 8 JANV L R2 | 5350 | TTCGCCTCATACTTGACTCC | - |  |
|  | 8 JANV L R1 | 5436 | ATTGCCTGAACTAGCTCGAAG | - |  |
|  | 9 JANV L F1 | 5166 | CATGTCATCTGCCTCGTCTG | + | 866 |
|  | 9 JANV L F2 | 5199 | GAAAACCCGTTCCTTCAGCTC | + |  |
|  | 9 JANV L R | 6044 | GRTCTGTGCTACCYCTCACCA | - |  |
|  | 10 JANV L F1 | 5872 | TRCTGCTGGCTCAAGTCCC | + | 824 |
|  | 10 JANV L F2 | 6010 | ACACTTACACGTCTAATCAACC | + |  |
|  | 10 JANV L R | 6813 | ATAACTTTGCTACGGATGCTT | - |  |
|  | 11 JANV L F | 6712 | ACAGACCTTGACTTTAAATGGAC | + | 965 |
|  | 11 JANV L R2 | 7657 | AACTGCCTGCATGGGATACA | - |  |
|  | 11 JANV L R1 | 7683 | ACAGTTATCACTTTGGCGTAG | - |  |
|  | 12 JANV L F | 7624 | TTTCACACYCACATGCCTGA | + | 897 |
|  | 12 JANV L R2 | 8501 | CCAACCATYAAAGTCCTCGT | - |  |
|  | 12 JANV L R1 | 8538 | CACAAGGGTGCTCTTCCTC | - |  |
|  | 13 JANV L F | 8378 | GGTTCAAYGACTTTAAAAGGCTA | + | 924 |
|  | 13 JANV L R2 | 9280 | CTTCTTTGAAAAGCCTAAGCTC | - |  |
|  | 13 JANV L R1 | 9327 | TTGAGCCTATTGCTATCTGAG | - |  |
|  | 14 JANV L F | 9198 | CAGCCTCTTCAACCCTGAC | + | 835 |
|  | 14 JANV L R2 | 10011 | GTCTCTAAACTGCTCCAACACA | - |  |
|  | 14 JANV L R1 | 10043 | CTCTTCCGCTAGGTTGCTCA | - |  |
|  | 15 JANV L F | 9836 | TTGAGACAGGCACYACAAGC | + | 871 |
|  | 15 JANV L R2 | 10686 | TCATCAGAAGATAGTGCCATG | - |  |
|  | 15 JANV L R1 | 10737 | GCCCAGTACWACTTTCATRTCA | - |  |
|  | 16 JANV L F1 | 10566 | CTCAAGGCAGAATGACCCAA | + | 1108 |
|  | 16 JANV L F2 | 10651 | CACGAATCCATACTRAGCCTA | + |  |
|  | 16 JANV L R | 11738 | CAGTTTCCRTCAAGRCCCA | - |  |
|  | 16 JANV L F1 | 11631 | TGCRAGCATATCAATGGAGC | + | 378 |
|  | 16 JANV L F2 | 11651 | ATGTAGAGGCCAACACAGTC | + |  |
|  | 16 JANV L R | 12007 | CACACCCAAATTTTTARTCACC | - |  |
| Segment M | 1 JANV M F | 1 | CCCARATCACACTAAGCAGGTTC | + | 400 |
|  | 1 JANV M R2 | 381 | CCTTGTCTAGCTTGGGCATC | - |  |
|  | 1 JANV M R1 | 418 | TTGAAGAGCAGCATCCCCAT | - |  |
|  | 2 JANV M F | 333 | TTCCATTCAGCAGGTRAGCTCA | + | 782 |
|  | 2 JANV M R2 | 1095 | CTTTTATGGCAGTTGTCCCT | - |  |
|  | 2 JANV M R1 | 1195 | CCATTGCAGAAAGTCCGGTA | - |  |
|  | 3 JANV M F | 1039 | TTTGAGTTCCYTGCTCCRA | + | 788 |
|  | 3 JANV M R2 | 1806 | CTTCCGTTCTTACAGTTCTCG | - |  |
|  | 3 JANV M R1 | 1897 | CCTCTTGTAARCAGGCAGA | - |  |
|  | 4 JANV M F | 1711 | ACAATAAGGTTCTTCCGTGTCA | + | 818 |
|  | 4 JANV M R2 | 2507 | TTAGGGCTAGTGATCTCCCACA | - |  |
|  | 4 JANV M R1 | 2542 | ATGGAGACAAAGACACGYTT | - |  |
|  | 5 JANV M F | 2368 | GTCAGTGATGCCTACTCTCCA | + | 743 |
|  | 5 JANV M R2 | 3091 | AGCTGTAAATCCCCAACAGG | - |  |
|  | 5 JANV M R1 | 3200 | RTTRACACCTGCCCATGACA | - |  |
|  | 6 JANV M F | 3022 | GACCTTACAAAGAAGCTGCACA | + | 731 |
|  | 6 JANV M R2 | 3732 | GAACATGCACTAACCTTCCCT | - |  |
|  | 6 JANV M R1 | 3786 | ARGGTTCTCCTATGCTCCAAC | - |  |
|  | 7 JANV M F1 | 3633 | GGCACAAAGTGATTCTACGAC | + | 766 |
|  | 7 JANV M F2 | 3666 | AAGGTTTTATGCTGCAGTTCC | + |  |
|  | 7 JANV M R | 4409 | AGCACACCCTAATATTTACRTCC | - |  |
| Segment S | 1 JANV S F | 1 | GTGCTGCACACCTAAAGTCT | + | 341 |
|  | 1 JANV S R2 | 321 | CACATTCATAGATRGGTGCCAT | - |  |
|  | 1 JANV S R1 | 348 | CCACTCCGTCACAAGAGGC | - |  |
|  | 2 JANV S F | 236 | TGATGCATACGTCACGGCAAA | + | 704 |
|  | 2 JANV S R2 | 921 | CTCCAAAGAAGGCAGCCAT | - |  |
|  | 2 JANV S R1 | 1067 | AAAGAGYTCAGGCTTCACC | - |  |
|  | 3 JANV S F | 802 | GGCTCATCAACAAAAGGTACAGA | + | 772 |
|  | 3 JANV S R2 | 1554 | CGGCCAACCTAGTTTATCCC | - |  |
|  | 3 JANV S R1 | 1630 | GCCTGTTTTCCCACACCCA | - |  |
|  | 4 JANV S F1 | 1509 | ATGGCCCCTTTGTCAACGTCT | + | 277 |
|  | 4 JANV S F2 | 1534 | CCCTTGCCGTCAACATCCA | + |  |
|  | 4 JANV S R | 1790 | CCTAACTATATTTTCACACAA | - |  |
| **Yichun nariovirus** |  |  |  |  |  |
|  | Detection |  |  |  |  |
|  | YCNV-F | 2172 | CCTTCAGAGTGGACAAAG | + | 130 |
|  | YCNV-R | 2295 | GAGGTTCTGTGAGGTTGA | - |  |
|  | YCNV-P | 2189 | ACCGACCAACAACAACACAGAGAA | + |  |
|  | Genome amplification | | | | |
| Segment L | 1 YCNV L F | 1 | TCTGTTCCCCGATTTCCAAAAA | + | 377 |
|  | 1 YCNV L R2 | 359 | GATCACCCAGCATGAGCAA | - |  |
|  | 1 YCNV L R1 | 407 | CTGACCAGATTCCTGATGTA | - |  |
|  | 2 YCNV L F | 269 | CTAACTGAGCAAACAGCCAC | + | 718 |
|  | 2 YCNV L R2 | 967 | TRTCAGCTGCAGAAACCCTC | - |  |
|  | 2 YCNV L R1 | 1092 | TCTTYTCTGCAACTCCAGCTA | - |  |
|  | 3 YCNV L F1 | 840 | TCTTAAGGAATGTCTCYGCTCA | + | 760 |
|  | 3 YCNV L F2 | 853 | CTCYGCTCAAGACAGTCAC | + |  |
|  | 3 YCNV L R | 1593 | YTCTGCCTTCATTTTCTTCC | - |  |
|  | 4 YCNV L F1 | 1464 | AGRCYAGRGCACTTGACTT | + | 746 |
|  | 4 YCNV L F2 | 1500 | AAAGAACTRTGCCYCTGCGAT | + |  |
|  | 4 YCNV L R | 2227 | TGTGTCTKCTGATCCCTTT | - |  |
|  | 5 YCNV L F | 2185 | CAAAGAGGMCACCGACCAA | + | 721 |
|  | 5 YCNV L R2 | 2886 | TTCATGCTGTGGTGCCCCTA | - |  |
|  | 5 YCNV L R1 | 2973 | CATCTGCCTCAGTGGTCATCG | - |  |
|  | 6 YCNV L F | 2761 | CACCCATGTAGGAGTAAGTGT | + | 816 |
|  | 6 YCNV L R2 | 3555 | CCATTCAACTCCATAATCCTGC | - |  |
|  | 6 YCNV L R1 | 3611 | GTCTTTCAGATGTTTCCCGAAT | - |  |
|  | 7 YCNV L F | 3437 | GACAATAGCTTACTAGGTCCA | + | 743 |
|  | 7 YCNV L R2 | 4159 | GACATTTCAGATTCACGGAAG | - |  |
|  | 7 YCNV L R1 | 4237 | TAGGCATCAGTTCTACTCCC | - |  |
|  | 8 YCNV L F | 4083 | GCTATGTCTTAGTWAACCCWGC | + | 773 |
|  | 8 YCNV L R2 | 4825 | GCCTCTAGTGGATATTGCTCT | - |  |
|  | 8 YCNV L R1 | 4957 | CCTCGAAGATTAGTAACTCTCGT | - |  |
|  | 9 YCNV L F1 | 4700 | CCGTCAATAGAGGATCCAAT | + | 902 |
|  | 9 YCNV L F2 | 4741 | AAAATACAGAATCGCACCTGA | + |  |
|  | 9 YCNV L R | 5623 | CTATGAAGCCATAATCTCCC | - |  |
|  | 10 YCNV L F | 5574 | AAAAYAGAACTTGCCCYAAA | + | 772 |
|  | 10 YCNV L R2 | 6325 | AAGTATCGACCTACCTCACAC | - |  |
|  | 10 YCNV L R1 | 6434 | CAAGTCGCCTGACAGTCTC | - |  |
|  | 11 YCNV L F | 6275 | GATAGGAAGGCAAATGCTCCC | + | 743 |
|  | 11 YCNV L R2 | 6995 | GCTATTACACGTTCTAACCGTTC | - |  |
|  | 11 YCNV L R1 | 7034 | GGCACTAATACACAGCCCAT | - |  |
|  | 12 YCNV L F | 6892 | TGATGCAATTAWGGAGTGCTA | + | 900 |
|  | 12 YCNV L R2 | 7771 | TAGATTGTGCAYAGTTCAGAC | - |  |
|  | 12 YCNV L R1 | 7687 | GTATACTGTGTCACGTCCC | - |  |
|  | 13 YCNV L F | 7727 | CTGTACCCAAACTTTGAACAGG | + | 803 |
|  | 13 YCNV L R2 | 8509 | GCCCCAATAAATATRTCAAGG | - |  |
|  | 13 YCNV L R1 | 8553 | GTCTATCAAGGGCTTACTCA | - |  |
|  | 14 YCNV L F | 8368 | ACTTGCTGAGATTGTTAACCAC | + | 829 |
|  | 14 YCNV L R2 | 9176 | CATACGGCCAGTCAATTTGTG | - |  |
|  | 14 YCNV L R1 | 9256 | CTGTCGTACTAGTYCCCATC | - |  |
|  | 15 YCNV L F | 9094 | AGCCCTCTATGCTATGGAA | + | 786 |
|  | 15 YCNV L R2 | 9859 | TTTGCTACTCTGGCTAAYTCT | - |  |
|  | 15 YCNV L R1 | 10037 | TCCTGCTCTATGAAACTTACCAG | - |  |
|  | 16 YCNV L F | 9711 | ACTTTATAATGCTGGTTATGCT | + | 1017 |
|  | 16 YCNV L R2 | 10709 | TACTACCCATGCCAGGTTC | - |  |
|  | 16 YCNV L R1 | 10754 | AATAGTCTCTTCAGTCCGAAC | - |  |
|  | 17 YCNV L F1 | 10511 | GGCAGAATTATATTGACTAGCA | + | 619 |
|  | 17 YCNV L F2 | 10571 | AACATGATTGAAGCTGGCCTA | + |  |
|  | 17 YCNV L R | 11169 | CTCACGGTTGTATGAAGCCT | - |  |
|  | 18 YCNV L F | 11070 | GTGTCAGCCTTACTGCACA | + | 857 |
|  | 18 YCNV L R2 | 11907 | TGAACCACGGCTAAACAACC | - |  |
|  | 18 YCNV L R1 | 11934 | AGCTGGRTTATTTGCAACCT | - |  |
|  | 19 YCNV L F | 11744 | CATTAYATTGCTATTGATGGT | + | 804 |
|  | 19 YCNV L R2 | 12529 | TTGCTCCATATCCAGTGTG | - |  |
|  | 19 YCNV L R1 | 12669 | TGAACTAGCTTGATCCCCTC | - |  |
|  | 20 YCNV L F | 12408 | TGATGCAGAACTTCTTAACCT | + | 836 |
|  | 20 YCNV L R2 | 13224 | TATTACCACCCTTCTGGAGC | - |  |
|  | 20 YCNV L R1 | 13252 | ACTAACATCCCTATTAAGCCTA | - |  |
|  | 21 YCNV L F1 | 13135 | ATCTGGCTCAACACTCCAA | + | 736 |
|  | 21 YCNV L F2 | 13161 | AAATTTTGAAAAGRGCYGGTC | + |  |
|  | 21 YCNV L R | 13877 | TTGAAGATGCAAGCAGCAAT | - |  |
|  | 22 YCNV L F1 | 13703 | TTTAGTCAGCTGGTTAGTTCC | + | 999 |
|  | 22 YCNV L F2 | 13752 | CTCCTCTYTTGTGCCGACT | + |  |
|  | 22 YCNV L R | 14731 | AATTTAATGTCAGCTGTCGG | - |  |
|  | 23 YCNV L F1 | 14396 | ATAGCTCATTCAACAAAACCTG | + | 378 |
|  | 23 YCNV L F2 | 14450 | ATAGACTTACTGCAACTCCTCA | + |  |
|  | 23 YCNV L R | 14809 | GTCCTTCCCCCCTATTTCC | - |  |
| Segment S | 1 YCNV S F | 1 | AAGATATCTAACCCCCGGAGT | + | 334 |
|  | 1 YCNV S R2 | 318 | TTTCCTGCAGAAGCTCAGT | - |  |
|  | 1 YCNV S R1 | 343 | TTTGAAATGAGATCAACCTGC | - |  |
|  | 2 YCNV S F | 194 | CGTGGATCCCAATTTATTTGC | + | 846 |
|  | 2 YCNV S R2 | 1019 | ATCTCCTCGACATACAAGTCT | - |  |
|  | 2 YCNV S R1 | 1045 | CCTGCTCTTGTTGTTTGCT | - |  |
|  | 3 YCNV S F | 972 | AACAACCGAACTTACTCTCC | + | 780 |
|  | 3 YCNV S R2 | 1733 | TGCAGCAGTCTCATCACCA | - |  |
|  | 3 YCNV S R1 | 1783 | CCAGCATTCCATCAAAGCAA | - |  |
|  | 4 YCNV S F | 1657 | GAGACACAGAGATCGACAACC | + | 785 |
|  | 4 YCNV S R2 | 2421 | GTAAGACTCCAGACCAGCAAA | - |  |
|  | 4 YCNV S R1 | 2503 | CCACTTCGTCCAAAAGCAT | - |  |
|  | 5 YCNV S F | 2363 | TCACAACCTGTCTGGCAAT | + | 614 |
|  | 5 YCNV S R2 | 2954 | CTAATCTTAGCAGCAAGCCAA | - |  |
|  | 5 YCNV S R1 | 2988 | TGCGGCATCAACTGACACA | - |  |
|  | 6 YCNV S F1 | 2595 | AAAACAGCCAAGAAACCAA | + | 756 |
|  | 6 YCNV S F2 | 2694 | AAAAGAAGGCTGTGATCCTC | + |  |
|  | 6 YCNV S R | 3429 | CTGCCCAACATAACAACACAA | - |  |
|  | 7 YCNV S F1 | 3313 | CCTGCTCAACTGCTATGGTG | + | 346 |
|  | 7 YCNV S F2 | 3406 | ACTTGTAATCCCCTTTAGTGT | + |  |
|  | 7 YCNV S R | 3732 | CCCCGAATTTACAGAACCTT | - |  |
| **Shanxi tick virus 2** |  |  |  |  |  |
|  | Detection |  |  |  |  |
|  | SXTV-2 -F | 75 | TGGTGGAGACAGTTTAAG | + | 106 |
|  | SXTV-2 -R | 162 | GGAAGTTRGTRTCAGGAGG | - |  |
|  | SXTV-2 -P | 97 | AYAACACCAGGCTCACCATAACTA | + |  |
|  | Genome amplification | | | | |
| Segment L | 1 SXTV-2 L F | 1 | TGGATGATGTTGTAAATAAGA | + | 305 |
|  | 1 SXTV-2 L R2 | 285 | TTCGCTTCATAGAGTTTGTCC | - |  |
|  | 1 SXTV-2 L R1 | 313 | ATACTCATCAGAGCTACCGTA | - |  |
|  | 2 SXTV-2 L F | 96 | CTCCAGCACTGATTTAATAGAGG | + | 760 |
|  | 2 SXTV-2 L R2 | 836 | TGGCAGCAACACATAGGTTC | - |  |
|  | 2 SXTV-2 L R1 | 943 | GACACTGCAACATTGTGCTT | - |  |
|  | 3 SXTV-2 L F | 751 | CAGGAGAAGACATTACCCCT | + | 685 |
|  | 3 SXTV-2 L R2 | 1417 | AACTGATTGCCATCTCGTC | - |  |
|  | 3 SXTV-2 L R1 | 1470 | TTGAAAAGCTCCTGAAGTCTG | - |  |
|  | 4 SXTV-2 L F | 1325 | CTTCTTACTTGCTGCTTAACTG | + | 728 |
|  | 4 SXTV-2 L R2 | 2032 | CAATGTGGGCTTCTGATCCTC | - |  |
|  | 4 SXTV-2 L R1 | 2081 | GAAGGAACTTTTCAAACCGAT | - |  |
|  | 5 SXTV-2 L F | 1935 | AGACACTCATTCRACAGCC | + | 847 |
|  | 5 SXTV-2 L R2 | 2763 | CATCATGTCGGTGACCTCT | - |  |
|  | 5 SXTV-2 L R1 | 2814 | GCACTGCCAGACTTTAAGACA | - |  |
|  | 6 SXTV-2 L F | 2639 | ACTGTTGTTGCCTGCTCAGA | + | 623 |
|  | 6 SXTV-2 L R2 | 3239 | CAACAGGACTCATGCGACT | - |  |
|  | 6 SXTV-2 L R1 | 3500 | TTAACCACTATTTGCCCGGTTG | - |  |
|  | 7 SXTV-2 L F | 3025 | CCTGTAGTAACGGCGGAGAA | + | 762 |
|  | 7 SXTV-2 L R2 | 3823 | GTTGATCACAAAGTTCTGCCT | - |  |
|  | 7 SXTV-2 L R1 | 3925 | AGTTTAAATGGCTTCTTTAGGTC | - |  |
|  | 8 SXTV-2 L F | 3633 | AGCGAATACGACAAACCAG | + | 797 |
|  | 8 SXTV-2 L R2 | 4466 | TTGAAGCCTCCTCAAAGTTGC | - |  |
|  | 8 SXTV-2 L R1 | 4495 | GCTTCTCTTACAGTGCTCGAT | - |  |
|  | 9 SXTV-2 L F | 4304 | CCATGCCTTACATAGTACTCA | + | 715 |
|  | 9 SXTV-2 L R2 | 4999 | TCAAGTACCTGCATCTCCCA | - |  |
|  | 9 SXTV-2 L R1 | 5110 | TCATCATCGCTTCTACCACT | - |  |
|  | 10 SXTV-2 L F | 4877 | TTAGTGGAGAYAGGCARCTCA | + | 857 |
|  | 10 SXTV-2 L R2 | 5712 | GGTATCAACTGTGCAAAGCATT | - |  |
|  | 10 SXTV-2 L R1 | 5750 | CCCCTTCTTGATTTCTTCCT | - |  |
|  | 11 SXTV-2 L F1 | 5558 | TYTCAGAGCCCAAAACRAGG | + | 754 |
|  | 11 SXTV-2 L F2 | 5594 | CCATCAAGGTCCTTAGCACAG | + |  |
|  | 11 SXTV-2 L R | 6325 | AGCGAAAAGATTAGGTTTACCAT | - |  |
|  | 12 SXTV-2 L F | 6163 | GCYATCAAGAAAGAGTACCCAG | + | 841 |
|  | 12 SXTV-2 L R2 | 7041 | TAGTCAGGCCATCATCGTTT | - |  |
|  | 12 SXTV-2 L R1 | 7088 | GAGACTATCATGTGCCGACT | - |  |
|  | 13 SXTV-2 L F | 6954 | ACACAGAGATTTGCTGGTTC | + | 690 |
|  | 13 SXTV-2 L R2 | 7623 | TTTAGTTCTGGCATGTGTTCC | - |  |
|  | 13 SXTV-2 L R1 | 7648 | CTTCCTGCGTGTTTTACAGA | - |  |
|  | 14 SXTV-2 L F1 | 7403 | ATAACCATATGATGCAGTTCC | + | 726 |
|  | 14 SXTV-2 L F2 | 7486 | CAGGTTTATATCTCAAAGGGT | + |  |
|  | 14 SXTV-2 L R | 8191 | GTCAACTAGGCTTTCTAGGTC | - |  |
|  | 15 SXTV-2 L F | 8029 | ATTGTACATGGTTTACCGTCT | + | 831 |
|  | 15 SXTV-2 L R2 | 8897 | TTGCTTTAGCCTATCTGCTT | - |  |
|  | 15 SXTV-2 L R1 | 8947 | TCTTTGATCCCTCCTGCCAA | - |  |
|  | 16 SXTV-2 L F | 8752 | GTCAAACAAATGGCAAGCAA | + | 750 |
|  | 16 SXTV-2 L R2 | 9539 | CTGCTCCAACTTGAGAGTGT | - |  |
|  | 16 SXTV-2 L R1 | 9681 | TCTTTTGTGTCAGCCTGCTA | - |  |
|  | 17 SXTV-2 L F | 9369 | GGAGCAAGTCAAAATCAGGA | + | 875 |
|  | 17 SXTV-2 L R2 | 10281 | TCAGAAGTATGTCCGACTCT | - |  |
|  | 17 SXTV-2 L R1 | 10303 | GAACATGCCCTTCCACCTC | - |  |
|  | 18 SXTV-2 L F1 | 9825 | GATTGAAACGGGAACCACCA | + | 829 |
|  | 18 SXTV-2 L F2 | 9854 | ATTGGAGAATACTACACTGCCTA | + |  |
|  | 18 SXTV-2 L R | 10663 | GCCACGTCTAACTTTCTCCT | - |  |
|  | 19 SXTV-2 L F | 10457 | GGACCAACCCAAGAATGCT | + | 797 |
|  | 19 SXTV-2 L R2 | 11234 | TAATGTGTCCCAACTCTCCG | - |  |
|  | 19 SXTV-2 L R1 | 11284 | TCAAATAGGCAAGCCACGAC | - |  |
|  | 20 SXTV-2 L F1 | 11006 | ACTCAGCAATCAAAGTAGAGG | + | 926 |
|  | 20 SXTV-2 L F2 | 11038 | GGTTTACAGACAGCATACCAA | + |  |
|  | 20 SXTV-2 L R | 12001 | CAACGATACATGTGCAGACC | - |  |
| Segment M | 1 SXTV-2 M F | 1 | TTGTGTCCATTCTTTGAACCC | + | 473 |
|  | 1 SXTV-2 M R2 | 457 | CATGCCGTCGGTTAGACTCA | - |  |
|  | 1 SXTV-2 M R1 | 483 | CACCTAGGCCATCAGCAAT | - |  |
|  | 2 SXTV-2 M F | 308 | CATAATGCCTGTGAAAGACCA | + | 759 |
|  | 2 SXTV-2 M R2 | 1047 | ATAACCTTTGTACCTGTGCT | - |  |
|  | 2 SXTV-2 M R1 | 1068 | TGTTTCAAATTTTAGGGCAGT | - |  |
|  | 3 SXTV-2 M F | 899 | GATCACYGTYGAAATCCTCC | + | 796 |
|  | 3 SXTV-2 M R2 | 1729 | CCTACCGTTTCTACAATTTTCGT | - |  |
|  | 3 SXTV-2 M R1 | 1794 | CACAATCTTTCGCGTGCTG | - |  |
|  | 4 SXTV-2 M F | 1598 | CTCYGCYATCTACGTGTTA | + | 796 |
|  | 4 SXTV-2 M R2 | 2373 | CATGAAACCAGTGCCTGTCTC | - |  |
|  | 4 SXTV-2 M R1 | 2437 | AGTGTGCTCAAGTATAGACACAA | - |  |
|  | 5 SXTV-2 M F1 | 2190 | CTCCATGTTGGTGAYGCATA | + | 844 |
|  | 5 SXTV-2 M F2 | 2211 | GCACCTTCYTACAGTGGYAAACA | + |  |
|  | 5 SXTV-2 M R | 3092 | AGTAAGATGTAACACCTGCC | - |  |
|  | 6 SXTV-2 M F | 2885 | GAGCCTTTGTGACATCCAGT | + | 799 |
|  | 6 SXTV-2 M R2 | 3720 | TAGGCAGCTAGCATAACCTGT | - |  |
|  | 6 SXTV-2 M R1 | 3806 | CAATGAGAAGCCCTACCCAG | - |  |
|  | 7 SXTV-2 M F1 | 3514 | TTATAGCTCGCAGAGACACCAC | + | 673 |
|  | 7 SXTV-2 M F2 | 3609 | CAAGACAGTTCAGTGGCATC | + |  |
|  | 7 SXTV-2 M R | 4263 | TGACCGGCCTCGTTAAGCA | - |  |
| Segment S | 1 SXTV-2 S F | 1 | TTAATCAAACTCCTGCCACC | + | 498 |
|  | 1 SXTV-2 S R2 | 478 | ACTGTAGACCTTAACAACCTGCT | - |  |
|  | 1 SXTV-2 S R1 | 565 | CATCTTCAATGCCCAGTGCTT | - |  |
|  | 2 SXTV-2 S F | 351 | ATGCTTAAAGGACGGCTCCCAA | + | 828 |
|  | 2 SXTV-2 S R2 | 1169 | CTTGCCATTCTCCCAGCTGTG | - |  |
|  | 2 SXTV-2 S R1 | 1190 | CCCAAAGCTGCATACCATCTCG | - |  |
|  | 3 SXTV-2 S F1 | 937 | GGTCTGCAATTGACACAGCCTT | + | 540 |
|  | 3 SXTV-2 S F2 | 1000 | AGACCTTCCCYTCTCTGTCA | + |  |
|  | 3 SXTV-2 S R | 1520 | CTGACATGCCTGATGCACGG | - |  |
| **Wetland virus** |  |  |  |  |  |
|  | Detection |  |  |  |  |
|  | WELV-F | 262 | CGAGGCCACTAGATTCTG | + | 110 |
|  | WELV-R | 365 | CCAGACCTTGACAATGTC | - |  |
|  | WELV-P | 294 | AGTGTGCTTGGTCCTCCTCTAC | + |  |
|  | Genome amplification | | | | |
| Segment L | 1 WELV L F | 1 | CCCTTATCCCCAGAGTTAACAT | + | 441 |
|  | 1 WELV L R1 | 437 | AGTGTTCAACGGTCCCCTCCCG | - |  |
|  | 1 WELV L R2 | 421 | TCCCGAATCTTGTTGCACCCA | - |  |
|  | 2 WELV L F | 373 | TGGGAGTCACTGTTGTGCTT | + | 763 |
|  | 2 WELV L R1 | 1249 | GTCTTTGCACCAACTTTCCGT | - |  |
|  | 2 WELV L R2 | 1114 | AGGCAGTTTGTAATGAACCTCT | - |  |
|  | 3 WELV L F1 | 918 | AGGCACAAGCTGTATCTCAC | + | 905 |
|  | 3 WELV L F2 | 1046 | CATCTGCACCGTTAGCCTT | + |  |
|  | 3 WELV L R | 1931 | GGAGAACATCCCCTCATAAGCG | - |  |
|  | 4 WELV L F | 1872 | ATACGAAACTCCACTAGCCAT | + | 989 |
|  | 4 WELV L R1 | 2942 | AACACCGTCAACTAGACCC | - |  |
|  | 4 WELV L R2 | 2840 | GTTTCTTGCACATCCTTGGTC | - |  |
|  | 5 WELV L F | 2763 | ATAGTTGTAGGGTCGATAAGCAC | + | 1047 |
|  | 5 WELV L R1 | 3834 | GTCTCTGATGTCTTCCTCGTC | - |  |
|  | 5 WELV L R2 | 3788 | CCTGCTGAAATTAACTTTGCTC | - |  |
|  | 6 WELV L F | 3723 | TTGCGTGATCTCAAAGTTCCT | + | 958 |
|  | 6 WELV L R1 | 4742 | AGTTCACAGTCTCTATTTGCG | - |  |
|  | 6 WELV L R2 | 4658 | CTGAGAATTTCTTGCCTAACTCC | - |  |
|  | 7 WELV L F | 4588 | AGCAAAACAAGCAGTTACAGA | + | 972 |
|  | 7 WELV L R1 | 5595 | GCACCTAGCTTCTTTAGTTTCCC | - |  |
|  | 7 WELV L R2 | 5540 | CCAGCAAGACCTTGTAGCAG | - |  |
|  | 8 WELV L F | 5259 | GATTATCAACAGGCCGTGAC | + | 1068 |
|  | 8 WELV L R1 | 6337 | AACACTTCCGTTGAATTCCA | - |  |
|  | 8 WELV L R2 | 6306 | ATGTCTAACAAGGTAGGCTTC | - |  |
|  | 9 WELV L F | 6246 | ACGAACTTCATATTCTACTGTGC | + | 1039 |
|  | 9 WELV L R1 | 7309 | TTAATCTCTCAACCCCGCCTT | - |  |
|  | 9 WELV L R2 | 7264 | CGTTCAGAATTTTCCGAATGC | - |  |
|  | 10 WELV L F | 7238 | CCGTCAGGTCGAAATACCAG | + | 1117 |
|  | 10 WELV L R1 | 8378 | ATGTGGCATAAAGCTCAGTC | - |  |
|  | 10 WELV L R2 | 8336 | TGCTGTCTGATCCCCGTTC | - |  |
|  | 11 WELV L F | 8319 | CTCAGGATTCTTAGAGAGAACGG | + | 1062 |
|  | 11 WELV L R1 | 9406 | AGTAATCCACCTTAGTCGGTT | - |  |
|  | 11 WELV L R2 | 9361 | TAGCAGGTAGCCCATGACCA | - |  |
|  | 12 WELV L F | 9335 | TGTGCGACTCTCCAATAACCC | + | 1143 |
|  | 12 WELV L R1 | 10494 | ATACACGATGGAGAGACACCC | - |  |
|  | 12 WELV L R2 | 10457 | CTAGGCTCGCTTTCAATGTCA | - |  |
|  | 13 WELV L F | 10386 | GTTGCAAAGAGACAGACACCG | + | 1081 |
|  | 13 WELV L R1 | 11481 | CAAAGAAAACAGCTCCTAGTGA | - |  |
|  | 13 WELV L R2 | 11446 | TTTTGGCAATCTTGACCCGTT | - |  |
|  | 14 WELV L F1 | 11339 | TTTGAACAGCCGTGTGTCCC | + | 721 |
|  | 14 WELV L F2 | 11361 | GAACTTAGGGCTGATGGGCCAA | + |  |
|  | 14 WELV L R | 12039 | CAAGTCAGTAGATGTTCTGCAC | - |  |
| Segment M | 1 WELV M F | 1 | AATACTTGAGCTTCAGGC | + | 426 |
|  | 1 WELV M R1 | 478 | TGGCTGAAAAGCACGACACC | - |  |
|  | 1 WELV M R2 | 408 | ACATCTGACGGGCTCATCC | - |  |
|  | 2 WELV M F | 374 | CTGCCCACTCTCAACCCGAA | + | 936 |
|  | 2 WELV M R1 | 1335 | CACCTGGGCAATCAATTTGGA | - |  |
|  | 2 WELV M R2 | 1289 | AGCTAGGCAATCCTTGTCACA | - |  |
|  | 3 WELV M F | 1151 | CACCCAAAAGAAGCCTCAACC | + | 1033 |
|  | 3 WELV M R1 | 2202 | TGCTGCATTGAGCCTCACGA | - |  |
|  | 3 WELV M R2 | 2162 | TGGCTTTCTTGCTGTACTGTCT | - |  |
|  | 4 WELV M F | 2101 | GGCCGCTGTTCTACTGACC | + | 863 |
|  | 4 WELV M R1 | 2965 | ACGCATCTGTTCTTACGTACTCA | - |  |
|  | 4 WELV M R2 | 2945 | TGCTCCACTTTACGGCGAA | - |  |
|  | 5 WELV M F | 2820 | CCACTTGCCTCTTTAAGTCCT | + | 1002 |
|  | 5 WELV M R1 | 3887 | AGCACATTTCTTGGTGTCTGT | - |  |
|  | 5 WELV M R2 | 3801 | CAGTGAAAGCCCTTACCTTGC | - |  |
|  | 6 WELV M F1 | 3701 | AAGCCTGATGAGTTCACTGTC | + | 966 |
|  | 6 WELV M F2 | 3727 | CAGAAGCACAGATCCTAACACC | + |  |
|  | 6 WELV M R | 4670 | CACACTCCAAACTTTAACACAAA | - |  |
| Segment S | 1 WELV S F | 1 | GTACGCCCCACGTTTTCACAC | + | 534 |
|  | 1 WELV S R1 | 577 | TTCCTTCGCCTGATCATGTCG | - |  |
|  | 1 WELV S R2 | 513 | TTGTACTCTGCTGCAATGGCTT | - |  |
|  | 2 WELV S F | 437 | TCCTACCAAAAGGCTGCGAT | + | 789 |
|  | 2 WELV S R1 | 1226 | ACCGAAAGTGATTCCAAGCTC | - |  |
|  | 2 WELV S R2 | 1206 | GGACATTCTCCCCGAGGTCA | - |  |
|  | 3 WELV S F1 | 986 | GTCTTCAGCAGCTTCTACTGG | + | 624 |
|  | 3 WELV S F2 | 1030 | AGTCACTTTCCCGTCGGTT | + |  |
|  | 3 WELV S R | 1631 | ATAAAGCAGAAAAGAATATCAAA | - |  |
| **Yanbian Nairo tick virus 1** | |  |  |  |  |
|  | Detection |  |  |  |  |
|  | YBNTV-1-F | 2349 | GARAGCTTTGACAAGGTC | + | 534 |
|  | YBNTV-1-R | 2457 | GTCCRACAGGTGATGATG | - |  |
|  | YBNTV-1-P | 2368 | TACCAAGRATAGCCAGGATAGCCAA | + |  |
|  | Genome amplification | | | | |
| Segment L | 1 YBNTV-1 L F | 1 | TCCTGCACACCCAAGAACCTA | + | 778 |
|  | 1 YBNTV-1 L R2 | 762 | CTYCCCACTCKCATTGGRA | - |  |
|  | 1 YBNTV-1 L R1 | 805 | TTTGCTGGACYTTAACGTTG | - |  |
|  | 2 YBNTV-1 L F | 695 | CTARCCAAGCCYRCTCCCA | + | 896 |
|  | 2 YBNTV-1 L R2 | 1571 | TAGTACATRTTGGAMACCCC | - |  |
|  | 2 YBNTV-1 L R1 | 1652 | TAYGCCTTGCCRTATTTGGA | - |  |
|  | 3 YBNTV-1 L F | 1494 | ARCAGAATGGCTTYACAAA | + | 918 |
|  | 3 YBNTV-1 L R2 | 2393 | CRGCCTTRCCYTCAGTGAC | - |  |
|  | 3 YBNTV-1 L R1 | 2422 | TCACTACWGCTKCYTCCG | - |  |
|  | 4 YBNTV-1 L F | 2283 | CYGAGGTGCCAATWCAYGA | + | 971 |
|  | 4 YBNTV-1 L R2 | 3234 | CTTTGAGTGGCARCARGCTA | - |  |
|  | 4 YBNTV-1 L R1 | 3255 | CCACTAGCCCCTTCTTHACTG | - |  |
|  | 5 YBNTV-1 L F1 | 3152 | ATYAAGAAAACTGCAAAAGCYTC | + | 834 |
|  | 5 YBNTV-1 L F2 | 3178 | TTCYCTAGCTGAGCAAATCC | + |  |
|  | 5 YBNTV-1 L R | 4114 | ACYCTCCTWACTTTRATTCCA | - |  |
|  | 6 YBNTV-1 L F | 3975 | CAACGCCRTACTTYGACTG | + | 1009 |
|  | 6 YBNTV-1 L R2 | 4963 | TGTAGTTCCCAMGAYACRTGC | - |  |
|  | 6 YBNTV-1 L R1 | 5182 | AAACTTCTTRCCTGAGCTA | - |  |
|  | 7 YBNTV-1 L F1 | 4802 | CTTTGCCCTAARGTYACACT | + | 1129 |
|  | 7 YBNTV-1 L F2 | 4860 | AYAGGCAGTTGATATTTGACA | + |  |
|  | 7 YBNTV-1 L R | 5968 | CTGYGWTAGGCTCTCYAACTC | - |  |
|  | 8 YBNTV-1 L F1 | 5758 | AAACTGGCATTYTTYCCCTG | + | 992 |
|  | 8 YBNTV-1 L F2 | 5862 | CAGCAGGYATYAAGAAGCAC | + |  |
|  | 8 YBNTV-1 L R | 6835 | CTGCTGTAATATYGCCATG | - |  |
|  | 9 YBNTV-1 L F | 6700 | GACAYTGAAYCTRATAGCCAA | + | 974 |
|  | 9 YBNTV-1 L R2 | 7652 | ATACTGTTATGACCTTRGCRTA | - |  |
|  | 9 YBNTV-1 L R1 | 7881 | ACTGTTAATRAGYCCWGTGA | - |  |
|  | 10 YBNTV-1 L F | 7515 | ARGGYATACACCATGCCAC | + | 1005 |
|  | 10 YBNTV-1 L R2 | 8500 | TAGGKGACTTRCTTARCCTT | - |  |
|  | 10 YBNTV-1 L R1 | 8566 | CATACCTRAGGAGCCTYAG | - |  |
|  | 11 YBNTV-1 L F | 8380 | GCTYGAATACCTYAGGACCA | + | 1043 |
|  | 11 YBNTV-1 L R2 | 9402 | TTCGGATYTTTACTTGTTCYC | - |  |
|  | 11 YBNTV-1 L R1 | 9441 | TTRCTYAGRTGGAAGCCAA | - |  |
|  | 12 YBNTV-1 L F | 9120 | TCTCTTCACTGYTRGTTCG | + | 1041 |
|  | 12 YBNTV-1 L R2 | 10141 | TGCCTCTGACATTGAAGTCG | - |  |
|  | 12 YBNTV-1 L R1 | 10333 | TAGTCATGCTGCTCYTTTGT | - |  |
|  | 13 YBNTV-1 L F | 10081 | GACTAYATAAAAGAYCCYA | + | 833 |
|  | 13 YBNTV-1 L R2 | 10895 | TCTCYGTYYTGTTTGACAC | - |  |
|  | 13 YBNTV-1 L R1 | 10939 | TGCCACRACCATYAAGTGC | - |  |
|  | 14 YBNTV-1 L F1 | 10730 | ATACAGTTCAAGCAAGAGGCAA | + | 1052 |
|  | 14 YBNTV-1 L F2 | 10773 | TGCACTTCTACCTTGTCCAC | + |  |
|  | 14 YBNTV-1 L R | 11762 | GCCTCYACDCCTATTAGGAA | - |  |
|  | 15 YBNTV-1 L F1 | 11424 | GCACCATTGAGTTTGAACC | + | 474 |
|  | 15 YBNTV-1 L F2 | 11478 | TAAAGGTRCCYCARATYAGAGC | + |  |
|  | 15 YBNTV-1 L R | 11933 | TTTACCTCCGGCATCTGTC | - |  |
| Segment M | 1 YBNTV-1 M F | 1 | GCTTTCCCACAAGTAACCC | + | 848 |
|  | 1 YBNTV-1 M R2 | 830 | GGCCRTGRTTYTCYACAGT | - |  |
|  | 1 YBNTV-1 M R1 | 920 | TTCACTGTWATATGVACBAC | - |  |
|  | 2 YBNTV-1 M F | 773 | GTTGTGATWGCYTAYGCYACC | + | 988 |
|  | 2 YBNTV-1 M R2 | 1740 | TGCCYCTTGTTYTRTCTGTYC | - |  |
|  | 2 YBNTV-1 M R1 | 1796 | TCATGCCGCTGCCARTGGAA | - |  |
|  | 3 YBNTV-1 M F1 | 1639 | TGCYTGGTACTTCCTYGGRT | + | 1123 |
|  | 3 YBNTV-1 M F2 | 1676 | CTYCTGTTYRYAATGTRTCTGG | + |  |
|  | 3 YBNTV-1 M R | 2779 | TASTCAAGYTCCCACTTYGT | - |  |
|  | 4 YBNTV-1 M F | 2641 | CACAGAYCAYCTYTGCCA | + | 951 |
|  | 4 YBNTV-1 M R2 | 3572 | CTYACAGARGGGACAACCAC | - |  |
|  | 4 YBNTV-1 M R1 | 3609 | CTCCCTTGCTTCTGTDGAGT | - |  |
|  | 5 YBNTV-1 M F1 | 3287 | CCTGCAGTYAAYTACTTYTTCCA | + | 830 |
|  | 5 YBNTV-1 M F2 | 3339 | CAACAYTGCTRCTYAAGGGRA | + |  |
|  | 5 YBNTV-1 M R | 4150 | GTAGACTCCTGTCCCACCA | - |  |
|  | 6 YBNTV-1 M F1 | 3789 | CCAACTGCACCACAGGCTA | + | 538 |
|  | 6 YBNTV-1 M F2 | 3865 | CTCHARGTTCTTTGGCTCCTG | + |  |
|  | 6 YBNTV-1 M R | 4383 | TCAARRATAAYGTGGCAGCA | - |  |
| Segment S | 1 YBNTV-1 S F | 1 | TTGCTGCACCACAAAACCTA | + | 804 |
|  | 1 YBNTV-1 S R2 | 784 | CAGTCYCCTCCYTTYAGCC | - |  |
|  | 1 YBNTV-1 S R1 | 818 | TTCTTRTCCCACTCTCCCCAT | - |  |
|  | 2 YBNTV-1 S F | 609 | CAGTACACCAAGGCCAACCAG | + | 737 |
|  | 2 YBNTV-1 S R2 | 1326 | CTTGCCATYCTTCCTGCAGT | - |  |
|  | 2 YBNTV-1 S R1 | 14673 | AACTCACGGAACAGTGCAGA | - |  |
|  | 3 YBNTV-1 S F1 | 1171 | TKTCRGCCATGCTGTTCTC | + | 556 |
|  | 3 YBNTV-1 S F2 | 1199 | ATCGCCRCAAGGTAAAACCAA | + |  |
|  | 3 YBNTV-1 S R | 1736 | TCTCCCGTCCGCCCTGCTG | - |  |
| **Yezo virus** |  |  |  |  |  |
|  | Detection |  |  |  |  |
|  | YEZV-F | 4206 | GTCTGGGATCAAAGTCAG | + | 89 |
|  | YEZV-R | 4275 | AGCACTTCATATTCTCCTTC | - |  |
|  | YEZV-P | 4225 | AGAGTAAGGCACACCAGCATCAAC | + |  |
|  | Genome amplification | | | | |
| Segment L | 1 YEZV L F | 1 | CCCACTAAGGCTAACCTCCA | + | 900 |
|  | 1 YEZV L R2 | 875 | TCTACGAACAGGACCRACAG | - |  |
|  | 1 YEZV L R1 | 916 | GTTTGTTCRTTTACCAGCAT | - |  |
|  | 2 YEZV L F | 706 | CCAGCTGTATCTAGAGAGGAC | + | 845 |
|  | 2 YEZV L R2 | 1530 | CGTGTTAARCTGTCTAACTCC | - |  |
|  | 2 YEZV L R1 | 1725 | GCTCYTGTCTTGAGATTCCA | - |  |
|  | 3 YEZV L F1 | 1380 | CTTAAACAAACTAACCCCACA | + | 954 |
|  | 3 YEZV L F2 | 1477 | GTTGATCTTTTCTCCTGCCTT | + |  |
|  | 3 YEZV L R | 2410 | TTTYACACCCGACTCTTCCAA | - |  |
|  | 4 YEZV L F | 2197 | TGCACTTACATACCTGAAGACC | + | 867 |
|  | 4 YEZV L R2 | 3044 | TCTGTCCATTGTAACCCCTT | - |  |
|  | 4 YEZV L R1 | 3194 | CTCAATGTCCTCCATMAGCCAA | - |  |
|  | 5 YEZV L F1 | 2861 | TTGCCAACATCAGCACAAC | + | 933 |
|  | 5 YEZV L F2 | 2971 | AARATTAGAGTAAGGCCYAC | + |  |
|  | 5 YEZV L R | 3882 | TAGCGAAAAGAGGTTAAACTTC | - |  |
|  | 6 YEZV L F1 | 3768 | GCGACTCACWGGATATTCAGA | + | 974 |
|  | 6 YEZV L F2 | 3734 | CCTGCAAGATAATGAACACTC | + |  |
|  | 6 YEZV L R | 4661 | TAAGTAACAAGGTGTCYCCAA | - |  |
|  | 7 YEZV L F1 | 4434 | CTRACTAAGGARCAYGCATC | + |  |
|  | 7 YEZV L F2 | 4577 | GACAGGCAACTTTCTTGACC | + |  |
|  | 7 YEZV L R | 5265 | TTTAGCCTRCCAAACAGAGT | - |  |
|  | 8 YEZV L F1 | 5044 | CAGCAGAAAAGCACATGTCA | + | 773 |
|  | 8 YEZV L F2 | 5153 | TAGGCATGAACTTCCAACTGA | + |  |
|  | 8 YEZV L R | 5905 | GCCATACTTTTCAACCAAC | - |  |
|  | 9 YEZV L F | 5763 | GACACACTAACAGAGGCACA | + | 853 |
|  | 9 YEZV L R1 | 6664 | CTCATCAGGCTTCATTGCAT | - |  |
|  | 9 YEZV L R2 | 6594 | AGTAYCTGTATAGCCAATCCAG | - |  |
|  | 10 YEZV L F1 | 6402 | TGTATGCACTATAAGTCCTTCG | + | 819 |
|  | 10 YEZV L F2 | 6440 | GATACTTGAAACCGAGCCTT | + |  |
|  | 10 YEZV L R | 7238 | CCCCACTTTGTATTRTCTCCT | - |  |
|  | 11 YEZV L F | 7026 | GGCACTAAAGTAGTCCATGCAA | + | 874 |
|  | 11 YEZV L R2 | 7879 | AAAACTCCAGCATTGTGTCTC | - |  |
|  | 11 YEZV L R1 | 7930 | AATAAACTTTATAACAGCTGGGG | - |  |
|  | 12 YEZV L F | 7721 | TGACTACGCAAAGTGCATC | + | 887 |
|  | 12 YEZV L R2 | 8590 | ACTCAAAGTTAGAYCGGAT | - |  |
|  | 12 YEZV L R1 | 8642 | ATCTAACAAGGCCAAATGTYC | - |  |
|  | 13 YEZV L F | 8399 | TTCATCCAACTTCCGCTTT | + | 958 |
|  | 13 YEZV L R2 | 9369 | GTTTCAACCTATTCGGCTCT | - |  |
|  | 13 YEZV L R1 | 9337 | CGCTAAACTCCTAAAYTCCGA | - |  |
|  | 14 YEZV L F1 | 9184 | GGATACTATCAAGCACYCTGT | + | 929 |
|  | 14 YEZV L F2 | 9216 | AAATCACAGGACTTTGAGCCA | + |  |
|  | 14 YEZV L R | 10126 | TCTTCTAGTTGATGCCGTGT | - |  |
|  | 15 YEZV L F | 9991 | GGAAGCCTAATAAATCCATTGC | + | 810 |
|  | 15 YEZV L R1 | 10865 | GCTCCATCCTAACTTCCTCAG | - |  |
|  | 15 YEZV L R2 | 10781 | TGATCTTTCCTATCACGACCT | - |  |
|  | 16 YEZV L F | 10671 | GARTCAGCCAMTTACCACGAG | + | 938 |
|  | 16 YEZV L R1 | 11625 | CATCAAAGAAAAGATTGTAGGCA | - |  |
|  | 16 YEZV L R2 | 115901 | TGAGCCATTGCCTTTTYYC | - |  |
|  | 17 YEZV L F1 | 11338 | AAYTGTCAGCATCYAGCCTA | + | 725 |
|  | 17 YEZV L F2 | 11384 | CAGTTGCACTTTGTATCAGCTA | + |  |
|  | 17 YEZV L R | 12074 | TACCCCCCTATTATAACCTTT | - |  |
| Segment M | 1 YEZV M F | 1 | TCATTCAAAAGTCCCAACTTG | + | 870 |
|  | 1 YEZV M R2 | 852 | GCCTCTTGCATCAAATCC | - |  |
|  | 1 YEZV M R1 | 910 | CTCAACRCTGTTTTCTGTCA | - |  |
|  | 2 YEZV M F1 | 711 | CCMGGATGTGAGAAATTACACAA | + | 859 |
|  | 2 YEZV M F2 | 779 | ATCAGAGAATATAAAGGTCAGC | + |  |
|  | 2 YEZV M R | 1592 | TAGATCAACCTTATTGGTGTGC | - |  |
|  | 3 YEZV M F | 1503 | ATCATAAAATGCACCAAGGAG | + | 926 |
|  | 3 YEZV M R2 | 2408 | TTTTGACTTTGCCAACTGAGG | - |  |
|  | 3 YEZV M R1 | 2449 | TCTGATTTGCCAGCCACCA | - |  |
|  | 4 YEZV M F1 | 2257 | TGCACAACCACTCCATCCA | + | 932 |
|  | 4 YEZV M F2 | 2294 | CCTACTTTCCCTRCCCGTA | + |  |
|  | 4 YEZV M R | 3204 | AGTAGTATTGAAGCACAGTTCC | - |  |
|  | 5 YEZV M F | 3104 | AATTTACAACCTCAATGGCTT | + | 827 |
|  | 5 YEZV M R2 | 3911 | GGCCYTTCCAAATTGATCCA | - |  |
|  | 5 YEZV M R1 | 4070 | GTYTRTATCTYTCCAGYTCT | - |  |
|  | 6 YEZV M F1 | 3714 | GAAGCWCCRAARGARTCAGA | + | 261 |
|  | 6 YEZV M F2 | 3981 | GGMCCTAACCTTTTRAGYCT | + |  |
|  | 6 YEZV M R | 4219 | AARAKTATWATGCACTAAATTCT | - |  |
| Segment S | 1 YEZV S F | 1 | CGTGCTGCGACCCCCCAATAG | + | 625 |
|  | 1 YEZV S R2 | 605 | GTGCTTCTTGTTCCTGCTC | - |  |
|  | 1 YEZV S R1 | 707 | TTTTGTTCCAWCCRCCCCA | - |  |
|  | 2 YEZV S F1 | 419 | GCACAGTAYCARATAGCAG | + | 659 |
|  | 2 YEZV S F2 | 523 | CCCKAAGAARTACGTGAAYGA | + |  |
|  | 2 YEZV S R | 1163 | CGTTRAARTCTCCYGTTGC | - |  |
|  | 3 YEZV S F1 | 944 | GACTCAGTGACCTCAGCRT | + | 698 |
|  | 3 YEZV S F2 | 984 | TTGACACTGCATTTTCCACCT | + |  |
|  | 3 YEZV S R | 1645 | TGTTGKTWSTGCATACCCC | - |  |
| **Hunchun nairovirus** | Detection |  |  |  |  |
|  | HCNV-F | 2832 | GCACTATAGGGCATTCCA | + | 148 |
|  | HCNV-R | 2980 | GTGAGTTGAGCCTTGTATG | - |  |
|  | HCNV-P | 2864 | TAACCAGCACAGAGGAAGCACT | + |  |
| Segment L | 1 HCNV L F | 1 | GAACCATCCTTCACTGTGCCAC | + | 812 |
|  | 1 HCNV L R2 | 793 | TTCATGCCCAACATCTGAG | - |  |
|  | 1 HCNV L R1 | 949 | TGCTGCAACTTTTCTTCCC | - |  |
|  | 2 HCNV L F1 | 576 | ACAAAGTTACCGAAAATGACA | + | 878 |
|  | 2 HCNV L F2 | 687 | AAGTACTAGAGCCTGCACAA | + |  |
|  | 2 HCNV L R | 1545 | TTCAGGCACAGAATCTCCAC | - |  |
|  | 3 HCNV L F1 | 1420 | GTCTGGCATAACAAAAGCAAA | + | 996 |
|  | 3 HCNV L F2 | 2396 | CATTCTGCTGTGCTTATCGG | + |  |
|  | 3 HCNV L R | 2476 | TCCCCACTTGGTTTTCTGT | - |  |
|  | 4 HCNV L F | 2308 | CAATGCCCCAGTACTCTCA | + | 994 |
|  | 4 HCNV L R2 | 3190 | CCTTCTCCATAAATCGGACT | - |  |
|  | 4 HCNV L R1 | 3284 | AATAGATATGTCAGCAGCATC | - |  |
|  | 5 HCNV L F | 3121 | TCCTTCCCCTTTCTTTGTCC | + | 909 |
|  | 5 HCNV L R2 | 4009 | GCATTCATGGCAGATATCCTT | - |  |
|  | 5 HCNV L R1 | 4125 | CTTGCAATCCTACTGGCAT | - |  |
|  | 6 HCNV L F1 | 3800 | GCTGAATTAAGACTTTGCTAC | + | 918 |
|  | 6 HCNV L F2 | 4697 | ATATGTAAGCTCCATAGGGTC | + |  |
|  | 6 HCNV L R | 4722 | TGGAGCAATCTTGTATTTTGACT | - |  |
|  | 7 HCNV L F | 4615 | AGAGGCTATAGATTCAGCA | + | 800 |
|  | 7 HCNV L R2 | 5394 | CTTCCTCACTAATTGCACACC | - |  |
|  | 7 HCNV L R1 | 5608 | TCTATGAAGCCATGTTCTCCT | - |  |
|  | 8 HCNV L F | 5242 | TAGAAAAGCAGGTTCAACTGTC | + | 854 |
|  | 8 HCNV L R2 | 6074 | AGTAATTCTTTAGTTCCCCTGA | - |  |
|  | 8 HCNV L R1 | 6148 | AATGCACAACTACTATTGGTC | - |  |
|  | 9 HCNV L F | 5998 | CATTAAGCCTTTATCTGCAAA | + | 1011 |
|  | 9 HCNV L R2 | 6988 | AAAGATGATATAACACGCTCT | - |  |
|  | 9 HCNV L R1 | 7032 | TTAGTAACATAGCAGGAACCA | - |  |
|  | 10 HCNV L F | 6804 | CATTGAGTAGCCCGAACAC | + | 867 |
|  | 10 HCNV L R2 | 7650 | AGCTCCTAACACTAGAGACCA | - |  |
|  | 10 HCNV L R1 | 7672 | GTATACTGTATCCCGACCC | - |  |
|  | 11 HCNV L F | 7538 | GCAGTTAAACTAGATCAGCTTC | + | 822 |
|  | 11 HCNV L R2 | 8340 | AGCTAATGGCTCTATACACC | - |  |
|  | 11 HCNV L R1 | 8585 | CTGATTAACTCTTTGTATGAGCC | - |  |
|  | 12 HCNV L F | 8203 | TGTAGAGGCACGATCTCTGA | + | 990 |
|  | 12 HCNV L R2 | 9173 | GTTCTCCCTTCATGCGACC | - |  |
|  | 12 HCNV L R1 | 9242 | CAGTTGTACTAGTCCCCAT | - |  |
|  | 13 HCNV L F1 | 8979 | TCTCTTCAAACAGTAACCCAT | + | 979 |
|  | 13 HCNV L F2 | 9936 | TGTCCCATATGATTATAACACT | + |  |
|  | 13 HCNV L R | 10022 | GTTCCACAAAGCTCACCAG | - |  |
|  | 14 HCNV L F | 9811 | AAAAGGAAGAACTCACCCAG | + | 1000 |
|  | 14 HCNV L R2 | 10792 | CTCAATGTCTTTCGGCAAC | - |  |
|  | 14 HCNV L R1 | 10865 | TATTCTATCACAGTCGCCCAT | - |  |
|  | 15 HCNV L F | 10609 | CCTTAGTAGGCTCTCTGAGT | + | 897 |
|  | 15 HCNV L R2 | 11483 | AGGCATATATCAGTCCTATCCAC | - |  |
|  | 15 HCNV L R1 | 11624 | CCATTACTTCAGAGTGCAT | - |  |
|  | 16 HCNV L F | 11401 | TAGCATTATCTGGGGACTAGC | + | 873 |
|  | 16 HCNV L R2 | 12254 | TATTTCCCCTCCTGTATGCC | - |  |
|  | 16 HCNV L R1 | 12481 | TGTTACCAAATAGTGCAACC | - |  |
|  | 17 HCNV L F | 12136 | GATGCTAACAACACTCTGTCG | + | 883 |
|  | 17 HCNV L R2 | 12996 | GCACAAGATGTTACTTTTCAGTC | - |  |
|  | 17 HCNV L R1 | 13027 | TGTCTAGGCTGTTCCTTTTCG | - |  |
|  | 18 HCNV L F | 12927 | AGCCATCATCTACTATACCAA | + | 979 |
|  | 18 HCNV L R2 | 13887 | TGTTTTCCCCTAGCTACCA | - |  |
|  | 18 HCNV L R1 | 13991 | AAGTAAAGTCCCTGTGCAGA | - |  |
|  | 19 HCNV L F1 | 13705 | TTCTCTAGTAGGGCTAGCATC | + | 878 |
|  | 19 HCNV L F2 | 13740 | CACTCTTATGTCGAATGGTC | + |  |
|  | 19 HCNV L R | 14597 | TTCGTCATCACTGTCATAGGC | - |  |
|  | 20 HCNV L F1 | 14223 | CAAACTTCCACCGAATTTGC | + | 439 |
|  | 20 HCNV L F2 | 14368 | ATTATCCCAAGGAATAGCACAC | + |  |
|  | 20 HCNV L R | 14785 | CCCCTATTTCCAAGAAAACCAT | - |  |
| Segment S | 1 HCNV S F | 1 | AAGACATTCAACCAGTAATCCGAG | + | 913 |
|  | 1 HCNV S R2 | 893 | AGGTCATCCTTCGTGGCAAA | - |  |
|  | 1 HCNV S R1 | 935 | TCGGTCATTTACATCTGGGTA | - |  |
|  | 2 HCNV S F1 | 765 | AGACTGAAACAGCTCCTGAA | + | 984 |
|  | 2 HCNV S F2 | 793 | CCGGTCGTCACAAACAGAA | + |  |
|  | 2 HCNV S R | 1757 | CAGCATTCCATCAGTGCAAG | - |  |
|  | 3 HCNV S F | 1678 | CAACTTCAGGGCTCACCAC | + | 886 |
|  | 3 HCNV S R2 | 2545 | GCCGTCTTTTCACACCTTC | - |  |
|  | 3 HCNV S R1 | 2583 | TGCCTCCCTGAGCTCTCTG | - |  |
|  | 4 HCNV S F | 2451 | TCCCCACACCAGGTCCATC | + | 902 |
|  | 4 HCNV S R2 | 3333 | AACAATAACAAGTCAGTGGC | - |  |
|  | 4 HCNV S R1 | 3407 | ATTACAAAGTACACGACACAGG | - |  |
|  | 5 HCNV S F | 3097 | TTTTCCTCAGCACAATGGAG | + | 633 |
|  | 5 HCNV S R2 | 3709 | TTCATAAGTGCAGCCTAGTCT | - |  |
|  | 5 HCNV S R1 | 3745 | TTCCTTCCTAGCTGCGTGA | - |  |
|  | 6 HCNV S F1 | 3508 | CCTGCTCAACTGTTATGGT | + | 396 |
|  | 6 HCNV S F2 | 3532 | CCTTAACTGTATAGGTGGCATT | + |  |
|  | 6 HCNV S R | 3907 | TCATTCCACAAAAAGCGGGTA | - |  |

| **Supplementary Table 10.** GenBank accession number (nucleotide) for phylogenetic analysis. | | | | | | | |
| --- | --- | --- | --- | --- | --- | --- | --- |
| **Family** | **Genus** | **Virus** | **Strain** | **Accession No.** | | | |
|  |  |  |  | **L** | **M** | | **S** |
| *Nairoviridae* | *Orthonairovirus* | Nairobi sheep disease virus | SDWL08/2020/China/Shandong | OR495605 | OR495606 | | OR495604 |
| *Nairoviridae* | *Orthonairovirus* | Nairobi sheep disease virus | 779 | HM991306 | HQ286599 | | HQ286602 |
| *Nairoviridae* | *Orthonairovirus* | Nairobi sheep disease virus | Jilin | NC034387 | KM464725 | | KM464724 |
| *Nairoviridae* | *Orthonairovirus* | Nairobi sheep disease virus | NSDV/SZWH2 | MW721870 | MW721871 | | MW721872 |
| *Nairoviridae* | *Orthonairovirus* | Nairobi sheep disease virus | NL/NS/NM | OQ716535 | OQ716534 | | OQ716536 |
| *Nairoviridae* | *Orthonairovirus* | Nairobi sheep disease virus | nlb-dd-tick | OQ581155 | OQ581154 | | OQ581153 |
| *Nairoviridae* | *Orthonairovirus* | Orthonairovirus haemorrhagiae | 813042UAE | MF289417 | MF289418 | | MF289419 |
| *Nairoviridae* | *Orthonairovirus* | Orthonairovirus haemorrhagiae | YL16070 | KY354082 | KY354081 | | KY354080 |
| *Nairoviridae* | *Orthonairovirus* | Orthonairovirus haemorrhagiae | HANM-18 | MN832723 | MN832722 | | MN832721 |
| *Nairoviridae* | *Orthonairovirus* | Hughes orthonairovirus |  | MK896563 | MK896562 | | MK896561 |
| *Nairoviridae* | *Orthonairovirus* | Estero Real virus | K329 | MH017280 | MH017286 | | MH017274 |
| *Nairoviridae* | *Orthonairovirus* | Qalyub virus | ErAg370 | NC034511 | KU343161 | | KU343162 |
| *Nairoviridae* | *Orthonairovirus* | Clo Mor virus | ScotAr7 | NC034561 | KU343140 | | KU343141 |
| *Nairoviridae* | *Orthonairovirus* | Tillamook virus | RML86 | KU925494 | KU925495 | | KU925496 |
| *Nairoviridae* | *Orthonairovirus* | Taggert virus | OTU36.IU7 | MN830228 |  | |  |
| *Nairoviridae* | *Orthonairovirus* | Orthonairovirus dugbeense | 15AC-T25 | LC579816 | LC579817 | | LC579818 |
| *Nairoviridae* | *Orthonairovirus* | Kupe virus | K611 | EU257628 | NC078128 | | NC 078129 |
| *Nairoviridae* | *Orthonairovirus* | Hazara virus | JC280 | NC038709 | KP406724 | | KP406725 |
| *Nairoviridae* | *Orthonairovirus* | Meihua Mountain virus | LJ01 | ON184084 | ON184085 | | ON184086 |
| *Nairoviridae* | *Orthonairovirus* | Tofla virus | Nag-Hfor-2014 | LC030115 | LC030114 | | LC030113 |
| *Nairoviridae* | *Orthonairovirus* | Tofla virus |  | NC029124 | NC029123 | | NC 029122 |
| *Nairoviridae* | *Orthonairovirus* | Wetland virus | PS1 | OR860402 | OR860408 | | OR860396 |
| *Nairoviridae* | *Orthonairovirus* | Wetland virus | T3G2 | OR860403 | OR860411 | | OR860400 |
| *Nairoviridae* | *Orthonairovirus* | Dadong tick virus |  |  |  | |  |
| *Nairoviridae* | *Orthonairovirus* | Shanxi tick virus 2 | CLCM-037 | OR114977 | OR114978 | | OR114976 |
| *Nairoviridae* | *Orthonairovirus* | Shanxi tick virus 2 | 21HBT4 | PP471909 | PP471908 | | PP471907 |
| *Nairoviridae* | *Orthonairovirus* | Shanxi tick virus 2 | TIGMIC20 | ON811936 |  | |  |
| *Nairoviridae* | *Orthonairovirus* | Shanxi tick virus 2 | SXO338nairoV | MZ244235 | MZ244236 | | MZ244237 |
| *Nairoviridae* | *Orthonairovirus* | Yanbian Nairo tick virus 1 | TIGMIC1 | ON746415 |  | |  |
| *Nairoviridae* | *Orthonairovirus* | Jian nariovirus | NE-JA | ON408088 | ON408089 | | ON408090 |
| *Nairoviridae* | *Orthonairovirus* | Jian nariovirus | NE-YC2 | ON408085 | ON408086 | | ON408087 |
| *Nairoviridae* | *Orthonairovirus* | Yanbian Nairo tick virus 2 | TIGMIC1 | ON746416 |  | |  |
| *Nairoviridae* | *Orthonairovirus* | Antu virus | YBtick202124 | OQ207703 | OQ207702 | | OQ207701 |
| *Nairoviridae* | *Orthonairovirus* | Songling virus | NE-TH2 | ON408079 | ON408080 | | ON408081 |
| *Nairoviridae* | *Orthonairovirus* | Songling virus | YC585 | MT328779 | MT328778 | | MT328780 |
| *Nairoviridae* | *Orthonairovirus* | Songling virus | NE-TH1 | ON408076 | ON408077 | | ON408078 |
| *Nairoviridae* | *Orthonairovirus* | Songling virus | HLJ1202 | NC079001 | MT328775 | | MT328777 |
| *Nairoviridae* | *Orthonairovirus* | Tacheng Tick Virus 1 | TC253 | NC031284 | KM817717 | | KM817743 |
| *Nairoviridae* | *Orthonairovirus* | Tacheng Tick Virus 1 | QH1 | MK554672 | MK554676 | | MK554680 |
| *Nairoviridae* | *Orthonairovirus* | Peribunyaviridae sp. | 21HBT6 | PP471896 | PP471895 | | PP471894 |
| *Nairoviridae* | *Orthonairovirus* | Henan tick virus | CLCM-083 | OR114972 | OR114985 | | OR114971 |
| *Nairoviridae* | *Orthonairovirus* | Henan tick virus | SZSX | OR573901 | OR573900 | | OR573899 |
| *Nairoviridae* | *Orthonairovirus* | Huangpi Tick Virus 1 | 21HBT2 | PP471901 | PP471900 | | PP471899 |
| *Nairoviridae* | *Orthonairovirus* | Huangpi Tick Virus 1 | H124-1 | NC031135 | KM817706 | | KM817734 |
| *Nairoviridae* | *Orthonairovirus* | Pangolin orthonairovirus | BIME1 | ON024087 | ON024088 | | ON024089 |
| *Nairoviridae* | *Orthonairovirus* | Wenzhou Tick Virus | TS1-2 | NC031291 | KM817718 | | KM817745 |
| *Nairoviridae* | *Orthonairovirus* | Tamdy virus | YL16083 | MT815992 | MT815994 | | MT815994 |
| *Nairoviridae* | *Orthonairovirus* | Yezo virus | T-JL01 | ON563280 | ON563281 | | ON563282 |
| *Nairoviridae* | *Orthonairovirus* | Yezo virus | YBQG1739 | OR148889 | OR148887 | | OR148888 |
| *Nairoviridae* | *Orthonairovirus* | Yezo virus | T-IM01 | ON563277 | ON563278 | | ON563279 |
| *Nairoviridae* | *Orthonairovirus* | Yezo virus | HH003-2020 | LC621355 | LC621356 | | LC621357 |
| *Nairoviridae* | *Orthonairovirus* | Yezo virus | HH001-2019 | LC621352 | LC621353 | | LC621354 |
| *Nairoviridae* | *Orthonairovirus* | Yezo virus | T-HLJ02 | ON563271 | ON563272 | | ON563273 |
| *Nairoviridae* | *Orthonairovirus* | Yezo virus | H-IM01 | ON563283 | ON563284 | | ON563285 |
| *Nairoviridae* | *Orthonairovirus* | Yezo virus | T-HLJ01 | ON563268 | ON563269 | | ON563270 |
| *Nairoviridae* | *Orthonairovirus* | Sulina virus | b30 | PP260003 | PP260005 | | PP260004 |
| *Nairoviridae* | *Orthonairovirus* | Sulina virus | IxriSL16-03 | MT263288 | MT263289 | | MT263290 |
| *Nairoviridae* | *Orthonairovirus* | Sulina virus | IxriSL16-01 | NC078999 | MT263283 | | MT263284 |
| *Nairoviridae* | *Orthonairovirus* | Yezo virus | TIGMIC2 | ON811841 | PQ231074 | | PQ231069 |
| *Nairoviridae* | *Orthonairovirus* | Yezo virus | TIGMIC1 | ON811839 | PQ231073 | | PQ231068 |
| *Nairoviridae* | *Norwavirus* | Beiji nairovirus | UTR-4158 | LC795515 |  | | LC795514 |
| *Nairoviridae* | *Norwavirus* | Beiji nairovirus | HL8 | PP473590 |  | | PP473602 |
| *Nairoviridae* | *Norwavirus* | Beiji nariovirus | NE-TH3 | ON408098 |  | | ON408099 |
| *Nairoviridae* | *Norwavirus* | Beiji nairovirus | YKS44 | MN122079 |  | | MN122080 |
| *Nairoviridae* | *Norwavirus* | Beiji nairovirus | BSQG1734 | OR148703 |  | | OR148704 |
| *Nairoviridae* | *Norwavirus* | Beiji nariovirus | NE-TH4 | ON408100 |  | | ON408101 |
| *Nairoviridae* | *Norwavirus* | Beiji nariovirus | NE-DH3 | ON408106 |  | | ON408107 |
| *Nairoviridae* | *Norwavirus* | Beiji nairovirus | H39 | MW315103 |  | | MW315109 |
| *Nairoviridae* | *Norwavirus* | Norway nairovirus 1 | NOR/B1V/Tofte/2014 | MF141048 |  | | MF141049 |
| *Nairoviridae* | *Norwavirus* | Grotenhout virus | Gierle-1 | NC078265 |  | | KY700683 |
| *Nairoviridae* | *Norwavirus* | Mikuni nairovirus | FKS-1108 | LC795519 |  | | LC795518 |
| *Nairoviridae* | *Norwavirus* | Pustyn virus | Rus/Ixricinus/Moscow/2018 | MN542360 |  | | MN542361 |
| *Nairoviridae* | *Norwavirus* | Gakugsa tick virus | Rus/Ixpersulcatus/Karelia/1/2018 | MN542362 |  | | MN542363 |
| *Nairoviridae* | *Norwavirus* | Norwavirus sp. | p74 | OQ269595 |  | | OQ269596 |
| *Nairoviridae* | *Norwavirus* | Norwavirus sp. | p51 | OQ269601 |  | | OQ269602 |
| *Nairoviridae* | *Norwavirus* | Pustyn virus | IxRic1 | KT007142 |  | | KT007143 |
| *Nairoviridae* | *Norwavirus* | Yichun nariovirus | NE-FZ3 | ON408112 |  | | ON408113 |
| *Nairoviridae* | *Norwavirus* | Yichun nariovirus | NE-YC3 | ON408110 |  | | ON408111 |
| *Nairoviridae* | *Norwavirus* | South Bay virus | SC1 | KX184198 |  | | KX184199 |
| *Large segment of the viral genome (L); Medium segment of the viral genome (M); Small segment of the viral genome (S). | | | | | |  |  |
